# Constraints and tunability of antigen-agnostic memory durability

**DOI:** 10.1101/2025.07.18.663399

**Authors:** Shubham Tripathi, Can Liu, William W. Lau, Rachel Sparks, John S. Tsang

## Abstract

Apart from antigen-specific immune memory, infection or vaccination can also induce antigen-agnostic memory including bystander T cell activation memory and trained innate immunity. Determinants of the durability of such memory remain unclear. We developed mathematical models to show that antigen-agnostic memory durability is constrained by immune cell turnover and cytokine dependence. Trained immunity durability is further constrained by the fidelity of epigenetic state transmission during self-renewal and differentiation. Using computer simulations and a machine learning-based parameter-phenotype mapping approach, we find that positive feedback mediated by immune cell-secreted cytokines, a motif seen across immune circuits, can lead to tunable memory involving excitable responses to acute challenges followed by slow relaxation to the pre-challenge baseline. We propose specific experiments to test predictions from our models. Our findings thus reveal a cell extrinsic mechanism that can overcome the constraints imposed by cell turnover and epigenetic state transmission to drive long-lasting, tunable antigen-agnostic memory.

## INTRODUCTION

Acute response to inflammatory challenges such as infection or vaccination includes both antigen-agnostic and antigen-specific components. The antigen-agnostic response is triggered upon the recognition of pathogen-associated and damage-associated molecular patterns.^1,2^ These early responding cells then signal to other immune cells via inflammatory cytokines to orchestrate downstream responses^3^, including the activation and expansion of antigen-specific naïve T and B cells, culminating in the generation of memory T and B cells. Such memory T and B cells have long been recognized as the basis of immunological memory since they can be rapidly activated and expanded upon subsequent encounter with the same or related antigens.^4^ Thus, long-term immune memory has traditionally been deemed as being mainly antigen-specific.^5^ However, recent studies have documented long-term molecular and cellular changes in antigen non-specific myeloid and lymphoid cells, including in their counts, gene expression, and epigenetic states, after inflammatory challenges in both mice^6–10^ and humans.^11–14^ Such changes potentially represent different forms of antigen-agnostic memory since they can last beyond the acute response phase and have been implicated in altering the immune responses to subsequent antigenically unrelated challenges. However, the kinetics and durability of such memory, and how they arise from the molecular and cellular interactions that underly the antigen-agnostic response, are not well understood.

One component of the antigen-agnostic response involves the T cell receptor (TCR)-independent activation and expansion of CD4+ and CD8+ memory T cells, termed “bystander activation”.^15–17^ In response to inflammatory cytokines such as IL-15 and IL-18 upregulated by both immune and non-immune cells in response to inflammatory challenges, antigen-non-specific CD8+ memory T cells can expand, secrete IFN-*γ*, and express cytotoxic molecules including perforin and granzyme B.^15^ In the case of CD4+ memory T cells, the bystander activation response can be triggered by cytokines such as IL-1*β*, IL-12, and IL-18; these cells then secrete cytokines in a cell subset-dependent manner: IFN-*γ* in the case of Th1, IL-5 and IL-13 in the case of Th2, and IL-17 and GM-CSF in the case of Th17 memory T cells.^16^ In mouse studies, bystander activation of memory T cells has been shown to drive a protective immune response upon exposure to previously unseen pathogens (see Table 1 in Paprckova *et al*.^17^), primarily involving IFN-*γ* production and innate-like cytotoxicity via the NKG2D receptor. In humans, IL-15-driven bystander activation of CD8+ memory T cells that lack specificity for the hepatitis A virus has been reported in patients with acute hepatitis A.^18^ IL-15 can also activate and expand CD8+ memory T cells independent of their antigen specificity in human immunodeficiency virus (HIV)-infected individuals.^19^ Exposure to dengue antigens^20^ and booster injection with tetanus toxoid^21^ have both been shown to drive bystander CD4+ memory T cell activation. A recent study from our group comparing the immune profiles of clinically healthy individuals who had recovered from mild COVID-19 with age- and sex-matched controls (*i.e*., individuals with who no history of COVID-19) showed that the counts of bystander activated, non-SARS-CoV-2-specific CD8+ memory T cells can be elevated for months in the blood after clinical recovery from SARS-CoV-2 infection, particularly in males.^11^ Elevated counts of these cells were shown to predict a higher response to subsequent influenza vaccination, supporting the notion that bystander activation of CD8+ memory T cells during the acute response can lead to the development of antigen-agnostic immune memory that is encoded by the counts of these cells and can last for multiple months.

**Table 1.**
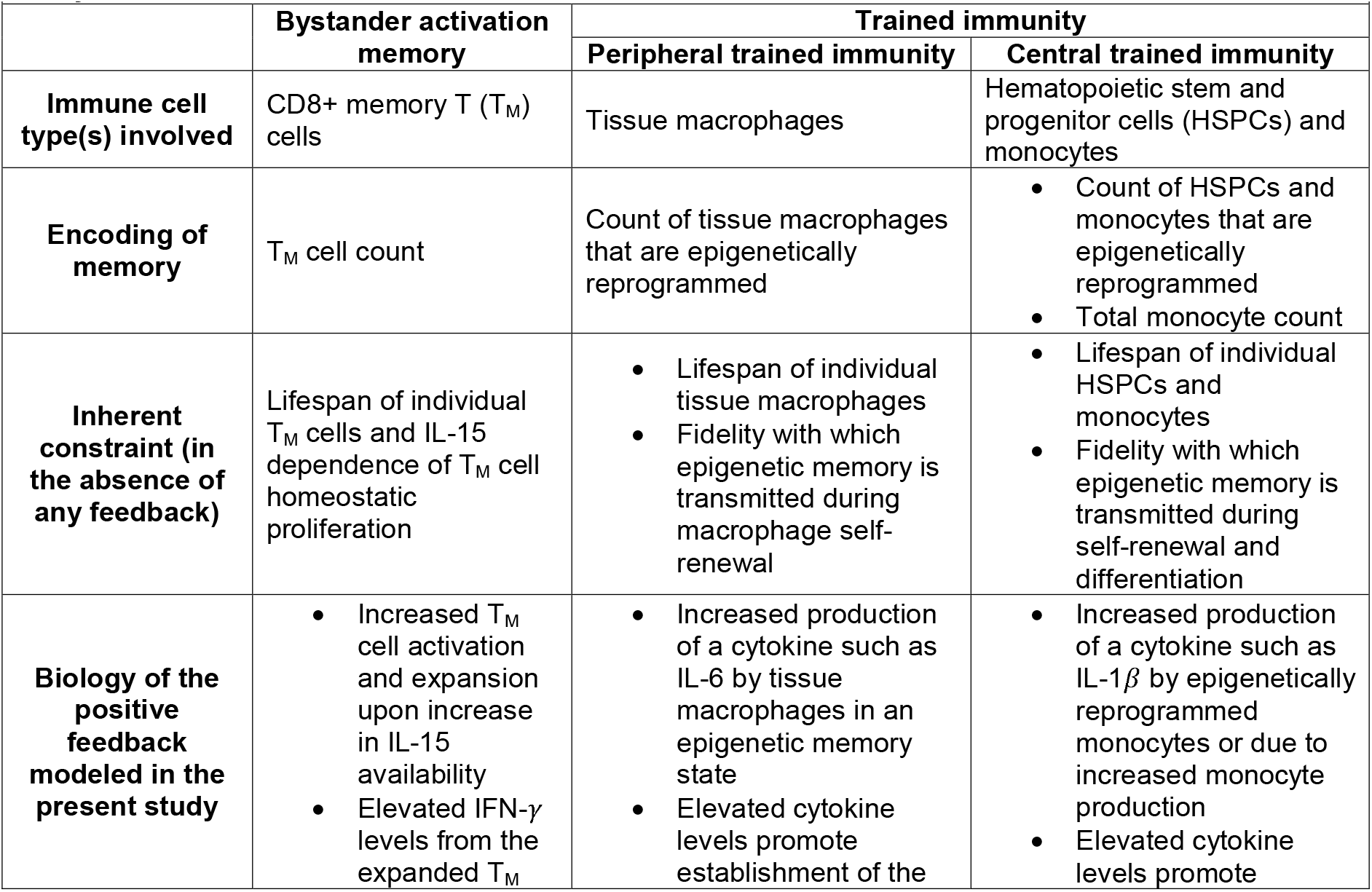
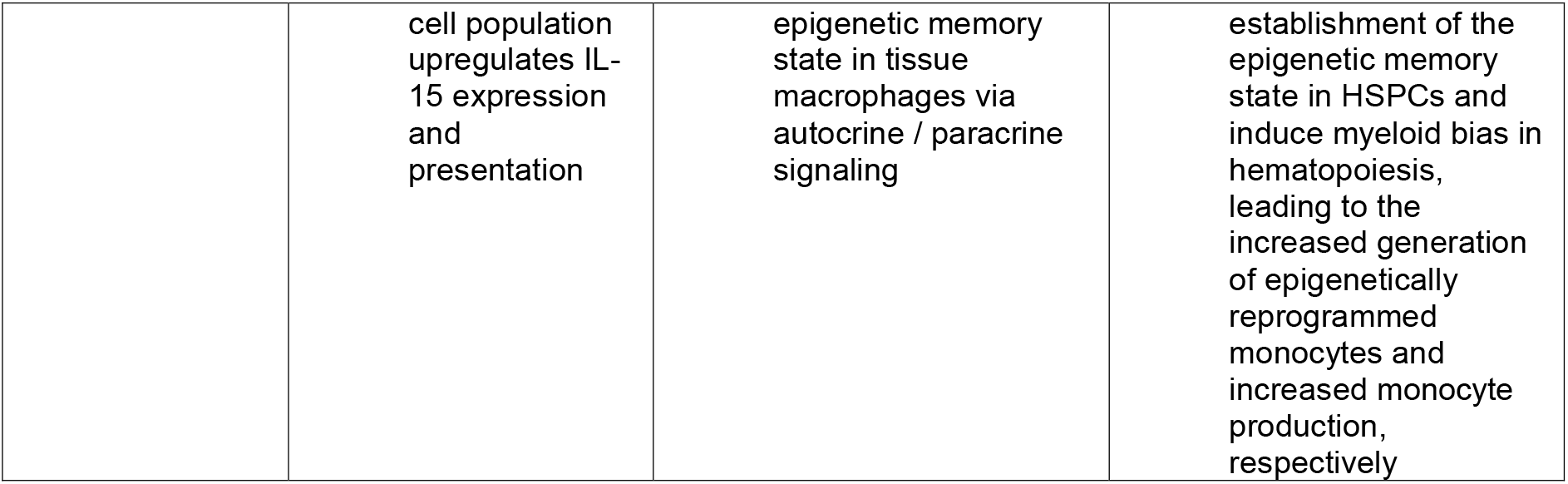
An overview of the types of antigen-agnostic immune memory as modeled in the present study.

|  | **Bystander activation memory** | **Trained immunity** |  |
| --- | --- | --- | --- |
|  |  | **Peripheral trained immunity** | **Central trained immunity** |
| **Immune cell type(s) involved** | CD8+ memory T ($T_M$) cells | Tissue macrophages | Hematopoietic stem and progenitor cells (HSPCs) and monocytes |
| **Encoding of memory** | $T_M$ cell count | Count of tissue macrophages that are epigenetically reprogrammed | Count of HSPCs and monocytes that are epigenetically reprogrammed<br>Total monocyte count |
| **Inherent constraint (in the absence of any feedback)** | Lifespan of individual $T_M$ cells and IL-15 dependence of $T_M$ cell homeostatic proliferation | Lifespan of individual tissue macrophages<br>Fidelity with which epigenetic memory is transmitted during macrophage self-renewal | Lifespan of individual HSPCs and monocytes<br>Fidelity with which epigenetic memory is transmitted during self-renewal and differentiation |
| **Biology of the positive feedback modeled in the present study** | Increased $T_M$ cell activation and expansion upon increase in IL-15 availability<br>Elevated IFN-γ levels from the expanded $T_M$ | Increased production of a cytokine such as IL-6 by tissue macrophages in an epigenetic memory state<br>Elevated cytokine levels promote establishment of the | Increased production of a cytokine such as IL-1β by epigenetically reprogrammed monocytes or due to increased monocyte production<br>Elevated cytokine levels promote |
|  | cell population upregulates IL-15 expression and presentation | epigenetic memory state in tissue macrophages via autocrine / paracrine signaling | establishment of the epigenetic memory state in HSPCs and induce myeloid bias in hematopoiesis, leading to the increased generation of epigenetically reprogrammed monocytes and increased monocyte production, respectively |

Another key component of the antigen-agnostic immune response involves the epigenetic reprogramming of immune cells. This phenomenon has been most studied in myeloid cells including monocytes and macrophages, although emerging evidence suggests that other immune cell types, such as bystander activated memory T cells^11^ and memory B cells,^22^ can also undergo epigenetic reprogramming in response to inflammatory challenges. Such reprogramming can involve chromatin state changes at genomic loci associated with inflammatory response genes, potentially facilitating more rapid production of inflammatory cytokines in response to a subsequent challenge. Multiple studies have shown that effector immune cells with such reprogrammed epigenetic states can be maintained even after the initial inflammatory challenge has resolved, creating innate immune memory this is often termed “trained immunity”.^23^ In mouse models of ligature-induced periodontitis^8^ and ischemic stroke,^9^ the reprogrammed epigenetic state in monocytes could be detected for at least a month after the initial challenge. Similarly, epigenetic memory of infection with a mouse-adapted strain of SARS-CoV-2 could be detected in murine alveolar macrophages more than a month after infection clearance, and these epigenetically reprogrammed cells were then shown to ameliorate the disease severity of the subsequent influenza infection.^10^ Bacillus Calmette-Guérin (BCG) vaccination can also epigenetically reprogram hematopoietic progenitors in both mice^7^ and humans,^12,14^ leading to increased counts of inflammatory monocytes and bone marrow-derived macrophages. Such epigenetic reprogramming of hematopoietic progenitors following an inflammatory challenge has been posited to be a key driver of long-lasting trained immunity since these reprogrammed self-renewing progenitors can generate reprogrammed effectors such as monocytes over the long-term; myeloid effectors are short-lived and non-renewing, and reprogrammed cells would otherwise be lost quickly after the end of the acute response.^23^

While the role of antigen-agnostic memory in shaping immune responses has increasingly come into focus, key questions remain unanswered. What determines the durability of such memory? How is the memory, as encoded by the cell counts and reprogrammed epigenetic states, maintained and how fast does it decay? It is often assumed that the immune cell populations expanded and / or epigenetically altered in response to an inflammatory challenge can be maintained for a period of time beyond the acute phase of the response to the challenge. However, given that immune cell population sizes and epigenetic states can undergo changes as cells undergo self-renewal, differentiation, and death, conditions under which antigen-agnostic immune memory can be maintained long-term are unclear. Answering these questions can reveal how inflammatory exposures shape the immune state of an individual at both short and long timescales, and will thus lead to a better understanding of how baseline immune setpoints evolve as a function of successive exposures and how these setpoints impact the response to new inflammatory challenges.^24^

Here, we address the durability question by using mathematical models of immune cell dynamics, together with a machine learning-based approach, to quantitatively explore the durability of both bystander activation memory and trained immunity (see Table 1 for an overview). We show that bystander activation memory durability is inherently constrained by the lifespan of CD8+ memory T cells and by the limited availability of self-renewal-driving cytokines such as IL-15. Trained immunity durability is similarly constrained by the lifespan of myeloid effectors. It is constrained further by the expected loss of epigenetic information during self-renewal and differentiation. Building on supporting evidence in the literature that cytokine-mediated positive feedback is present in immune cell circuits that underly both bystander activation memory and trained immunity, we show that such feedback can overcome the aforementioned constraints and lead to an excitable acute response followed by antigen-agnostic memory with tunable durability. Our theoretical analysis thus reveals that despite the distinct molecular and cellular players involved in bystander activation memory and trained immunity, positive feedback is a shared circuit motif that can drive tunable memory durability.

## RESULTS

### Durability of bystander activation memory is constrained by memory T cell half-life and the IL-15-dependence of their maintenance

We first developed a mathematical model of IL-15 signaling-driven, TCR-independent proliferation of memory T cells (SI Section I). CD8+ memory T cells (hereafter referred to as T_M_ cells) are relatively short-lived, with an estimated half-life of around 14 days in mice^25^ and around 46 days in humans.^26^ T cell memory is maintained long-term via the homeostatic proliferation of these cells primarily in response to IL-15 stimulation from both non-immune and immune cell types.^27–30^ Competition for IL-15, whose availability is limited, constrains the size of the T_M_ cell pool at steady state (Figures 1A and S1A). We modeled this behavior using a system of ordinary differential equations wherein T_M_ cells die at a fixed rate while their proliferation rate is dependent on the IL-15 stimulation received (Equation S1-S3 and Figure S1A). IL-15 expression is known to be upregulated acutely in response to bacterial and viral infections.^27^ IL-15 upregulation in response to inflammatory challenges increases the amount of proliferation signaling available (Figure S1A), thereby increasing T_M_ cell proliferation in a TCR-independent manner; additionally, T_M_ cells secrete IFN-*γ* in response to IL-15 stimulation.^11,15^ This TCR signaling-independent increase in T_M_ cell counts and secreted IFN-*γ* levels in response to inflammatory challenges is termed bystander activation (Figure S1B, shaded region). Once the inflammatory challenge has resolved and IL-15 availability returned to its pre-infection, homeostatic level, the T_M_ cell count declines since the homeostatic IL-15 level is insufficient to maintain the T_M_ cell pool expanded in response to the acute challenge (Figure S1A-B). The rate of cell count decline is determined by the half-life of T_M_ cells, which is thus a key determinant of the duration after inflammation resolution for which T_M_ cell counts are above the pre-inflammatory challenge baseline. If a subsequent challenge is encountered while T_M_ cell counts are still elevated, the bystander T_M_ cell activation-driven IFN-*γ* response will be stronger as compared to the scenario without any prior inflammatory exposure (Figure S1C). This effect is termed bystander activation memory. Such memory is encoded by T_M_ cell counts and its durability is therefore constrained by the half-life of T_M_ cells (Figure 1B).

**Figure 1.**
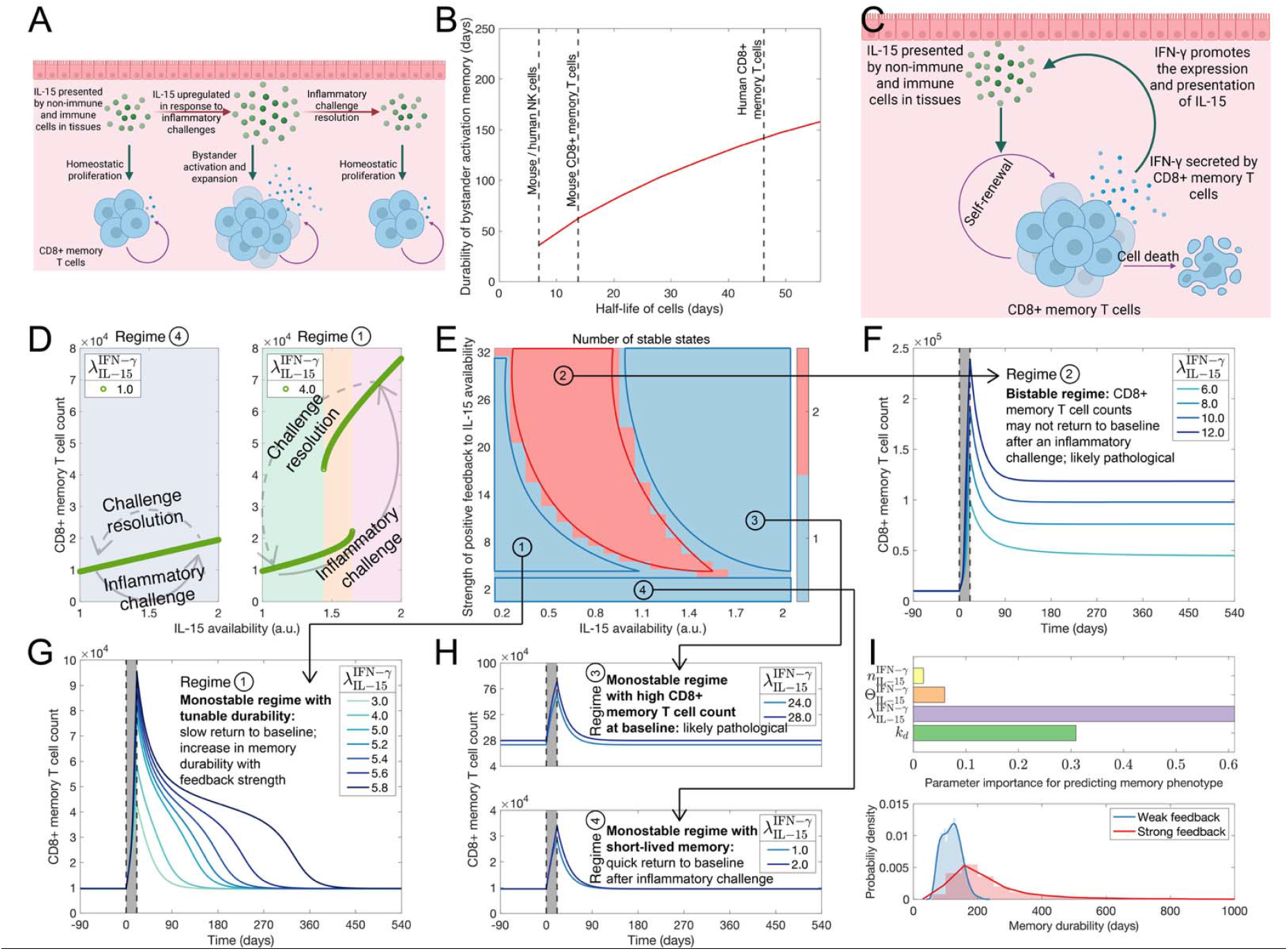
Constraints and tunability of bystander activation memory durability. (A) Schematic showing bystander activation of CD8+ memory T (T_M_) cells in response to an inflammatory challenge. (B) Durability of bystander activation memory as a function of the half-life of T_M_ cells (and in the absence of feedback from T_M_ cell-secreted IFN-*γ* to IL-15 availability). Half-lives for some mouse and human cell types are indicated by dashed vertical lines. See also Figures S1B-C. (C) Schematic showing the positive feedback loop involving IL-15 signaling-driven T_M_ cell activation and expansion, secretion of IFN-*γ* by T_M_ cells, and upregulation of IL-15 expression and presentation in response to IFN-*γ* signaling. (D) T_M_ cell count as a function of IL-15 availability in regime 4 (left panel; very weak feedback from T_M_ cell-secreted IFN-*γ* to IL-15 availability) and regime 1 (right panel; strong positive feedback). Arrows indicate how the IL-15 availability and T_M_ cell count may change in response to an inflammatory challenge (bottom arrow) and upon challenge resolution (top arrow). The different regimes are shown in E. (E) Phase diagram showing the number of possible stable states for T_M_ cell dynamics as a function of the IL-15 availability (X axis) and the strength of positive feedback from T_M_ cell-secreted IFN-*γ* to IL-15 availability (Y axis). See also Figures S1D-G. (F-H) Kinetics of T_M_ population contraction post-inflammatory challenge resolution in different regimes (F: regime 2, G: regime 1, H: regimes 3 and 4). 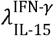 indicates the positive feedback strength with higher values corresponding to stronger feedback. In each panel, the region shaded in grey indicates the duration of the acute inflammatory challenge; there is no challenge outside of those regions. See also Figures S1G-I. (I) Top: relative importance or contribution of different model parameters to predicting the memory durability phenotype using a random forest model trained on simulated data. Bottom: distribution of bystander activation memory durability values for weak feedback 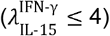 and strong feedback 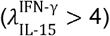 cases. See also Figure S1J-L. Panels A and D were created using Biorender.com. Additional details concerning how each panel was generated are included in SI Section IC.

Note that, here, we have analyzed the memory T cell response to antigens that have not been encountered previously, *i.e*., in the scenario wherein, prior to the inflammatory challenge, there are no T_M_ cells in the memory pool specific to the antigen involved in the challenge. While such a challenge would be accompanied by TCR-dependent, antigen-specific naïve T cell responses in parallel, IL-15 upregulation by myeloid cells as well as non-immune cells (such as tissue epithelial cells) drives the TCR-independent bystander memory T cell response modeled here. This simpler case has been the primary focus of experimental studies on bystander activation memory.^17^ In the case of challenge with a previously encountered antigen, IL-15-driven bystander T_M_ cell activation will be accompanied by the activation and expansion of antigen-specific T_M_ clones. Although such a scenario can be accommodated by our mathematical setup, we focus on the simpler case of first-time antigen encounter here to identify inherent constraints on bystander activation memory durability.

### IFN-*γ* signaling-mediated positive feedback can tune bystander activation memory durability

Till now, we have considered how IL-15 availability determines T_M_ cell dynamics at homeostasis and in response to inflammatory challenges. Note that T_M_ cells can, in turn, alter IL-15 availability by secreting IFN-*γ* which is known to upregulate IL-15 expression and presentation by both immune and non-immune cell types.^31,32^ This creates a positive feedback loop: IL-15 promotes the activation (including IFN-*γ* expression and secretion) and expansion of T_M_ cells while the IFN-*γ* secreted by T_M_ cells promotes IL-15 expression and presentation (Figure 1C). Incorporating this behavior into our mathematical model (Equation S4), we find rich T_M_ cell dynamics that can operate in four different behavioral regimes depending on (a) the IL-15 availability at baseline and (b) the strength of IFN-*γ*-mediated feedback (model parameter 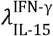), defined as the maximum fold-change in IL-15 availability that IFN-*γ* can cause (Figures 1D-E and S1D-G).

Both in regime 1, with strong feedback from TM cell-secreted IFN-*γ* to IL-15, and in regime 4, with weak feedback, there is only one stable state (*i.e*., monostable regimes). The stable states in both regimes are characterized by low, homeostatic TM cell counts and IFN-*γ* levels. However, the acute response to inflammatory challenges as well as the tunability of memory durability are regime dependent. When starting from a regime 1 homeostatic state, the acute response is switch-like: jump to a state with multi-fold higher T_M_ cell counts and IFN-*γ* levels (*i.e*., turning the switch “on”). This switch-like, excitable acute response arises as the dynamics transition from monostable regime 1 to bistable regime 2 and then to monostable regime 3 in response to IL-15 upregulation (bottom arrow in Figure 1D, right panel). Upon inflammation resolution, both T_M_ cell counts and IFN-*γ* levels return to homeostatic levels (turning the switch “off”) as the system returns from regime 3 to regime 1 via regime 2 (top arrow in Figure 1D, right panel). Note that the return path is different from the one taken during the acute response. Such behavior, termed hysteresis, is characteristic of biological systems involving positive feedback.^33^ Importantly, return to regime 1 is slower and can be tuned by the strength of the feedback; in such a scenario, T_M_ cell counts can remain elevated compared to the homeostatic state long after challenge resolution (Figures 1G and S1H-I). In contrast, when starting from a homeostatic state in regime 4 in response to an inflammatory challenge, the dynamics remain in regime 4 as IL-15 availability increases, and the corresponding T_M_ cell count increase during the acute response phase is proportional to the increase in IL-15 (Figure 1D, left panel). There is no hysteresis in this scenario, and the T_M_ cell counts decline quickly (due to exponential decay of the T_M_ cell population) upon challenge resolution (Figure 1H, top panel).

Regimes 1 and 4 correspond to physiological scenarios since they exhibit stable states with low T_M_ cell counts and IFN-*γ* levels in the absence of inflammatory challenges and permit eventual return to the baseline state after the challenge has resolved. In contrast, regimes 2 and 3 are likely pathological. The only stable state in regime 3 is when T_M_ cell counts, IFN-*γ* levels, and IL-15 availability are simultaneously high even in the absence of an inflammatory challenge (Figures 1H, top panel and S1G), corresponding to a persistent inflammation phenotype. Regime 2, encountered during the dynamic transition between regimes 1 and 3 in response to an inflammatory challenge as discussed above, is bistable: one stable state is characterized by low T_M_ cell counts and the other by elevated T_M_ cell counts and IFN-*γ* levels. In this regime, an inflammatory challenge can lead to a switch from the low T_M_ cell count state to the high T_M_ cell count state and since the latter is also stable, T_M_ cell counts may never return to homeostatic levels even after challenge resolution (Figures 1F and S1G). This behavior may be interpreted as an “infinite” memory phenotype (Figures S1H-I) and could reflect pathological scenarios such as post-acute infection syndromes (*e.g*., long COVID)^34,35^ and infection-triggered autoimmunity.^36^

Altogether, our analysis shows that positive feedback from T_M_ cell-secreted IFN-*γ* to IL-15 can overcome the inherent constraints on bystander activation memory encoded by T_M_ cell counts. In particular, our modeling approach identifies regime 1 which corresponds to a tunable bystander activation memory phenotype: memory durability in this regime varies non-linearly with the feedback strength, increasing rapidly as one nears the transition between regimes 1 and 2 (Figure S1H-I). Figures 1D-H and S1D-I show this behavior for a representative parameter set. To more comprehensively assess the dependence of the durability phenotype on positive feedback strength, we simulated T_M_ cell dynamics and bystander activation for an ensemble of biologically plausible parameter sets (see SI Section IC for how the ensemble was generated). Figure S1J shows that memory durability correlates with the feedback strength across the parameter sets in the ensemble. Following a previously described machine learning-based approach,^37,38^ we next trained a random forest model to predict, given a parameter set, the memory durability phenotype and calculated the relative contribution of each parameter to model prediction (Figure S1K-J). Figure 1I (top panel) shows that the positive feedback strength parameter 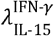 is the top contributor. The T_M_ cell half-life (captured by the parameter *k*_*d*_) makes a smaller yet important contribution, reflecting the durability constraint shown earlier (Figure 1B). Finally, the distribution of memory durability values across these biologically plausible parameter sets differs significantly between the strong versus weak feedback cases. The distribution is both broader (*i.e*., more variable memory durability) and has a longer tail in the strong feedback case, confirming across parameter sets that the inclusion of positive feedback confers tunable memory that could last over a timescale of years (Figure 1I, bottom panel). Note that the long-tail in the strong feedback regime arises from the non-linear dependence of the durability on feedback strength in regime 1 (Figures S1H-J).

### Constraints on the durability of trained immunity

Trained innate immunity posits that inflammatory challenges can cause epigenetic reprogramming of effector myeloid cells such as macrophages and monocytes to an altered state such as one with activating histone modifications and open chromatin at specific genomic loci, including loci associated with inflammatory response genes. Cells in such altered epigenetic states are poised to give rise to an altered immune response upon encountering a subsequent inflammatory challenge.^23^ An epigenetic state encoding the memory of a prior inflammatory exposure is hereafter referred to as an “epigenetic memory state”. Trained immunity can involve cells in the periphery (“peripheral trained immunity”), *e.g*., circulating monocytes and tissue macrophages, as well as hematopoietic progenitors in the bone marrow (“central trained immunity”).^23,40^ To identify the parameters that determine the durability of peripheral and central trained immunity, we employed the same mathematical modeling approach as above, focusing now on epigenetic cell state dynamics in tissue macrophages and in the hematopoietic hierarchy (Figures 2A and 2C, and SI Sections II and III).

**Figure 2.**
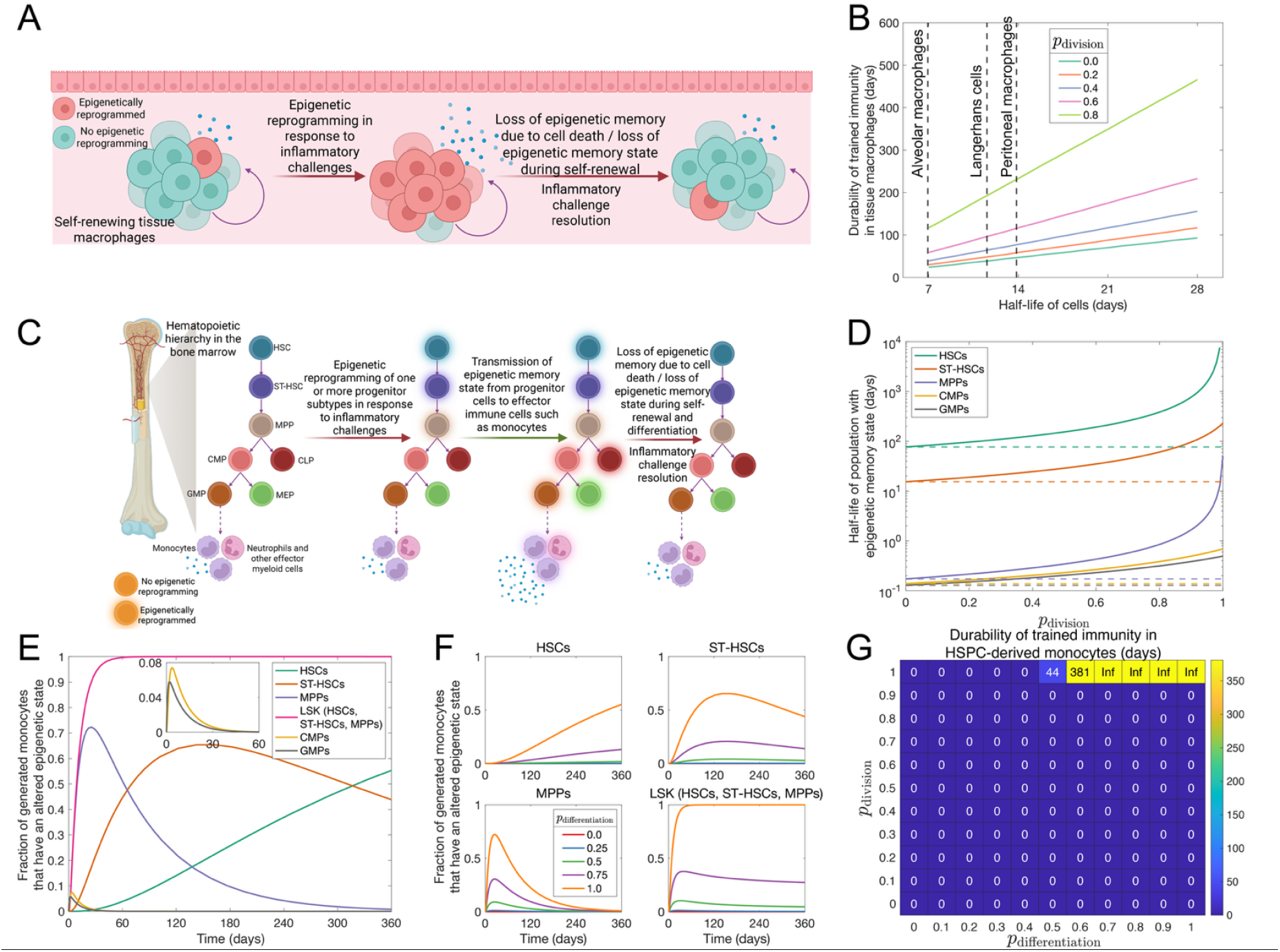
Constraints on the durability of trained immunity. (A) Schematic showing the dynamics of the epigenetic memory state in self-renewing tissue macrophages in response to an inflammatory challenge. (B) Durability of trained immunity as encoded by epigenetically reprogrammed tissue macrophages, shown as a function of the macrophage half-life and for different values of the probability that epigenetic memory state is preserved during cell division (*p*_division_). Estimated half-lives of some mouse tissue macrophage subtypes are indicated by dashed vertical lines. See also Figure S2A-B. (C) Schematic showing the dynamics of the epigenetic memory state during hematopoiesis in the bone marrow in response to an inflammatory challenge. We show the case wherein the inflammatory challenge induces epigenetic reprogramming of HSCs, ST-HSCs, and MPPs (Lin-cKit+Sca1+, or LSK cells in mice), causing these cell subtypes to acquire an epigenetic memory state. The epigenetic memory state can then be transmitted from LSK cells to more differentiated cell types downstream in the hematopoietic hierarchy, ultimately driving the generation of epigenetically reprogrammed monocytes. HSC: hematopoietic stem cell, ST-HSC: short-term hematopoietic stem cell, MPP: multipotent progenitor, CMP: common myeloid progenitor, CLP: common lymphoid progenitor, GMP: granulocyte-monocyte progenitor, MEP: megakaryocyte-erythrocyte progenitor. See also Figure S2C. (D) Half-life of the population with epigenetic memory state induced by the inflammatory challenge as a function of *p*_division_, shown for different HSPC subtypes. The horizontal dashed line in each case indicates the half-life of a single cell. (E) Kinetics of generation of epigenetically reprogrammed monocytes when the epigenetic memory state is induced at *t* = 0 in different HSPC subtypes. Here, we have set *p*_division_ = *p*_differentiation_ = 1. (F) Kinetics of generation of epigenetically reprogrammed monocytes for different values of *p*_differentation_ when epigenetic memory is induced at t = 0 in different HSPC subtypes. Here, we set *p*_division_ = 1 and vary *p*_differentation_. = (G) Durability of trained immunity encoded by epigenetically reprogrammed monocytes, shown as a function of *p*_division_ and *p*_differentiation_, when the initial inflammatory challenge epigenetically reprograms LSK cells (HSCs, ST-HSCs, and MPPs). Behavior is shown for fixed self-renewal and differentiation rates estimated in Busch *et al*.^39^ See also Figure S2D. Panels A and C were created using Biorender.com.

#### Durability of peripheral trained immunity

Tissue macrophage populations are maintained over time via homeostatic self-renewal driven by cytokines such as M-CSF, IL-34, and GM-CSF.^41,42^ Similar to the case of T_M_ cells, such continuous cell turnover (Figure 2A and Table S2) imposes a constraint on the durability of trained immunity (Equations S17 and S19): macrophage population with an epigenetic memory state, expanded in response to an inflammatory challenge, will shrink over time due to cell death once the challenge has resolved unless epigenetic information is faithfully transmitted to the daughter cells during macrophage self-renewal (Figure 2A). Defining trained immunity durability as the duration, after inflammatory challenge resolution, for which the fraction of tissue macrophages with an epigenetic memory state is at least 10% higher than the pre-challenge baseline, we find that the durability is dependent on two key parameters (Figures 2B and S2A-B): (a) macrophage half-life and (b) the probability with which the epigenetic memory state is transmitted to the daughter cells (model parameter *p*_division_). Multiple studies have shown that regions of open chromatin, such as those with transcriptionally active histone marks, can be lost during cell division^43–45^, except in scenarios wherein transcription in the genomic region is persistently present.^46^ Thus, *p*_division_ is often lower than 1 (likely significantly lower than 1), corresponding to significant epigenetic information loss over time; this is predicted to constrain peripheral trained immunity durability to short timescales. Given that macrophages in different tissues exhibit different turnover rates (Table S2), trained immunity durability can also differ between tissues (Figure 2B).

#### Durability of central trained immunity

The mathematical model for central trained immunity is more complex since it involves the hematopoietic process that gives rise to myeloid effectors such as monocytes (Figure 2C). Production of monocytes involves a cascade of differentiation events starting with hematopoietic stem cells (HSCs) at the top of the hematopoietic hierarchy that are capable of long-term self-renewal. Thereafter, the differentiation cascade proceeds through short-term HSCs (ST-HSCs), multipotent progenitors (MPPs), and common myeloid progenitors (CMPs) to granulocyte / monocyte progenitors (GMPs) which ultimately give rise to monocytes (Figure 2C). Along this cascade, self-renewal capability is progressively lost and cells become increasingly lineage-restricted.^47^ Inflammatory challenge-induced epigenetic reprogramming of these hematopoietic stem and progenitors cells (HSPCs) in the bone marrow is believed to underlie durable trained immunity in otherwise short-lived, non-self-renewing effector cells such as monocytes and dendritic cells.^23^ This raises two questions: (a) which specific HSPC subtypes are involved in central trained immunity? and (b) what determines the durability of central trained immunity?

Central trained immunity involves the retention of the epigenetic memory state induced by prior inflammatory exposure in hematopoietic progenitors, which thus serve as a “central memory reservoir”. Once epigenetic memory is established in HSPCs, its durability will be constrained by self-renewal as in the case of tissue macrophages: HSPC subpopulations can lose the epigenetic memory state due to error-prone transmission of the epigenetic state during self-renewal. Further loss can be incurred as the epigenetic memory state is transmitted from HSPCs to effectors like monocytes along the differentiation cascade since cell differentiation during hematopoiesis is accompanied by large-scale changes in chromatin architecture,^48^ and the epigenetic memory state can be lost (at least partially) during each differentiation step.

The abovementioned processes were incorporated into an ordinary differential equations-based model (SI Section IIIB, Equation S40-S45) to probe the determinants of the durability of central trained immunity. The model was instantiated with self-renewal and differentiation rates for different mouse HSPC subtypes reported in Busch *et al*.^39^ where a mouse model with inducible genetic labeling of HSCs was used to analyze the kinetics of hematopoiesis under homeostatic conditions (Figure S2C). We first used the mathematical model to assess the extent to which each HSPC subtype can serve as a reservoir of epigenetic memory. Specifically, for each HSPC subtype, we computed the half-life of epigenetic memory of prior inflammatory exposure, defined as the time it takes for the fraction of cells in an epigenetic memory state to decrease by 50%. Our computation incorporated the loss of the epigenetic memory state during self-renewal as dependent on *p*_division_, the probability that the epigenetic memory state is faithfully transmitted to daughter cells (Equation S56-S60). Figure 2D shows that the epigenetic memory half-life is the highest in the case of HSCs and as expected, decreases in more differentiated progenitors and increases with increase in *p*_division_ (*i.e*., with more accurate transmission of the epigenetic memory state during self-renewal). When *p*_division_is not close to 1, CMPs, GMPs, and MPPs exhibit memory half-lives of less than a day. As *p*_division_ approaches 1, the half-life in the case of CMPs and GMPs remains shorter than one day but can be up to two weeks in MPPs, which are higher up in the hematopoietic hierarchy. In the case ST-HSCs, the epigenetic memory half-life can be up to multiple months for high *p*_division_ and decreases to ∼4 weeks for *p*_division_ close to 0.5. Note that four weeks after the initial inflammatory challenge is the typical time point at which trained immunity has been probed in mouse experiments.^8–10^ Finally, the HSC population can retain epigenetic memory over multiple months even for low *p*_division_. Thus, in mice, only HSCs, ST-HSCs, and to some extent, MPPs can serve as relatively long-term (more than a few days) reservoirs of the epigenetic memory state for trained immunity.

We next assessed the kinetics and extent of transmission of the epigenetic memory state to monocytes from different HSPC subtypes. Different HSPC subpopulations contribute to monocyte production at different rates. The contribution is slower towards the top of the hematopoietic hierarchy: while epigenetic memory state in MPPs will be quickly transmitted to monocytes, it could take weeks if epigenetic memory resides in ST-HSCs and months if it resides in HSCs (Figure 2E). This prediction is consistent with studies showing that under homeostatic conditions in mice, production of effector immune cells is largely driven by ST-HSCs and MPPs, with HSCs only contributing over the long run (> 1 year).^39,49^ Next, given that differentiation can be accompanied by the loss of the epigenetic memory state, we modeled the fidelity of epigenetic memory state transmission during differentiation with the model parameter *p*_differentiation_, the probability that an epigenetic memory state is preserved during a differentiation step (Figure 2F). Trained immunity in monocytes will be possible when the epigenetic memory state is (a) retained in an HSPC subtype(s) for a sufficiently long period for the memory state to be transmitted to monocytes and (b) is sufficiently preserved along the differentiation cascade from the HSPC subtype to monocytes. The first condition depends on *p*_division_ (Figures 2D and 2E), the second on *p*_differentiation_ (Figure 2F); both depend on the HSPC subtype that serves as the central reservoir of epigenetic memory. Figures 2G and S2D show, quantitatively, how the durability of trained immunity, encoded by the production of monocytes with an altered epigenetic state, depends on *p*_division_ and *p*_differentiation_ when the initial inflammatory challenge induces an epigenetic memory state in different HSPC subtypes. We find that trained immunity is durable only when *p*_division_ is very close to 1, *p*_differentiation_ is not too low (> 0.5), and the inflammatory challenge induces an epigenetic memory state in one or more HSPC subtypes near the top of the hematopoietic hierarchy, namely HSCs, ST-HSCs, and MPPs (*i.e*., Lin-cKit+ Sca1+, or LSK cells^47^). Our analysis thus identifies constraints on central trained immunity durability from self-renewal and differentiation during hematopoiesis. Note that *p*_division_ and *p*_differentiation_ are likely context-dependent, and their true values in homeostatic and inflammatory challenge contexts remain unknown. Experimental determination of the values of these parameters will provide valuable quantitative insights into the durability of trained immunity.

Given that the hematopoietic hierarchies in mice and humans are structurally similar (see Figure 1 in Seita and Weissman^47^), our conclusion regarding the importance of *p*_division_ and *p*_differentiation_ in determining central trained immunity durability in mice is expected to hold for humans as well. We further note that our approach to model trained immunity disregards heterogeneity in the epigenetic memory state. Different inflammatory triggers may induce different combinations of epigenetic changes in HSPCs and myeloid effectors. In addition, effector cells with distinct epigenetic memory states may drive different future immune response outcomes. A model to capture such heterogeneity will need to keep track of additional epigenetically distinct subpopulations and, possibly, different *p*_division_ and *p*_differentiation_ values for each epigenetic memory state. However, the overall behavior will still be subject to the same constraints as in the simpler case described above.

### Positive feedback can tune the durability of trained immunity

We showed earlier that the inclusion of positive feedback from IFN-*γ* to IL-15 availability in T_M_ cell dynamics can tune bystander activation memory durability, overcoming the inherent constraints from T_M_ cell half-life and IL-15 dependence. This behavior arises not from characteristics specific to IL-15 or IFN-*γ* signaling but is a general feature of biological systems with positive feedback.^50^ Therefore, we hypothesize that potential positive feedback involving epigenetic reprogramming of tissue macrophages and HSPCs could lead to tunable trained immunity durability. Any such feedback must couple the epigenetic memory state to a signal that can help maintain the memory state even when the initial inflammatory challenge is no longer present. We next discuss examples of immune signaling pathways that could mediate such positive feedback and then assess how such feedback can result in tunable trained immunity durability.

#### Potential positive feedback in trained immunity

IL-6 is one potential positive feedback mediator. In the periphery, macrophages secrete IL-6 in response to inflammatory signals including lipopolysaccharides and the mycobacteria lipoprotein Pam3Cys^6^, and can also respond to IL-6 signaling in both autocrine and paracrine manners.^51,52^ IL-6 signaling in macrophages triggers the phosphorylation of STAT3 and its translocation into the nucleus where it can upregulate the production of IL-6 itself as well as establish more open chromatin at the associated genomic loci.^13^ Provided macrophages in an epigenetic memory state secrete more IL-6 even in the absence of inflammatory challenges, the increased IL-6 can signal to the same or neighboring macrophage(s) to maintain the epigenetic memory state, thereby forming a positive feedback loop. Since STAT3 also links IL-6 signaling to other inflammatory response regulator genes such as IFNAR1, IRF1, JUNB, and JAK2 (as documented in the TRRUST database;^53^ also see SI Section IIA), the epigenetic memory state in this scenario can also involve increased chromatin accessibility at genomic loci associated with other signaling players. Given the centrality of IL-6 in the macrophage inflammatory response and its involvement in trained immunity^23^ including in peritoneal^6^ and alveolar macrophages,^54^ we postulate that IL-6 could be a key mediator of positive feedback in peripheral trained immunity (Figure 3A).

**Figure 3.**
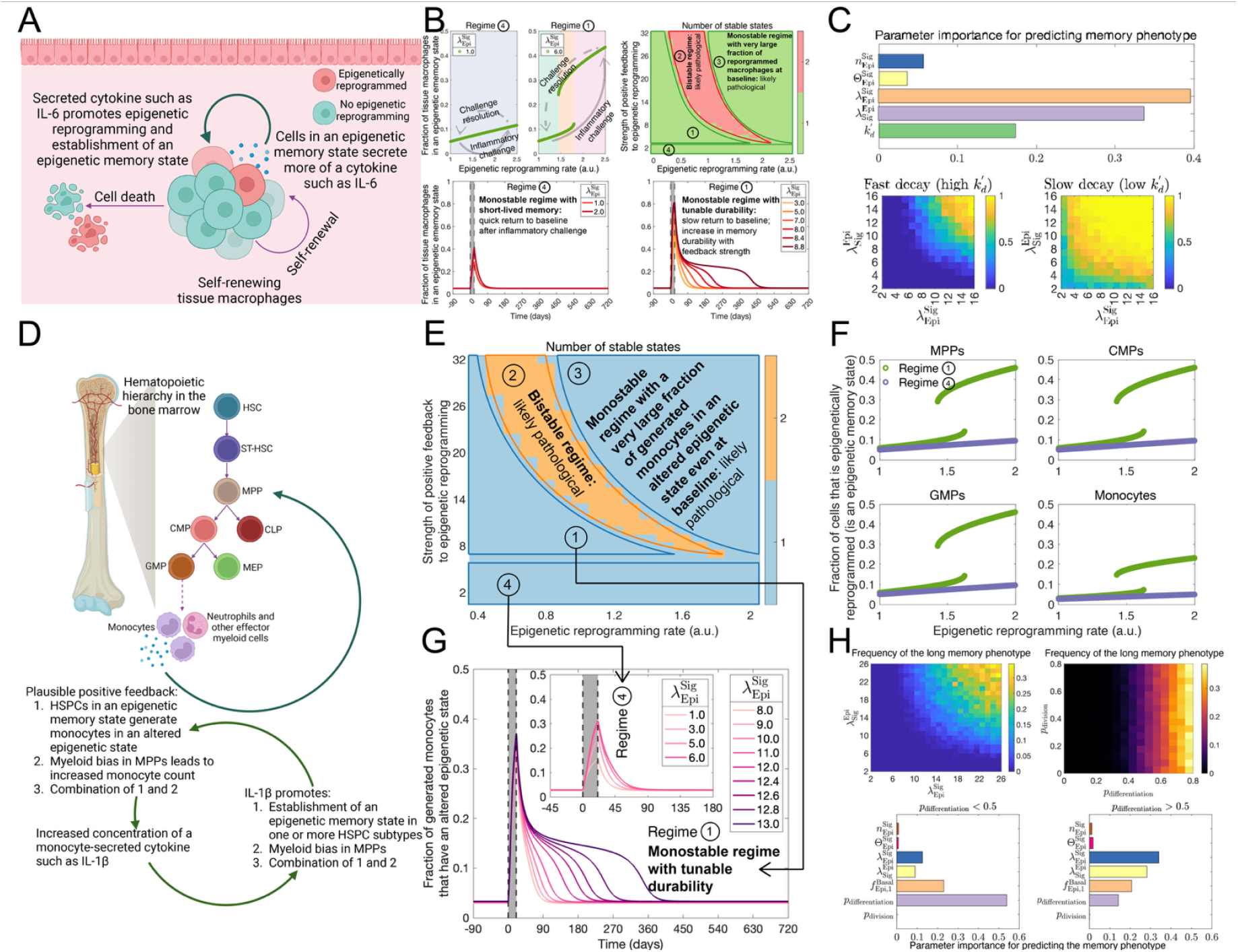
Tuning of trained immunity durability by positive feedback. (A) Schematic showing possible feedback from macrophage-secreted IL-6 to the establishment of an epigenetic memory state in tissue macrophages. (B) Top right: phase diagram showing the number of possible stable states for tissue macrophage epigenetic state dynamics as a function of the epigenetic reprogramming rate (X axis) and strength of positive feedback from a macrophage-secreted cytokine such as IL-6 to the tissue macrophage epigenetic reprogramming (Y axis). See also Figures S2A-D. Bottom: kinetics of decline, after inflammatory challenge resolution, in the fraction of tissue macrophages that are in an epigenetic memory state in regime 1 (bottom right) and in regime 4 (bottom left). Different regimes are shown in the top right panel. 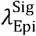 indicates the positive feedback strength with higher values corresponding to stronger feedback. In the two bottom panels, the region shaded in grey indicates the duration of the acute inflammatory challenge; there is no challenge outside of the shaded regions. See also Figures S3F-G. Top left: fraction of epigenetically reprogrammed tissue macrophages as a function of the reprogramming rate in regime 4 (left; very weak feedback) and in regime 1 (right; strong positive feedback). Arrows indicate how the epigenetic reprogramming rate and the reprogrammed macrophage fraction may change in response to an inflammatory challenge (bottom arrow) and upon challenge resolution (top arrow). See also Figure S3E. (C) Top panel: relative importance or contribution of different model parameters to predicting the memory durability phenotype using a random forest model trained on simulated data. Here, 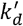 is a composite parameter that determines the memory durability in the absence of feedback (Equations S18-S19). See also Figure S3H-I. Bottom panels: frequency of the long-lasting trained immunity phenotype (memory durability > 180 days) across parameter sets, shown as a function of 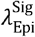 and 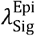. Behavior is shown separately for parameter sets wherein the epigenetically reprogrammed fraction will decay fast (half-life < 14 days) in the absence of feedback (left) and for parameter sets wherein such decay will be slow (half-life > 28 days; right). See also Figure S3G. (D) Schematic showing IL-1*β*-driven positive feedback to the establishment of an epigenetic memory state in one or more HSPC subtypes. An alternate feedback mechanism wherein IL-1*β* induces myeloid bias in hematopoiesis is also shown. Behavior in the case of such an alternate feedback mechanism is shown in Figure S5. (E) Phase diagram showing the number of possible stable steady states for epigenetic state dynamics in the hematopoietic hierarchy as a function of the epigenetic reprogramming rate (X axis) and the strength of positive feedback from a monocyte-secreted cytokine such as IL-1*β* to the epigenetic reprogramming of HSPCs (Y axis). See also Figures S4A-D. (F) Fraction of different HSPC subtypes and monocytes in an epigenetic memory state as a function of the epigenetic reprogramming rate, shown for regimes 1 and 4 in E. (G) Kinetics of decline in the fraction of generated monocytes that are epigenetically reprogrammed post-inflammatory challenge resolution in regime 1 and regime 4 (inset). 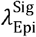 indicates the positive feedback strength with higher values corresponding to stronger feedback. The region shaded in grey indicates the duration of the acute inflammatory challenge. See also Figure S4E-G. (H) Top: frequency of the long-lasting trained immunity phenotype (memory durability > 180 days) across parameter sets, shown as a function of 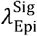 and 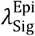 (top left) and as a function of *p*_division_ and *p*_differentiation_ (top right). See also Figure S4H-I. Bottom: relative importance or contribution of different model parameters to predicting the memory durability phenotype using a random forest model when *p*_differentiation_ < 0.5 (bottom left) and when *p*_differentiation_ 0.5 (bottom right). See also Figure S4J-K. Panels A and D were created using Biorender.com.

IL-1*β* is another potential positive feedback mediator, particularly in central trained immunity as reported in multiple studies. Peripheral blood mononuclear cells from elderly individuals vaccinated with the BCG vaccine secrete more IL-1*β* upon stimulation with heat-killed *Candida albicans* in comparison to cells from those receiving placebo.^13^ LSK (Lin-cKit+ Sca1+) cells, a subset that includes HSCs, ST-HSCs, and MPPs, have been shown to undergo epigenetic reprogramming in response to IL-1 *β* stimulation,^8,9^ exhibiting more accessible chromatin at loci bound by transcription factors known to be involved in the IL-1 *β* signaling response including EGR1, IRF1, and RELA. These LSK cells can then generate monocytes expressing elevated levels of IL-1*β* at baseline even in the absence of ongoing inflammatory stimulation.^8^ IL-1*β* can thus drive a positive feedback loop: LSK cells undergo epigenetic reprogramming in response to IL-1*β* and, in turn, generate epigenetically reprogrammed monocytes that upregulate IL-1*β* expression (Figure 3D). There is also evidence for an additional positive feedback mechanism: IL-1*β* has been shown to promote PU.1 expression in HSCs which is associated with myeloid bias in the hematopoietic process.^55,56^ IL-1*β* can also increase the frequency of the myeloid-biased MPP3 subset at the expense of the lymphoid-biased MPP4 subset.^8^ These IL-1 *β*-driven effects could lead to increased monocyte production, and thus, increased IL-1 *β* production, strengthening the IL-1*β*-mediated positive feedback (Figure 3D).

#### Tuning of trained immunity durability by positive feedback

We incorporated feedback from a macrophage-secreted cytokine to the epigenetic state of the same tissue macrophage population into the mathematical model of trained immunity dynamics described above: tissue macrophages in an epigenetic memory state secrete 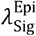-fold more of a cytokine such as IL-6 and the cytokine, in turn, promotes the establishment of the epigenetic memory state (Figure 3A and Equation S20-S21). As in the case of T_M_ cell dynamics (Figure 1E), we find that macrophage epigenetic state dynamics can operate in four distinct behavioral regimes (shown in Figure 3B, top right panel; see also Figure S3A-E) depending on: (a) the baseline epigenetic reprogramming rate (which determines the fraction of tissue macrophages that exhibit an epigenetic memory state in the absence of an inflammatory challenge) and (b) the positive feedback strength, defined as the maximum fold-change in the epigenetic reprogramming rate that the macrophage-secreted cytokine can induce (model parameter 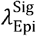). Just as in the case of the bystander T cell activation response above, regime 1, with strong feedback, is characterized by a switch-like, excitable acute response to inflammatory challenges: the fraction of macrophages in an epigenetic memory state can increase multi-fold from a baseline homeostatic level in response to the challenge, followed by a multi-fold decline and subsequent return to the baseline fraction once the challenge has resolved (Figure 3B, top right panel). The acute response and return to homeostasis involve transitioning between regimes 1 and 3 via regime 2, and the dynamics exhibit hysteresis (Figure 3B, top left panel). In regime 4, where the feedback is weak, there is no regime switching or hysteresis during the acute response, and the increase in the epigenetically reprogrammed fraction is proportionate to the inflammatory challenge magnitude (Figure 3B, top left panel). Importantly, while regime 4 involves fast decline in the fraction of epigenetically reprogrammed macrophages once the inflammatory challenge has resolved (Figure 3B, bottom left panel), return to the pre-challenge baseline in regime 1 is slow and depends on the feedback strength (Figures 3B, bottom right panel and S3G). For a fixed value of *p*_division_, trained immunity durability increases non-linearly with the feedback strength (*p*_division_ = 0.5 in Figures 3B and S3G). Thus, strong positive feedback can relax the constraint on trained immunity from error-prone epigenetic state transmission during self-renewal. To further assess this effect across broad ranges of parameter values, we simulated the model dynamics for an ensemble of biologically plausible parameter sets (SI Section IIC) and trained a random forest model to predict the memory durability phenotype for each parameter set (Figure S3H-I). As in the case of bystander activation memory above, the overall feedback strength (determined together by 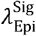 and 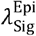) is a key determinant of the memory durability phenotype (Figure 3C, top panel). Consistently, in the strong feedback regime (*i.e*., high values of 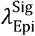 and 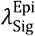), long-lasting trained immunity is possible even in scenarios with short macrophage half-life (high *k*_*d*_) and low fidelity of epigenetic memory state transmission during self-renewal, *i.e*., low p_division_, (Figure 3C, bottom panels), scenarios that would otherwise be characterized by rapid decay of epigenetic memory (Equation S19).

We next tested the effect of positive feedback on central trained immunity by incorporating IL-1*β*-mediated feedback into our hematopoiesis model: IL-1*β* can promote the epigenetic reprogramming of MPPs, CMPs, and GMPs, and these reprogrammed progenitors ultimately generate monocytes with an altered epigenetic state that secrete 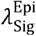-fold more IL-1*β* (Figure 3D and Equation S46-S47). Figures 3E and S4A-E show that in the presence of such feedback, we once again have four dynamical regimes as in the cases of T_M_ cell and tissue macrophage dynamics. The regime is determined by (a) the rate of progenitor epigenetic reprogramming at baseline (which determines the fraction of monocytes in an altered epigenetic state in the absence of any inflammatory challenge) and (b) the feedback strength, encoded by the effect of IL-1*β* on the epigenetic reprogramming of progenitor cells and defined quantitatively as the maximum fold-change in the reprogramming rate that IL-1*β* can induce (model parameter 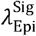). Regime 1, with strong positive feedback, is once again characterized by a switch-like, excitable acute response (Figure 3F) and exhibits tunable memory durability given the slow, feedback strength-dependent return to the pre-challenge state after the inflammatory challenge has resolved (Figures 3G and S4G). For fixed *p*_division_ and *p*_differentiation_ values, the durability of central trained immunity increases with the feedback strength in regime 1 (Figure S4G). Thus, it is no longer strictly constrained by epigenetic state transmission fidelity. Simulations for an ensemble of biologically plausible parameter sets (SI Section IIIC) confirmed the association of strong feedback with long-lasting trained immunity (Figures 3H, top left panel, S4H, and S4K). Importantly, we find that across the parameter sets in the ensemble, very high values of *p*_division_ are no longer necessary for long-lasting trained immunity (Figures 3H, top right panel and S4I; compare with Figure 2G). The memory durability, however, remains dependent on *p*_differentiation_ since this parameter determines the fidelity of epigenetic state transmission from GMPs to the generated monocytes (Figure 3H, top right panel). We next trained a random forest model to predict the memory durability phenotype for each parameter set (Figures S4J and 3H) and found that while the value of *p*_differentiation_ is a key contributor to the prediction when transmission fidelity during differentiation is low (for *p*_differentiation_ < 0.5), the overall feedback strength (determined together by 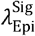 and 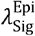) becomes the key contributor when *p*_differentiation_ > 0.5 (Figure 3H, bottom panels). There is no contribution from *p*_division_ in either scenario.

We also modeled the scenario wherein an inflammatory challenge induces myeloid bias in MPPs, thus increasing monocyte production (Figures 3D and S5A). We find that the memory durability, now defined as the duration after challenge resolution for which the monocyte counts remain elevated compared to the pre-challenge baseline, is dependent on the fidelity with which the myeloid-biased state is retained during MPP self-renewal (model parameter 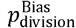). Here, long-lasting memory is possible only when 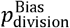 is very close to 1 (Figure S5A) and the memory durability is not tunable. We then incorporated positive feedback from monocyte-secreted IL-1 *β* to myeloid-bias induction and find that the overall behavior (Figure S5B-I) is qualitatively similar to the previously discussed positive feedback cases, *i.e*., tunable, long-lasting memory that depends on the feedback strength and is no longer constrained by 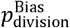 (Figure S5I-L).

Our analysis thus shows that the presence of positive feedback can drive tunable trained immunity durability, overcoming the constraints due to short effector cell half-life and poor / variable epigenetic state transmission fidelity during self-renewal and differentiation. While we have analyzed specific positive feedback cases (IL-6 signaling and IL-1*β* signaling), our approach as well as the principles revealed are generalizable. For example, our mathematical model can be adapted to scenarios involving other feedback mediators or mechanisms such as increased myelopoiesis driven by a secretory MPP subset.^57^

### Potential experiments to assess the role of feedback in antigen-agnostic immune memory

Our models predict that both bystander activation memory and trained immunity can exhibit tunable durability in the presence of positive feedback. Thus, experimentally quantifying the durability itself *in vivo* is the first step towards assessing the role of feedback in antigen-agnostic immune memory. Existing experimental studies have largely focused on demonstrating the development of such memory following acute responses to inflammatory challenges. These studies test for the presence of memory and its functional consequences at a single timepoint after the initial challenge (typically after 2-4 weeks in mouse studies^6,8–10^). The same experimental setup can be used to investigate the memory durability: after exposing all animals to an initial challenge, subsequent challenges can be introduced to different animals at different time points to estimate the memory durability (Figure 4A). In addition to quantifying memory durability, the setup can be used to simultaneously track immune circuits components underlying the memory, *e.g*., T_M_ cell and monocyte counts, and levels of cytokines such as IFN-*γ*, IL-6, and IL-1*β*, and to assess their decay kinetics after the initial challenge by measuring these quantities before the subsequent challenge in each case (Figure 4B). Given that the immune circuits with cytokine-mediated feedback we have studied here operate within tissues, it is important to measure the relevant cell counts and cytokine levels in tissues. If these immune circuit components decay at similar timescales as the memory response (Figure 4C), it would support their role in the maintenance of antigen-agnostic immune memory.

**Figure 4.**
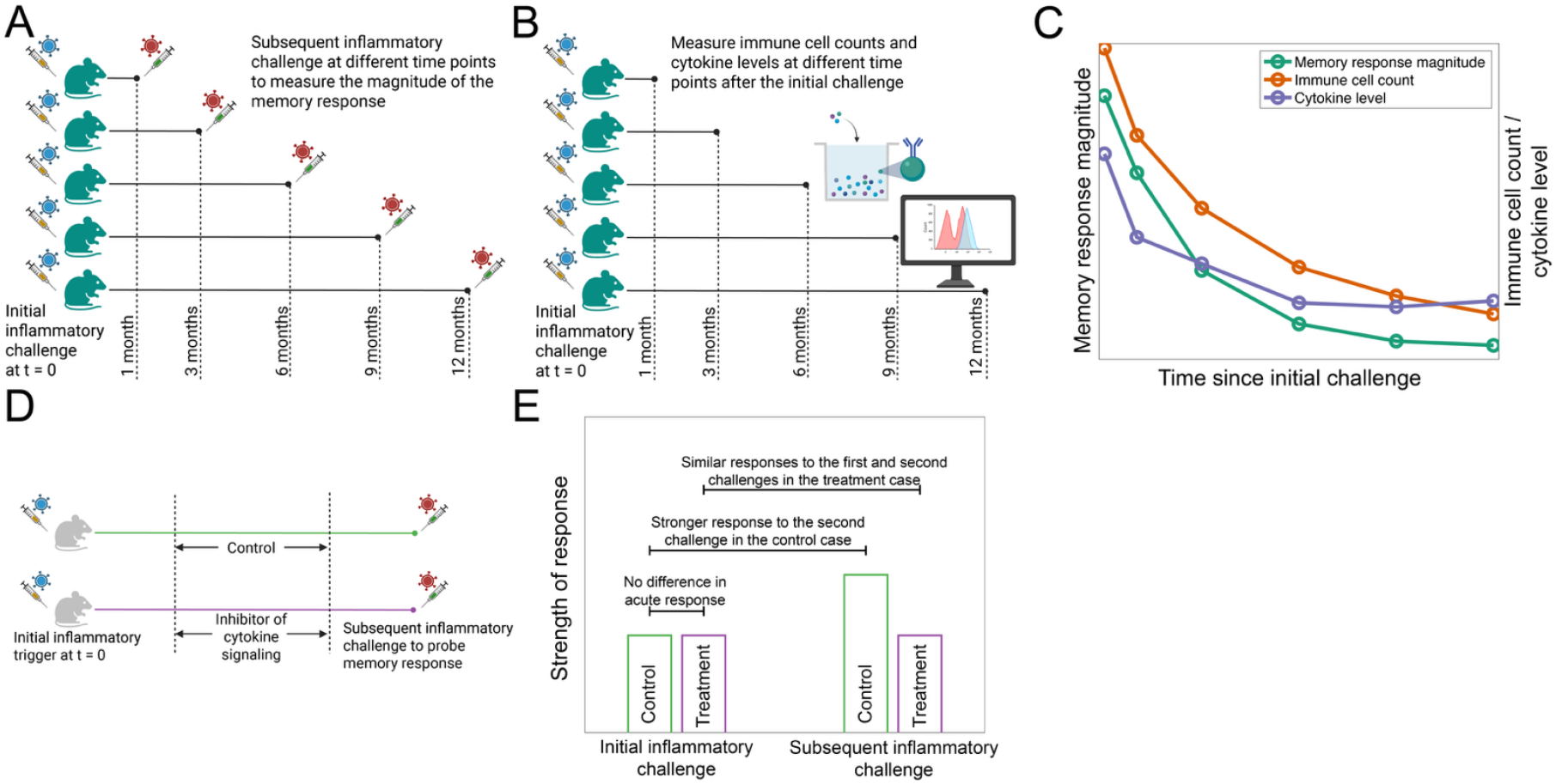
Potential experiments to assess the role of feedback in antigen-agnostic immune memory. (A) Experimental design to determine the durability of antigen-agnostic immune memory by probing the memory response at multiple time points including long after the initial inflammatory challenge. (B) Measuring cell counts and cytokine levels before each subsequent inflammatory challenge to determine the decay kinetics of these immune circuits components after the initial challenge. (C) Comparing the timescale of decay in the memory response with the timescale of change in immune circuit components such as cell counts and cytokine levels would help identify a role for cytokine-mediated feedback in tuning memory durability. (D) Experimental setup to directly test the role of cytokine-mediated feedback in the maintenance of antigen-agnostic immune memory. Cytokine signaling is inhibited after the acute inflammatory challenge has resolved and the inhibition withdrawn before introducing the second challenge so that the inhibition does not directly affect the acute response. (E) Predicted outcome of the experiment in D if cytokine-mediated positive feedback plays a key role in the maintenance of antigen-agnostic immune memory. Panels A, B, and D were created with Biorender.com.

Bystander activation memory lasting longer than the estimate based on the half-life of T_M_ cells (around 14 days in mice;^25^ Figure 1B) would suggest a role for additional mechanisms in determining memory durability. For trained immunity, memory durability in the absence of positive feedback is dependent on two key parameters, *p*_division_ and *p*_differentiation_ (Figure 2). While the exact values of these parameters are not known, they are expected to be less than one, likely significantly so.^43,44,48^ Since the self-renewal and differentiation rates of hematopoietic progenitors and myeloid effectors in mice have been estimated,^39^ the mathematical framework described here can be used to infer both *p*_division_ and *p*_differentiation_ from the trained immunity durability values once they have been measured. If the estimated values of *p*_division_ and / or *p*_differentiation_ are close to 1, indicating very high fidelity of epigenetic state transmission during self-renewal and / or differentiation, it would contradict expectations since such very high fidelity is biologically unlikely, and thus support a role for additional mechanisms in tuning trained immunity durability.

Contribution of positive feedback to determining memory durability can be directly tested by determining how the memory response changes upon inhibiting signaling from the cytokine mediating the feedback (Figure 4D-E). In the case of bystander activation memory, a role for IFN-*γ* signaling-driven feedback can be directly tested by determining if treatment with an IFN-*γ* signaling inhibitor after the initial challenge abolishes or weakens bystander activation memory. A role for positive feedback in tuning trained immunity durability can be confirmed by testing if treatment with an inhibitor of cytokine signaling mediating the positive feedback (an IL-6 or IL-1*β* signaling inhibitor, for example) after the initial inflammatory challenge has resolved abolishes or weakens trained immunity. Thus, predictions from our mathematical modeling-based analysis can be experimentally tested in multiple ways to either further support or contradict our proposed theory of tunable antigen-agnostic memory durability.

## DISCUSSION

Recent studies have highlighted the development and function of antigen-agnostic immune memory.^6–12,14^ However, a quantitative, mechanistic understanding of the maintenance of such memory has been lacking. Towards this end, here we developed mathematical models of the dynamics of bystander activation memory and trained immunity, two components of antigen-agnostic immune memory. In both cases, we identified constraints on memory durability due to immune cell turnover, dependence on cytokines with limited availability, and epigenetic information transmission fidelity. We then show that positive feedback mediated by known cytokine signaling pathways can result in immune circuits that exhibit switch-like, excitable acute response to inflammatory challenges followed by slow return to pre-challenge baseline, resulting in tunable memory durability. Our analysis highlights features shared across antigen-agnostic memory components encoded by different mechanisms (Table 2) and demonstrates how these features are critical determinants of both the magnitude of the acute response and the durability of the resultant memory. We show that in the presence of cytokine-mediated positive feedback, antigen-agnostic immune memory can be encoded by the state of the overall immune circuit rather than by the intrinsic state of individual cells or the cell population size. This makes the memory durability less dependent on cell intrinsic parameters such as lifespan and fidelity of epigenetic state transmission during self-renewal and differentiation, and more dependent on circuit-level parameters such as the feedback strength. Ultimately, this allows for the memory to be tunable and more durable than permitted by the dynamics of the individual immune cells involved.

**Table 2.**
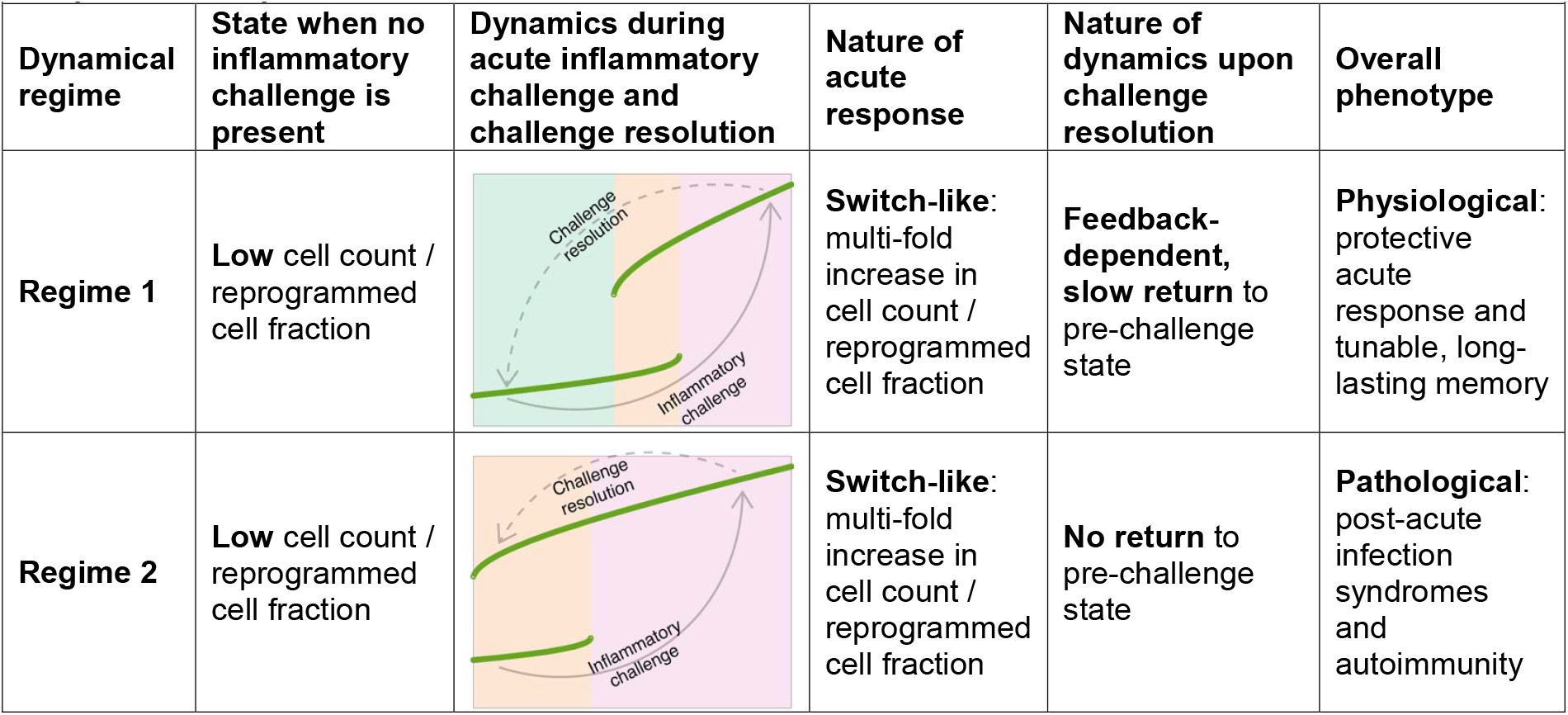
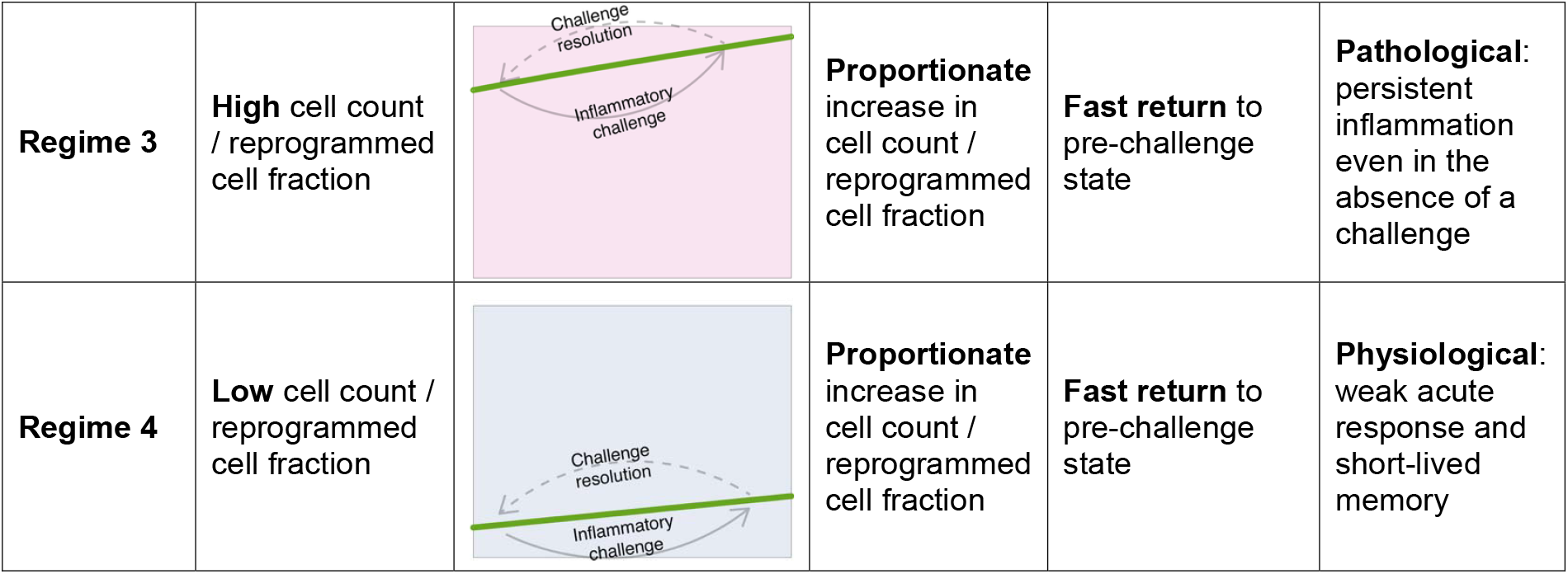
Overview of the different dynamical regimes immune cell circuits modeled here an exhibit in the presence of positive feedback.

| Dynamical regime | State when no inflammatory challenge is present | Dynamics during acute inflammatory challenge and challenge resolution | Nature of acute response | Nature of dynamics upon challenge resolution | Overall phenotype |
| --- | --- | --- | --- | --- | --- |
| **Regime 1** | **Low** cell count / reprogrammed cell fraction |  | **Switch-like:** multi-fold increase in cell count / reprogrammed cell fraction | **Feedback-dependent, slow return** to pre-challenge state | **Physiological:** protective acute response and tunable, long-lasting memory |
| **Regime 2** | **Low** cell count / reprogrammed cell fraction |  | **Switch-like:** multi-fold increase in cell count / reprogrammed cell fraction | **No return** to pre-challenge state | **Pathological:** post-acute infection syndromes and autoimmunity |
| **Regime 3** | **High** cell count / reprogrammed cell fraction |  | **Proportionate** increase in cell count / reprogrammed cell fraction | **Fast return** to pre-challenge state | **Pathological:** persistent inflammation even in the absence of a challenge |
| **Regime 4** | **Low** cell count / reprogrammed cell fraction |  | **Proportionate** increase in cell count / reprogrammed cell fraction | **Fast return** to pre-challenge state | **Physiological:** weak acute response and short-lived memory |

In addition to tuning memory durability, we show that positive feedback in immune cell circuits can drive a switch-like, excitable response to acute challenges, wherein the T_M_ cell count (in the case of bystander T_M_ cell activation) and the fraction of myeloid effectors in an altered epigenetic state (in the case of trained immunity) can increase multi-fold in response to a strong inflammatory challenge. Such a response could be advantageous when responding to an acute infection, for example, compared to the proportionate response when there is no or weak positive feedback. Given that feedback strength can tune both the magnitude of the antigen-agnostic acute response and the memory durability, these two can be correlated. However, when the positive feedback is too strong, an inflammatory challenge can cause a permanent switch to a persistently inflamed state. This is likely pathological; for example, it can lead to post-acute infection syndromes^34^ and autoimmunity.^9,35,58,59^ Thus, the feedback strength and other parameters that together determine both the acute response magnitude and memory durability are likely under evolutionary constraints to achieve strong, long-lasting protective responses while avoiding persistent inflammation.

Note that our goal in this study is not to develop detailed mathematical models that incorporate extensive molecular and cellular details of bystander T_M_ cell activation and trained immunity, followed by estimating model parameters by fitting to experimental datasets. Instead, we adopted a coarse-grained modeling approach and used the previously described MAPPA^37,38^ approach, simulating the mathematical models for a large number of biologically plausible parameter sets followed by using a machine learning approach to analyze model behavior. This approach allowed us to explore the range of possible behaviors of immune cell circuits, spanning both physiological and pathological scenarios. Aside from circumventing the challenge arising from the dearth of quantitative experimental data on antigen-agnostic memory durability and related parameters, here, by not restricting the analysis to parameter sets that fit specific experimental datasets, our approach enabled the discovery of novel relationships between molecular and cellular parameters and immune phenotypes such as the identification of the feedback strength, that may be encoded by different mechanisms in different types of immune circuits, as a key determinant of memory durability.

The immune cell circuits modeled here likely do not operate in isolation *in vivo*. For example, both monocytes and macrophages present IL-15 to T_M_ cells, and this presentation is upregulated in response to IFN-*γ*.^27,31,60,61^ IFN-*γ* has also been shown to modulate hematopoiesis, including suppression of HSC self-renewal and induction of myeloid bias.^62^ Thus, the T_M_ cell bystander activation circuit may be coupled with the tissue macrophage and / or hematopoiesis circuits to model the interplay between bystander activation and trained immunity. In general, by incorporating the interplay between immune cells and signaling pathways, complex circuits can be constructed. Such circuits, analyzed using our MAPPA approach for example, could shed light on how complex baseline immune setpoints, encoded by the relative abundances of immune cell types together with their transcriptional and epigenetic states,^63^ are established, maintained, and modulated by inflammatory challenges,^11,64^ and how such setpoints can modulate, both qualitatively and quantitatively, the response to subsequent infection or vaccination.^11,65–67^

### Limitations of the study

The present study makes predictions concerning memory durability in the absence and presence of positive feedback. However, the contribution of positive feedback to memory durability *in* vivo remains to be assessed experimentally. Contributions to tunable memory from mechanisms other than positive feedback cannot be ruled out based on the analysis presented herein. Finally, in order to keep the mathematical modeling task manageable, we have made multiple assumptions (listed in SI Sections IE, IID, and IIID) and these may need to be revisited as more experimental data becomes available. Our overall approach and conceptual framework, however, is generic and can be adapted to incorporate new experimental data.

## ACKNOWLEDGEMENTS

This study was supported by National Institutes of Health (NIH) grant nos. R01AI170116 and U01AI153700 (awarded to J.S.T.), and by a Chan Zuckerberg Biohub Investigator Award (to J.S.T.). S.T. is supported by the Yale-Boehringer Ingelheim Biomedical Data Science Fellowship.

## AUTHOR CONTRIBUTIONS

S.T. and J.S.T. conceived and designed the study. S.T. carried out the research. S.T., C.L., W.W.L., and R.S. interpreted the data. S.T. and J.S.T. wrote the manuscript with input from the other authors.

---

**Figure S1.**
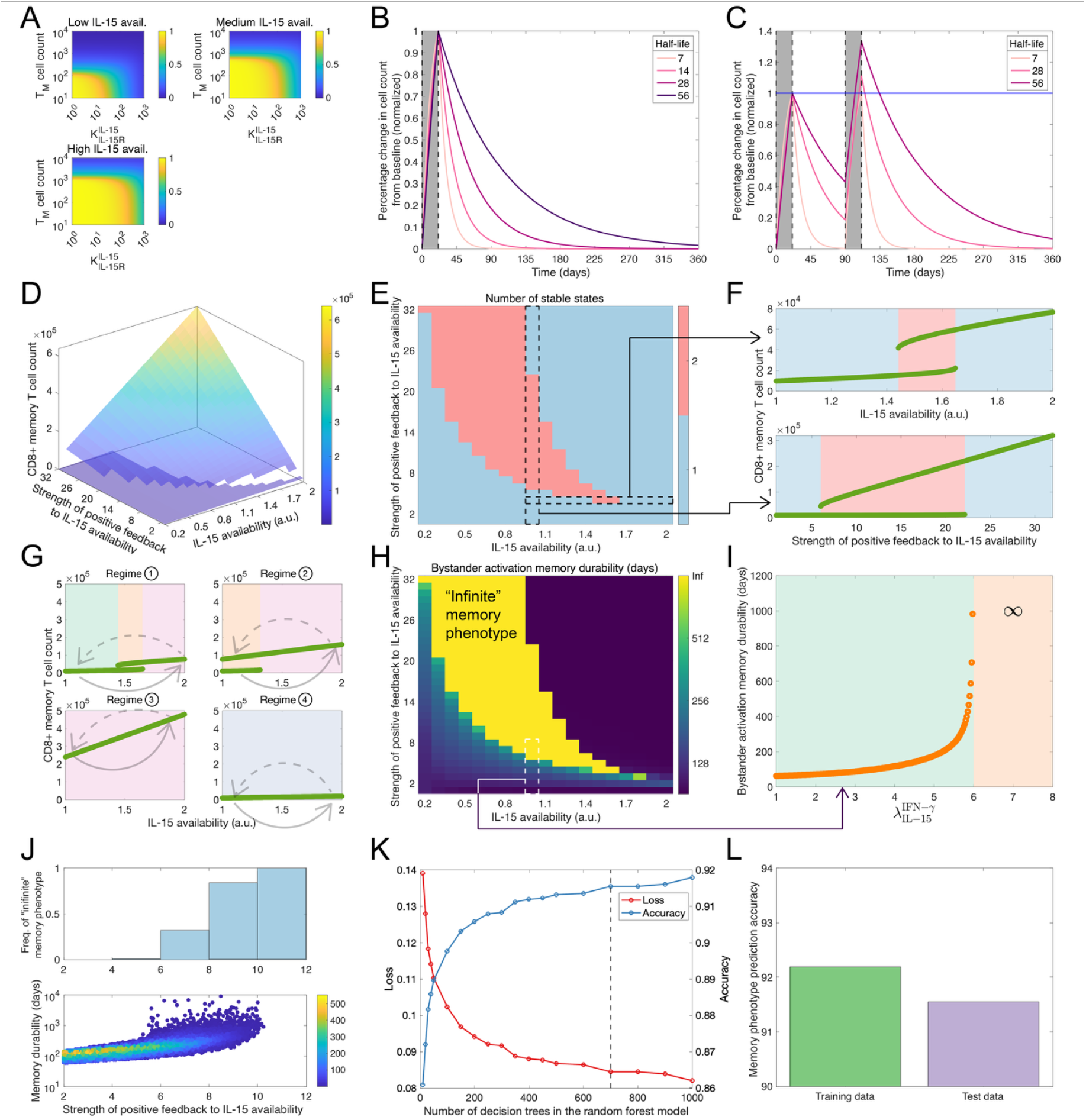
Modeling bystander activation memory (related to Figure 1). (A) Fraction of T_M_ cells in the population that receive IL-15 stimulation at any given point in time, shown as a function of the T_M_ cell count and the effective Michaelis constant for IL-15 binding to the IL-15 receptor expressed by T_M_ cells. Different panels show the fraction for different levels of IL-15 availability. See SI Section IB (Equations S2-S3) for the mathematical description. (B) Kinetics of T_M_ cell population contraction post-inflammatory challenge resolution in the absence of any feedback from T_M_ cell-secreted IFN-*γ* to IL-15 availability, shown for different values of the T_M_ cell half-life. The region shaded in grey indicates the duration of the acute inflammatory challenge during which the T_M_ population expands in response to the increase in IL-15 availability triggered by the challenge. (C) T_M_ cell population kinetics during two consecutive inflammatory challenges. Periods of acute challenges are indicated by the regions shaded in grey. For the case with a half-life of 7 days, T_M_ cell count returns to homeostatic, pre-first challenge levels by the time of the second challenge, and the acute T_M_ cell response to the second challenge is the same as the response to the first challenge. For the cases with longer half-lives, T_M_ cell counts at the onset of the second challenge are higher that the counts at the onset of the first challenge. Consequently, the acute T_M_ cell response to the second challenge is stronger than the response to the first challenge. For each half-life value, the T_M_ cell count is normalized by the peak count during the first inflammatory challenge. (D) Steady state T_M_ cell counts as a function of the baseline IL-15 availability and the strength of positive feedback from T_M_ cell-secreted IFN-*γ* to IL-15 availability. (E) Same as Figure 1E. (F) Top: steady state T_M_ cell counts as a function of the baseline IL-15 availability, shown for the feedback strength indicated in E. Bottom: steady state T_M_ cell counts as a function of the strength of positive feedback from T_M_ cell-secreted IFN-*γ* to IL-15 availability, shown for the baseline IL-15 availability indicated in E. (G) T_M_ cell count as a function of the IL-15 availability in different regimes. Arrows indicate how the IL-15 availability and the T_M_ cell count may change in response to an inflammatory challenge (solid arrow) and upon challenge resolution (dashed arrow). Different regimes are shown in Figure 1E. (H) Bystander activation memory durability shown as a function of the IL-15 availability at baseline, *i.e*., before the onset of the inflammatory challenge, and of the strength of the positive feedback from T_M_ cell-secreted IFN-*γ* to IL-15 availability. The “infinite” memory phenotype corresponds to bistable regime 2. (I) Bystander activation memory durability as a function of the positive feedback strength 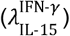, shown for the IL-15 availability indicated in H. The orange shaded regime corresponds to the “infinite” memory scenario: in this regime, T_M_ cell counts may never return to the pre-challenge baseline after the inflammatory challenge has resolved. (J-L) Parameter-phenotype mapping analysis. (J) Durability of bystander activation memory for biologically plausible parameter sets, shown as a function of the positive feedback strength 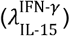. Bottom: memory durability in the case of parameter sets for which the durability is finite. Color indicates the number of parameter sets. Top: frequency of the “infinite” memory phenotype when the feedback strength is within the indicated ranges. (K) 10-fold cross-validation performance of random forest models with different numbers of decision trees at predicting the bystander activation memory phenotype. Here, loss is the misclassification error and thus, loss and accuracy add to 1. The dashed black line indicates the random forest model size that was used for the analyses shown in L and in Figure 1I. (L) Performance of a 700-tree random forest model on training and test datasets. Additional details concerning how each panel was generated are in SI Section IC.

**Figure S2.**
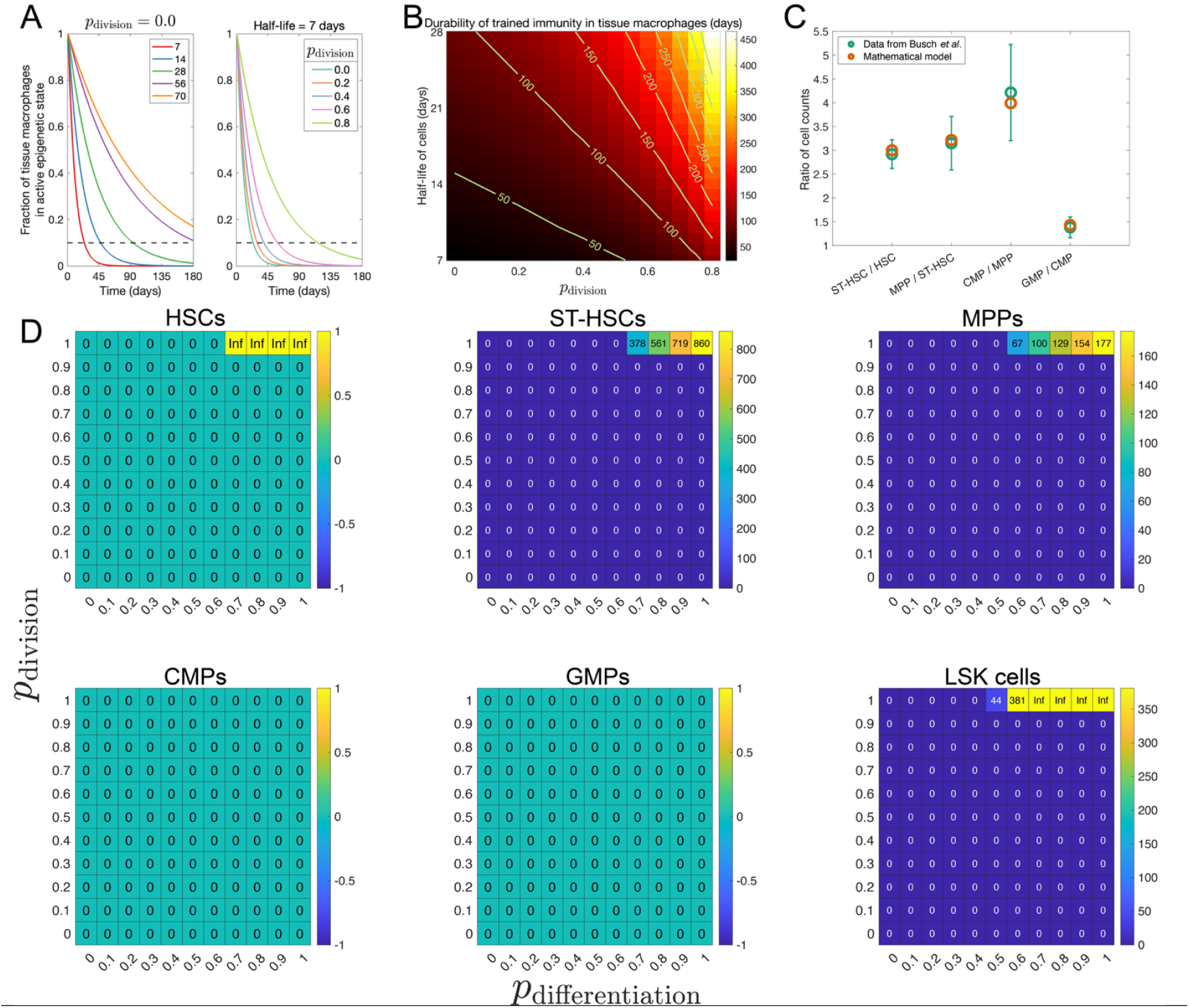
Durability of trained immunity in the absence of feedback (related to Figure 2). (A) Kinetics of decline in the fraction of tissue macrophages that are in an epigenetic memory state, shown for different values of the macrophage half-life (left panel) and *p*_division_ (right panel). The dashed black line indicates a fraction of 10%. (B) Durability of trained immunity as encoded by epigenetically reprogrammed tissue macrophages, shown as a function of the macrophage half-life and *p*_division_. (C) Confirmation that ratios of steady state counts of different HSPC subtypes in our mathematical model of hematopoiesis match those reported in Bush *et al*.^39^ (D) Durability of trained immunity as encoded by epigenetically reprogrammed monocytes, shown as a function of *p*_division_ and *p*_differentiation_, when the inflammatory challenge epigenetically reprograms different HSPC subtypes. The last panel is the same as Figure 2G. Additional details concerning how each panel was generated are in SI Section IIC for panels A-B and in SI Section IIIC for panels C-D.

**Figure S3.**
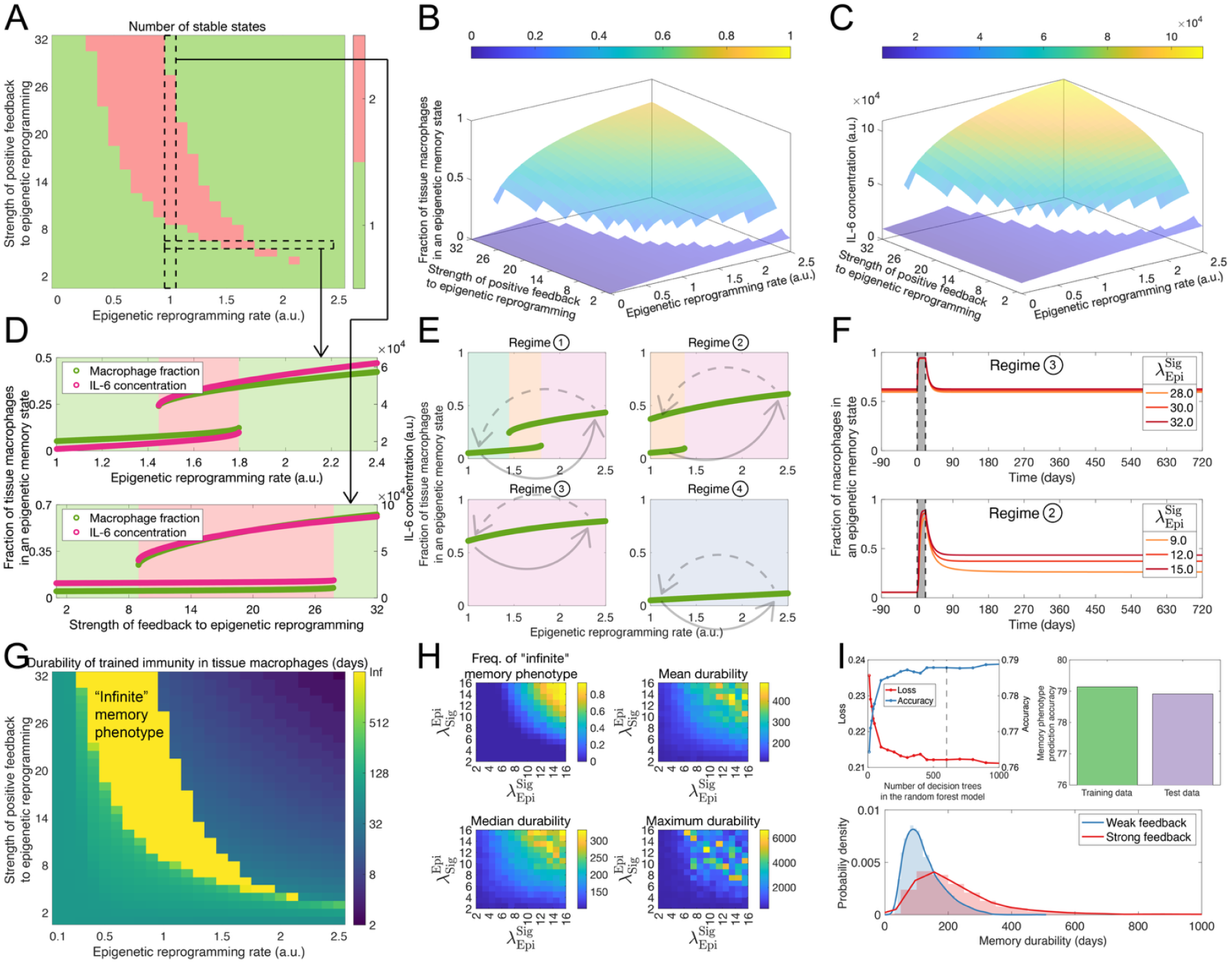
Tuning of peripheral trained immunity durability by positive feedback (related to Figure 3A-C). (A) Same as the top right panel in Figure 3B. (B) Fraction of tissue macrophages in an epigenetic memory state as a function of the baseline epigenetic reprogramming rate and the strength of positive feedback from a macrophage-secreted cytokine such as IL-6 to the epigenetic reprogramming rate. (C) IL-6 concentration as a function of the baseline epigenetic reprogramming rate and the positive feedback strength. (D) Top: fraction of tissue macrophages in an epigenetic memory state and IL-6 concentration as a function of the baseline epigenetic reprogramming rate, shown for the feedback strength highlighted in A. Bottom: fraction of tissue macrophages in an epigenetic memory state and IL-6 concentration as a function of the feedback strength, shown for the baseline reprogramming rate highlighted in A. (E) Fraction of tissue macrophages in an epigenetic memory state as a function of the reprogramming rate in different regimes. Arrows indicate how the reprogramming rate and the macrophage fraction in an epigenetic memory state may change in response to an inflammatory challenge (solid arrow) and upon challenge resolution (dashed arrow). Different regimes are shown in the top right panel of Figure 3B. (F) Kinetics of decline in the fraction of tissue macrophages in an epigenetic memory state in regime 2 (bottom) and in regime 4 (top).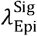 indicates the positive feedback strength. In both panels, the region shaded in grey indicates the duration of the acute inflammatory challenge. (G) Peripheral trained immunity durability shown as a function of the epigenetic reprogramming rate at baseline, *i.e*., in the absence of any inflammatory challenge, and of the positive feedback strength. The “infinite” memory phenotype corresponds to bistable regime 2. (H) Peripheral trained immunity durability across the ensemble of biologically plausible parameter sets, shown as a function of 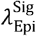 and 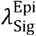, the two model parameters that together determine the positive feedback strength. (I) Parameter-phenotype mapping analysis. Top left: 10-fold cross-validation performance of random forest models with different numbers of decision trees at predicting the durability phenotype. Here, loss is the misclassification error and thus, loss and accuracy add to 1. The dashed black line indicates the random forest model size that was used for the analyses shown in the top right panel and in Figure 3C (top panel). Top right: performance of a 600-tree random forest model on training and test datasets. Bottom: distribution of memory durability values in the weak feedback 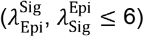 and strong feedback 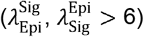 cases. Additional details concerning how each panel was generated are in SI Section IIC.

**Figure S4.**
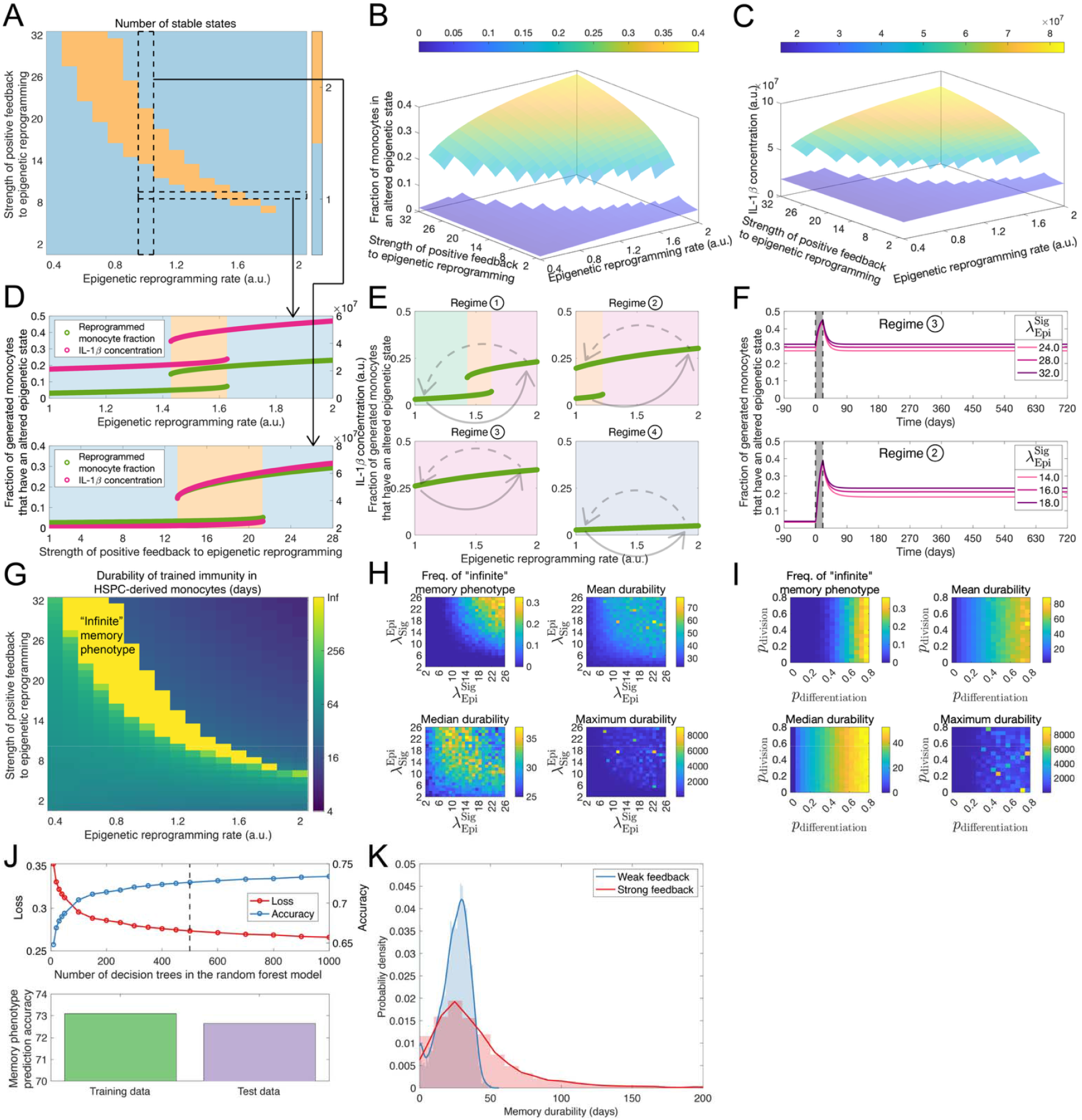
Tuning of central trained immunity durability by positive feedback (related to Figure 3D-H). (A) Same as Figure 3E. (B) Fraction of generated monocytes that are in an altered, reprogrammed epigenetic state as a function of the baseline HSPC epigenetic reprogramming rate and the strength of positive feedback from a monocyte-secreted cytokine such as IL-1*β* to the HSPC epigenetic reprogramming rate. (C) IL-1*β* concentration as a function of the baseline HSPC epigenetic reprogramming rate and the positive feedback strength. (D) Top: fraction of generated monocytes that are in an altered epigenetic state and the IL-1*β* concentration as a function of the baseline reprogramming rate, shown for the feedback strength highlighted in A. Bottom: fraction of generated monocytes that are in an altered epigenetic state and the IL-1*β* concentration as a function of the feedback strength, shown for the baseline reprogramming rate highlighted in A. (E) Fraction of generated monocytes that are in an altered epigenetic state as a function of the reprogramming rate in different regimes. Arrows indicate how the reprogramming rate and the monocyte fraction in an altered epigenetic state may change in response to an inflammatory challenge (solid arrow) and upon challenge resolution (dashed arrow). Different regimes are shown in Figure 3E. (F) Kinetics of decline in the fraction of generated monocytes that are epigenetically reprogrammed (*i.e*., are in an altered epigenetic state) in regime 2 (bottom) and in regime 4 (top).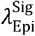 indicates the positive feedback strength. In both panels, the region shaded in grey indicates the duration of the acute inflammatory challenge. (G) Central trained immunity durability shown as a function of the HSPC epigenetic reprogramming rate at baseline, *i.e*., in the absence of any inflammatory challenge, and of the positive feedback strength. The “infinite” memory phenotype corresponds to bistable regime 2. (H) Central trained immunity durability across the ensemble of biologically plausible parameter sets, shown as a function of 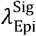 and 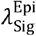, the two model parameters that together determine the positive feedback strength. (I) Central trained immunity durability across the ensemble of biologically plausible parameter sets, shown as a function of *p*_division_ and *p*_differentiation_. (J) Parameter-phenotype mapping analysis. Top panel: 10-fold cross-validation performance of random forest models with different numbers of decision trees at predicting the memory durability phenotype. Here, loss is the misclassification error and thus, loss and accuracy add to 1. The dashed black line indicates the random forest model size that was used for the analyses shown in the bottom panel and in Figure 3H (bottom panels). Bottom panel: performance of a 500-tree random forest model on training and test datasets. (K) Distribution of memory durability values in the weak feedback 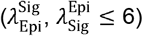 and strong feedback 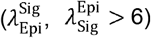 cases. Additional details concerning how each panel was generated are in SI Section IIIC2.

**Figure S5.**
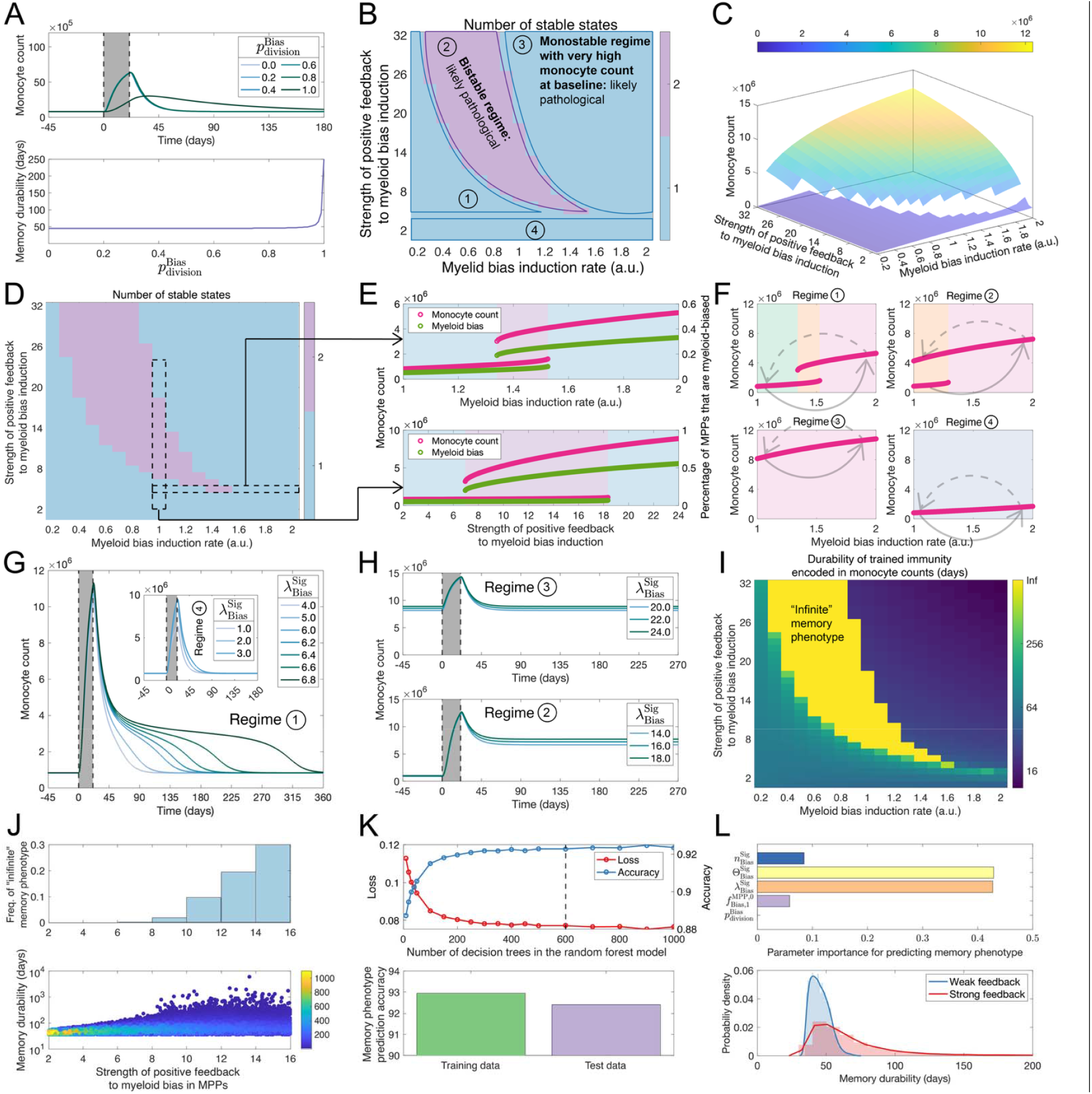
Tuning of the durability of monocyte count-encoded central trained immunity by positive feedback (related to Figure 3D). (A) Top: kinetics of decline in monocyte counts after an acute inflammatory challenge in the absence of feedback from a monocyte-secreted cytokine to myeloid bias in MPPs. Behavior is shown for different values of 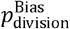 which is the fidelity with which the epigenetic state associated with myeloid bias is transmitted during the self-renewal of MPPs. The region shaded in grey indicates the acute inflammatory challenge during which the challenge induces myeloid bias in MPPs, leading to an increase in monocyte production. Bottom: durability of memory, as encoded by elevated monocyte counts, as a function of 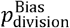. Here, memory durability is defined as the length of the period post-inflammatory challenge resolution during which the monocyte count is at least 10% above the pre-challenge baseline. (B) Phase diagram showing the number of possible stable states of monocyte count dynamics as a function of the baseline myeloid bias induction rate and the strength of positive feedback from a monocyte-secreted cytokine such as IL-1*β* to myeloid bias induction. (C) Monocyte count as a function of the baseline myeloid bias induction rate and the strength of feedback from a monocyte-secreted cytokine to myeloid-bias induction. (D) Same as B. (E) Top: monocyte count and the fraction of MPPs that are myeloid-biased as a function of the baseline bias induction rate, shown for the feedback strength highlighted in D. Bottom: monocyte count and the fraction of MPPs that are myeloid-biased as a function of the feedback strength, shown for the baseline bias induction rate highlighted in D. (F) Monocyte count as a function of the myeloid bias induction rate in different regimes. Arrows indicate how the myeloid bias induction rate and the monocyte count may change in response to an inflammatory challenge (solid arrow) and upon challenge resolution (dashed arrow). Different regimes are shown in B. (G-H) Kinetics of decline in monocyte counts in different regimes. 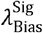 indicates the positive feedback strength, defined as the maximum fold-change in the myeloid bias induction rate that the monocyte-secreted cytokine can cause; higher values of 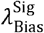 thus correspond to stronger feedback. The region shaded in grey indicates the duration of the acute inflammatory challenge. (I) Durability of central trained immunity as encoded by monocyte counts, shown as a function of the baseline myeloid-bias induction rate (*i.e*., the bias induction rate in the absence of an inflammatory challenge) and the positive feedback strength. The “infinite” memory phenotype corresponds to bistable regime 2. (J) Memory durability across the ensemble of biologically plausible parameter sets, shown as a function of the positive feedback strength. Bottom panel: memory durability in the case of parameter sets for which the durability is finite. Color indicates the number of parameter sets. Top panel: frequency of the “infinite” memory phenotype when the feedback strength is within the indicated ranges. (K-L) Parameter-phenotype mapping analysis. (K) Top panel: 10-fold cross-validation performance of random forest models with different numbers of decision trees at predicting the memory durability phenotype. Here, loss is the misclassification error and thus, loss and accuracy add to 1. The dashed black line indicates the random forest model size that was used for the analyses shown in the bottom panel and in I (top panel). Bottom panel: performance of a 600-tree random forest model on training and test datasets. (L) Top panel: relative importance or contribution of different model parameters to predicting the memory durability phenotype using a random forest model. Bottom panel: Distribution of memory durability values for weak feedback 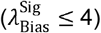 and strong feedback 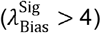. Additional details concerning how each panel was generated are in SI Section IIIC3.

