## Supplemental Information for "Constraints and tunability of antigen-agnostic memory durability"

### SUPPLEMENTAL INFORMATION
### Constraints and tunability of antigen-agnostic memory durability

Shubham Tripathi,[1] Can Liu,[1] William W. Lau,[2] Rachel Sparks,[2] and John S. Tsang[1,3,4,*]

[1] *Yale Center for Systems and Engineering Immunology and Department of Immunobiology, Yale School of Medicine, New Haven, CT, USA*

[2] *Multiscale Systems Biology Section, Laboratory of Immune System Biology, NIAID, National Institutes of Health, Bethesda, MD, USA* [†]

[3] *Department of Biomedical Engineering, Yale University, New Haven, CT, USA*

[4] *Chan Zuckerberg Biohub New York, New Haven, CT, USA*

(Dated: July 17, 2025)

#### CONTENTS

I. Dynamics of CD8+ memory T cell bystander activation 1
A. CD8+ memory T cell biology 1
B. Mathematical model 2
C. Model parameters and analysis of parameter-phenotype mapping 3
D. Dynamics in the case of effector CD8+ memory T cells 5
E. Assumptions, limitations, and other comments 6

II. Dynamics of trained immunity in tissue macrophages 7
A. Tissue macrophage biology 7
B. Mathematical model 8
C. Model parameters and analysis of parameter-phenotype mapping 11
D. Assumptions, limitations, and other comments 13

III. Dynamics of central trained immunity 14
A. Hematopoiesis biology 14
B. Mathematical model 15
C. Model parameters and analysis of parameter-phenotype mapping 20
1. Modeling hematopoietic dynamics 20
2. Modeling epigenetic state dynamics during hematopoiesis 20
3. Modeling feedback from monocyte-secreted cytokines to myeloid bias during hematopoiesis 23
D. Assumptions, limitations, and other comments 25

Code and data availability 26

References 26

#### I. DYNAMICS OF CD8+ MEMORY T CELL BYSTANDER ACTIVATION

##### A. CD8+ memory T cell biology

Each CD8+ memory T ($\mathrm{T_M}$) cell is relatively short-lived, with a half-life of $\sim$ 14 days in mice [1] and $\sim$ 46 days in humans [2]. Population of these cells, and thus CD8+ T cell memory of past antigen exposure, is maintained long-term via constant homeostatic proliferation: CD8+ $\mathrm{T_M}$ cells self-renew to replace dead cells (hereafter, unless specified, $\mathrm{T_M}$ refers to CD8+ $\mathrm{T_M}$ cells). $\mathrm{T_M}$ cells rely on IL-15 signaling for homeostatic proliferation [3] and knockout or inhibition of IL-15 has been shown to compromise the durability of $\mathrm{T_M}$ cell memory [4–6]. IL-15 can be expressed by numerous cell types, both immune and non-immune, including monocytes, dendritic cells, B cells, stromal cells in

*
† Present address: Division of Cancer Treatment and Diagnosis, NCI, National Institutes of Health, Rockville, MD, USA

the bone marrow and lymph nodes, endothelial cells, fibroblasts and epithelial cells in various tissues, hair follicles, and keratinocytes [3, 7]. Instead of being secreted, IL-15 is typically presented on the cell surface (in complex with IL-15R$\alpha$) and signals *in trans*, *i.e.*, cell-cell contact is needed for IL-15 signaling (this mode of cell signaling is referred to as juxtacrine signaling [8]). While some studies have reported secreted IL-15 in the serum, generally in complex with IL-15R$\alpha$ [9–11], juxtacrine signaling is the dominant mode, at least under homeostatic conditions. Thus, $\mathrm{T_M}$ cells receive IL-15 stimulation during trafficking through tissues. Different subsets of $\mathrm{T_M}$ cells exhibit different trafficking characteristics [12]. CD8+ effector memory T cells circulate continuously between tissues via the blood, and must receive IL-15 stimulation during tissue residence (and not while they are in the blood). CD8+ central memory T cells circulate through lymph nodes (but not through non-lymphoid tissues) may receive IL-15 stimulation form IL-15-presenting stromal cells in the lymph nodes. Tissue-resident CD8+ memory T cells reside in tissues long-term and may continuously receive IL-15 stimulation.

IL-15 also functions as a "danger signal": its expression is upregulated in response to inflammatory challenges such as infection or tissue damage [3]. Bacterial and viral infections upregulate IL-15 via IFN-$\alpha$ signaling [13, 14] and via MYD88-mediated TLR signaling [15]. In addition to acute inflammation, IL-15 expression is also upregulated in chronic inflammation contexts including autoimmune conditions [7, 11, 16] and in the tumor microenvironment [17]. Given the dependence of $\mathrm{T_M}$ cell proliferation on IL-15, upregulation of IL-15 during the response to an inflammatory challenge drives the TCR-independent expansion of these cells. IL-15 stimulated $\mathrm{T_M}$ cells express increased levels of TNF-$\alpha$, perforin, granzyme, and more importantly, IFN-$\gamma$ [18], which is known to promote IL-15 presentation by various cell types [19, 20]. Thus, overall, the population dynamics of $\mathrm{T_M}$ cells involve a positive feedback loop (Fig. 1 C) with (a) IL-15 promoting the activation and expansion of $\mathrm{T_M}$ cells and (b) $\mathrm{T_M}$ cell-secreted IFN-$\gamma$ promoting the expression and presentation of IL-15. We next describe a mathematical model of the dynamics of $\mathrm{T_M}$ cells.

#### B. Mathematical model

We first consider the case of tissue-resident $\mathrm{T_M}$ cells that may not exit the tissue niche to enter the blood or lymph, or spend far more time in the tissue niche compared to the circulation. Based on the biology of $\mathrm{T_M}$ cell population dynamics (Sec. I A), we can write the following equation for $N$, the total tissue-resident $\mathrm{T_M}$ cell count in a tissue:

$$\frac{dN}{dt} = rNf_{\text{Bound}}^{\text{IL-15}} - k_d N \tag{S1}$$

Here, $r$ is the proliferation rate of cells in response to IL-15 stimulation, $f_{\text{Bound}}^{\text{IL-15}}$ is the fraction of cells receiving IL-15 stimulation at a given point in time, and $k_d$ is the death rate. Let $C_{\text{IL-15}}$ be the total availability of IL-15 in the tissue niche and $C_{\text{IL-15R}}^{0}$ be the expression of the IL-15 receptor per $\mathrm{T_M}$ cell (assuming each $\mathrm{T_M}$ cell expresses the same level of the IL-15 receptor). Then, $C_{\text{IL-15R}}^{0} \cdot N$ is the total concentration of the IL-15 receptor in the tissue. Using the quadratic reaction velocity equation which relaxes the free ligand approximation in Michaelis-Menten kinetics [21], we can write $C_{\text{IL-15R}}^{\text{Bound}}$, the steady state value for the total concentration of IL-15-bound receptors:

$$C_{\text{IL-15R}}^{\text{Bound}} = \frac{1}{2}\left((C_{\text{IL-15}} + C_{\text{IL-15R}}^{0} \cdot N + K_{\text{IL-15R}}^{\text{IL-15}}) - \sqrt{(C_{\text{IL-15}} + C_{\text{IL-15R}}^{0} \cdot N + K_{\text{IL-15R}}^{\text{IL-15}})^2 - 4C_{\text{IL-15}}C_{\text{IL-15R}}^{0} \cdot N}\right) \tag{S2}$$

Here, $K_{\text{IL-15R}}^{\text{IL-15}}$ is an effective Michaelis constant for IL-15 binding to the IL-15 receptor. Since we are assuming that the expression of the IL-15 receptor is the same for each $\mathrm{T_M}$ cell, we have

$$f_{\text{Bound}}^{\text{IL-15}} = \frac{C_{\text{IL-15R}}^{\text{Bound}}}{C_{\text{IL-15R}}^{0} \cdot N} \tag{S3}$$

At steady state, Eq. S1 gives:

$$f_{\text{Bound}}^{\text{IL-15, SS}} = \frac{k_d}{r} \tag{S4}$$

Note that the mathematical framework in Eq. S1-S3 is generic and can be used to model the population dynamics of any immune cell type that relies on signaling from a cytokine (or another signaling mechanism, *e.g.*, tonic TCR signaling in the case of CD8+ naïve T cells) for homeostatic proliferation. The identity of the cytokine driving

| Parameter | Range |
| --- | --- |
| $k_d$ | 0.032–0.1 day$^{-1}$(half-life between 7 and 21 days) |
| $\lambda_{\text{IL-15}}^{\text{IFN-}\gamma}$ | 2–12 |
| $\Theta_{\text{IL-15}}^{\text{IFN-}\gamma}$ | 4–6 times the IFN-$\gamma$ level when $\lambda_{\text{IL-15}}^{\text{IFN-}\gamma} = 1$ |
| $n_{\text{IL-15}}^{\text{IFN-}\gamma}$ | 2–8 (integer values only) |

TABLE S1. Ranges from which different model parameters were sampled for parameter-phenotype mapping in the case of $\text{T}_\text{M}$ cell bystander activation dynamics.

homeostatic proliferation and the exact mechanism of signaling are unimportant; these may be unknown. Later on, we will describe how we used the same mathematical framework to model the population dynamics of tissue macrophages and hematopoietic stem cells (see Sec. II and Sec. III).

We next consider the effect of IFN-$\gamma$ secreted by $\text{T}_\text{M}$ cells on IL-15 availability in the tissue niche. Let $C_{\text{IFN-}\gamma}^0$ be the IFN-$\gamma$ expression per $\text{T}_\text{M}$ cell and $C_{\text{IL-15}}^0$ be the IL-15 availability in the absence of IFN-$\gamma$. We use a Hill function to model the effect of IFN-$\gamma$ on IL-15 availability:

$$C_{\text{IL-15}} = C_{\text{IL-15}}^0 \mathscr{H}(C_{\text{IFN-}\gamma}^0 \cdot N, \lambda_{\text{IL-15}}^{\text{IFN-}\gamma}, \Theta_{\text{IL-15}}^{\text{IFN-}\gamma}, n_{\text{IL-15}}^{\text{IFN-}\gamma}) \tag{S5}$$

Here, $\mathscr{H}$ is the shifted Hill function:

$$\mathscr{H}(C, \lambda, \Theta, n) = \lambda + (1 - \lambda) \left( \frac{1}{1 + \left(\frac{C}{\Theta}\right)^n} \right) \tag{S6}$$

$\lambda_{\text{IL-15}}^{\text{IFN-}\gamma}$ is the maximum fold-change in the availability of IL-15 that IFN-$\gamma$ can cause. $\lambda_{\text{IL-15}}^{\text{IFN-}\gamma}$ thus determines the strength of positive feedback: $\lambda_{\text{IL-15}}^{\text{IFN-}\gamma} = 1$ indicates the absence of any feedback and $\lambda_{\text{IL-15}}^{\text{IFN-}\gamma} > 1$ indicates positive feedback. Note that we assume that IFN-$\gamma$ expression per $\text{T}_\text{M}$ cell is the same and does not change in response to IL-15 signaling. Thus, the increase in IFN-$\gamma$ level in response to IL-15 upregulation is solely due to the expanded $\text{T}_\text{M}$ cell population.

*Simulating acute inflammatory challenge*— Under homeostatic conditions $H$, the IL-15 availability in the tissue niche is $C_{\text{IL-15}}^{0,H}$. To simulate an acute inflammatory challenge $I$ between the time points $t_{\text{start}}^{\text{I}}$ and $t_{\text{end}}^{\text{I}}$, we include the inflammatory challenge-driven upregulation of IL-15:

$$C_{\text{IL-15}}^0 = \begin{cases} C_{\text{IL-15}}^{0,H} \cdot \lambda_{\text{IL-15}}^{\text{I}} & \text{for} \quad t_{\text{start}}^{\text{I}} \leq t \leq t_{\text{end}}^{\text{I}} \\ C_{\text{IL-15}}^{0,H} & \text{otherwise} \end{cases} \tag{S7}$$

Here, $\lambda_{\text{IL-15}}^{\text{I}} > 1$ is the fold-change in IL-15 availability caused by the acute inflammatory challenge. Eq. S1-S5 and Eq. S7 were used to model $\text{T}_\text{M}$ cell dynamics in the present study.

##### C. Model parameters and analysis of parameter-phenotype mapping

In Eq. S1, $r$ is the proliferation rate of $\text{T}_\text{M}$ cells once they have received the signal to proliferate from IL-15 stimulation. Since $r$ is determined by the cell cycle duration once a proliferation signal has been received, its value is unlikely to change across contexts; therefore, this parameter is fixed throughout the present study. We set $r = 1.0$ day$^{-1}$, corresponding to a population doubling time of $\sim 16$ hours. Next, in Eq. S2, we set $K_{\text{IL-15R}}^{\text{IL-15}} = 500$ and specify $C_{\text{IL-15}}$ (and $C_{\text{IL-15}}^{0,H}$) in units of $K_{\text{IL-15R}}^{\text{IL-15}}$. This choice is based on the form of Eq. S2: the exact values of $C_{\text{IL-15}}$ and $K_{\text{IL-15R}}^{\text{IL-15}}$ are unimportant; only their relative value matters. Since $C_{\text{IL-15R}}^0$ and $N$ appear together in Eq. S2 and since we are not interested in the exact value of $N$, just the nature of its dynamics, we can set $C_{\text{IL-15R}}^0 = 1$ without any loss of generality. Finally, since we are only interested in the effect of IFN-$\gamma$ on IL-15 availability and not in the IFN-$\gamma$ actual concentration in the tissue niche, we set $C_{\text{IFN-}\gamma}^0 = 1$.

The remaining parameters are the death rate $k_d$ and the three Hill function parameters $\lambda_{\text{IL-15}}^{\text{IFN-}\gamma}$, $\Theta_{\text{IL-15}}^{\text{IFN-}\gamma}$, and $n_{\text{IL-15}}^{\text{IFN-}\gamma}$. Fig. 1B and Fig. S1B-C show the behavior for the no feedback case, *i.e.*, $\lambda_{\text{IL-15}}^{\text{IFN-}\gamma} = 1$. In this scenario, values of $\Theta_{\text{IL-15}}^{\text{IFN-}\gamma}$ and $n_{\text{IL-15}}^{\text{IFN-}\gamma}$ are irrelevant (since in Eq. S6, $\mathscr{H} = 1$ if $\lambda = 1$). Fig. 1D-H and Fig. S1D-I show the behavior for a single set of Hill function parameters, chosen such that the resultant behavior is representative of the dynamics in the presence of a positive feedback loop: a supercritical pitchfork bifurcation leading to bistability [22]. In these figures, we set $\Theta_{\text{IL-15}}^{\text{IFN-}\gamma} = 36000$ and $n_{\text{IL-15}}^{\text{IFN-}\gamma} = 4$. We further set $k_d = 0.05$ day$^{-1}$, *i.e.*, a half-life of $\sim 14$ days, corresponding to the case of mouse CD8+ memory T cells. Other details regarding Fig. 1 and Fig. S1 are mentioned below:

- Fig. 1B: An acute inflammatory challenge was simulated as per Eq. S7 from $t = 0$ to $t = 21$ days and with $\lambda_{\text{IL-15}}^{\text{I}} = 4$. Half-life determines the parameter $k_d$: $k_d = \frac{\log 2}{\text{Half-life}}$. There is no feedback from IFN-$\gamma$ to IL-15 availability, *i.e.*, $\lambda_{\text{IL-15}}^{\text{IFN-}\gamma} = 1$. We set $C_{\text{IL-15}}^{0,H} = 1 \cdot K_{\text{IL-15R}}^{\text{IL-15}}$. The durability of bystander activation memory, defined as the length of the period post-challenge resolution during which the $\text{T}_{\text{M}}$ cell count is at least 10% above the count at baseline (*i.e*, before the onset of inflammation), is shown as a function of the $\text{T}_{\text{M}}$ cell half-life.

- Fig. 1D: We simulated the dynamics under homeostatic conditions (no inflammatory challenge), varying $\frac{C_{\text{IL-15}}^{0,H}}{K_{\text{IL-15R}}^{\text{IL-15}}}$ (shown on the X axis). Y axis shows the $\text{T}_{\text{M}}$ cell count at steady state.

- Fig. 1E: We simulated the dynamics under homeostatic conditions (no inflammatory challenge). Both $\frac{C_{\text{IL-15}}^{0,H}}{K_{\text{IL-15R}}^{\text{IL-15}}}$ and $\lambda_{\text{IL-15}}^{\text{IFN-}\gamma}$ were varied. For each combination of $\frac{C_{\text{IL-15}}^{0,H}}{K_{\text{IL-15R}}^{\text{IL-15}}}$ and $\lambda_{\text{IL-15}}^{\text{IFN-}\gamma}$ values, we simulated the $\text{T}_{\text{M}}$ cell dynamics starting from low $\text{T}_{\text{M}}$ cell count and from high $\text{T}_{\text{M}}$ cell count, and obtained the steady state in both cases. The two steady states thus obtained were considered distinct if the $\text{T}_{\text{M}}$ count in them differed by more than 100.

- Fig. 1F-H We set $C_{\text{IL-15}}^{0,H} = 1 \cdot K_{\text{IL-15R}}^{\text{IL-15}}$. Acute inflammatory challenge was simulated as per Eq. S7 from $t = 0$ to $t = 21$ days with $\lambda_{\text{IL-15}}^{\text{I}} = 4$. We varied $\lambda_{\text{IL-15}}^{\text{IFN-}\gamma}$ (indicated by the different colors) and plotted the $\text{T}_{\text{M}}$ cell count as a function of time.

- Fig. S1A shows $f_{\text{Bound}}^{\text{IL-15}}$ calculated as per Eq. S2 and Eq. S3. Low IL-15 availability: $\frac{C_{\text{IL-15}}^{0,H}}{K_{\text{IL-15R}}^{\text{IL-15}}} = 0.2$; medium IL-15 availability: $\frac{C_{\text{IL-15}}^{0,H}}{K_{\text{IL-15R}}^{\text{IL-15}}} = 1.0$; high IL-15 availability: $\frac{C_{\text{IL-15}}^{0,H}}{K_{\text{IL-15R}}^{\text{IL-15}}} = 2.0$.

- Fig. S1B: Same as Fig. 1B; $\text{T}_{\text{M}}$ cell count is shown as a function of time.

- Fig. S1C: Same as Fig. 1B and Fig. S1B; a second inflammatory challenge is introduced as per Eq. S7 from $t = 90$ to $t = 111$ days and with $\lambda_{\text{IL-15}}^{\text{I}} = 4$.

- Fig. S1D-F: Same as Fig. 1 E.

- Fig. S1G: Same as Fig. 1 E; Regime 1: $\lambda_{\text{IL-15}}^{\text{IFN}-\gamma} = 4$; regime 2: $\lambda_{\text{IL-15}}^{\text{IFN}-\gamma} = 8$; regime 3: $\lambda_{\text{IL-15}}^{\text{IFN}-\gamma} = 24$; regime 4: $\lambda_{\text{IL-15}}^{\text{IFN}-\gamma} = 1$

- Fig. S1H-I: Acute inflammatory challenge simulation, same as Fig. 1F-H. Bystander activation memory durability is defined as the length of the period post-challenge resolution during which the $\text{T}_{\text{M}}$ cell count is at least 10% above the count at baseline (*i.e*, before the onset of the inflammatory challenge).

*Parameter-phenotype mapping analysis*— The parameter-phenotype mapping analysis was carried out using the framework described previously [23]. Briefly, we simulated an acute inflammatory challenge for an ensemble of parameter sets sampled from within the biological range. Sampled parameters included the half-life (given by $k_d$) and the Hill function parameters $\left(\lambda_{\text{IL-15}}^{\text{IFN-}\gamma}, \Theta_{\text{IL-15}}^{\text{IFN-}\gamma}, n_{\text{IL-15}}^{\text{IFN-}\gamma}\right)$; the Hill function parameters determine the nature and strength of feedback from IFN-$\gamma$ to IL-15 availability in the tissue niche. Other parameters were fixed as per the discussion above. Ranges from which the different parameters were sampled are shown in Table S1. The range for half-life spans the half-lives of mouse NK cells ($\sim 7$ days [24]), CD8+ memory T cells ($\sim 14$ days [1]), and CD4+ memory T cells ($\sim 10$ days [1]), three cell types that can exhibit IL-15-driven homeostatic proliferation.

$\Theta_{\text{IL-15}}^{\text{IFN-}\gamma}$, the threshold parameter for the Hill function, was sampled such that under homeostatic conditions, the feedback from IFN-$\gamma$ to IL-15 is weak: $\mathscr{H}\left(C_{\text{IFN-}\gamma}^{0} \cdot N^{\text{No challenge}}, \lambda_{\text{IL-15}}^{\text{IFN-}\gamma}, \Theta_{\text{IL-15}}^{\text{IFN-}\gamma}, n_{\text{IL-15}}^{\text{IFN-}\gamma}\right) < \lambda_{\text{IL-15}}^{\text{IFN-}\gamma}/2$. Any sampled parameter set that did not satisfy this criterion was excluded from the ensemble. This ensured that none of the parameter sets in the ensemble corresponded to a scenario with persistently elevated $\text{T}_{\text{M}}$ cell counts and IFN-$\gamma$ levels even in the absence of an inflammatory challenge (*i.e.*, regime 3). To generate the ensemble of parameter sets, we used Latin hypercube sampling [25] to sample in the four-dimensional space $\left(k_d, \lambda_{\text{IL-15}}^{\text{IFN-}\gamma}, \Theta_{\text{IL-15}}^{\text{IFN-}\gamma}, n_{\text{IL-15}}^{\text{IFN-}\gamma}\right)$. Overall, we simulated an acute inflammatory challenge for a total of 45000 parameter sets and calculated the memory durability in each case. This memory durability is the phenotype of interest for parameter-phenotype mapping analysis. The sampled parameter sets and the corresponding durability values formed the basis for the analysis shown in Fig. 1I and Fig. S1J-L:

- Fig. S1J: Memory durability and frequency of occurrence of the "infinite" memory phenotype for the ensemble of parameter sets, shown as a function of the feedback strength ($\lambda_{\text{IL-15}}^{\text{IFN-}\gamma}$).
- Fig. S1K: We first defined four memory phenotypes based on the memory durability values: short-term memory ($\leq$ 90 days), medium-term memory ($>$ 90 and $\leq$ 180 days), long-term memory ($>$ 180 and $\leq$ 360 days), and very long-term memory ($>$ 360 days). The very long-term memory phenotype includes the "infinite" memory phenotype. We then trained random forest models using the MATLAB function `fitcensemble` to predict the memory phenotype for different parameter sets. Fig. S1K shows the 10-fold cross-validation model performance for different forest sizes. A model with 700 trees, indicated by the black dashed line in the panel, was used for the analysis shown in Fig. S1L and Fig. 1I (top).
- Fig. S1L: Performance of the 700-tree random forest model on the training set (80% of the overall dataset) and the test set (20% of the overall dataset).
- Fig. 1I (top): For the 700-tree model, we used the MATLAB function `predictorImportance` to calculate the importance of each parameter to predicting the memory phenotype. The normalized importance for each parameter (importance divided by the sum of the importance values across all parameters) is shown.
- Fig. 1I (bottom): Distribution of memory durability values obtained for parameter sets in the ensemble with weak feedback ($\lambda_{\text{IL-15}}^{\text{IFN-}\gamma} \leq 4$) and strong feedback ($\lambda_{\text{IL-15}}^{\text{IFN-}\gamma} > 4$). MATLAB function `ksdensity` was used to obtain a probability density estimate for each histogram; this estimate is shown by the darker curves.

##### D. Dynamics in the case of effector CD8+ memory T cells

Unlike tissue-resident $\text{T}_\text{M}$ cells which reside in a tissue long-term, effector $\text{T}_\text{M}$ cells continuously circulate between tissues via the blood. Let $N_{T_i}$ be the effector $\text{T}_\text{M}$ cell count in a tissue $T_i$ and $N_B$ the count in blood. Note that the IL-15 stimulation needed for $\text{T}_\text{M}$ cell homeostatic proliferation is only available in the tissue niche and, thus, there is no proliferation in the blood. We can write:

$$\frac{dN_{T_i}}{dt} = \overbrace{rN_{T_i} f_{\text{Bound}}^{\text{IL-15},\, i} - k_d N_{T_i}}^{\text{Proliferation and death}} + \underbrace{k_{\text{entry}}^{T_i} N_B - k_{\text{exit}}^{T_i} N_{T_i}}_{\text{Trafficking}} \tag{S8}$$

$$\frac{dN_B}{dt} = -k_d N_B - \sum_i k_{\text{entry}}^{T_i} N_B + \sum_i k_{\text{exit}}^{T_i} N_{T_i} \tag{S9}$$

Here, $k_{\text{entry}}^{T_i}$ and $k_{\text{exit}}^{T_i}$ are the rates of effector $\text{T}_\text{M}$ cell trafficking in and out of the tissue $T_i$, respectively. For each tissue, $f_{\text{Bound}}^{\text{IL-15},\, i}$ can be calculated from Eq. S2 and Eq. S3 by plugging in tissue-specific values of IL-15 availability. An acute inflammatory challenge in a specific tissue can then be simulated by transiently upregulating the IL-15 availability in that tissue.

Eq. S8-S9 describe a general framework for modeling effector $\text{T}_\text{M}$ cell dynamics. We can simplify this setup by assuming that each tissue is identical in terms of IL-15 availability and trafficking kinetics. With $N_T \equiv \sum_i N_{T_i}$ and $k_{\text{entry}}^{T_i} = k_{\text{entry}}^{T}$, $k_{\text{exit}}^{T_i} = k_{\text{exit}}^{T}$ as the tissue-independent trafficking parameters, we can write:

$$\frac{dN_T}{dt} = rN_T f_{\text{Bound}}^{\text{IL-15}} - k_d N_T + k_{\text{entry}}^{T} N_B - k_{\text{exit}}^{T} N_T \tag{S10}$$

$$\frac{dN_B}{dt} = -k_d N_B - k_{\text{entry}}^{T} N_B + k_{\text{exit}}^{T} N_T \tag{S11}$$

At steady state, Eq. S11 gives us:

$$N_B^{\text{SS}} = \frac{k_{\text{exit}}^{T} N_T^{\text{SS}}}{k_{\text{entry}}^{T} + k_d} \tag{S12}$$

Plugging into Eq. S10 at steady state:

$$\begin{aligned} f^{\text{IL-15, SS}}_{\text{Bound}} &= \left(\frac{1}{r}\right)\left(k_d - k^T_{\text{entry}}\left(\frac{k^T_{\text{exit}}}{k^T_{\text{entry}} + k_d}\right) + k^T_{\text{exit}}\right) \\ &= \left(\frac{k_d}{r}\right)\left(1 + \frac{k^T_{\text{exit}}}{k^T_{\text{entry}} + k_d}\right) \end{aligned} \tag{S13}$$

Eq. S13 has the same mathematical form as Eq. S4 with a scaled right-hand side. Thus, the steady state behavior in the case of effector $T_M$ cells will be qualitatively very similar to that in the case of tissue-resident $T_M$ cells.

##### E. Assumptions, limitations, and other comments

- In the tissue niche, tissue-resident $T_M$ cells reside long-term and compete for the limited IL-15 available. Effector $T_M$ cells traffic through a tissue surveilling for immune challenges before re-entering the blood and must receive IL-15 stimulation during their time in a tissue [12]. Here, we do not model spatial aspects of any plausible migration of tissue-resident $T_M$ cells within a tissue or of the trafficking of effector $T_M$ cells in and out of tissues. We treat the tissue niche as a well-mixed "bag" of cells presenting IL-15 or expressing the IL-15 receptor ($T_M$ cells); in fact, our modeling framework has no spatial component. Given our focus on the population dynamics of $T_M$ cells rather than their spatial distribution, this assumption is justified. Moreover, there is little data available on the spatial organization and migratory patterns of tissue-resident $T_M$ cells with a tissue or on the trafficking of effector $T_M$ cells through a tissue under homeostatic conditions. This may change with recent advances in spatial transcriptomics [26]. Lack of a spatial component makes the present framework unsuitable for modeling, in detail, the effect of chemokines on $T_M$ cell dynamics.

- The model does not track the dynamics of individual $T_M$ cells; only the overall effect of IL-15 on the population dynamics of $T_M$ cells is modeled. This choice greatly simplifies the modeling task, allowing us to write just one differential equation describing $T_M$ cell dynamics (Eq. S1) instead of having to write an equation for the signaling state of each $T_M$ cell or having to use a stochastic modeling setup which becomes very time-consuming to simulate as the number of cells grows. This modeling choice means that we cannot track an individual cell of interest, *e.g.*, a $T_M$ cell with a specific TCR.

- To incorporate the overall effect of IL-15 and IFN-$\gamma$ signaling on $T_M$ cell dynamics without modeling the corresponding signaling pathways in detail, we assume that all signaling events are much faster than cellular events such as division or death. Since signaling events in biology are expected to occur on a timescale of minutes to a few hours while cell division and death occur on a timescale of many hours to days, our assumption is justified. This assumption allows us to assume that signaling processes in the model— IL-15 binding to IL-15R, IFN-$\gamma$ production by $T_M$ cells, and change in IL-15 presentation in response to IFN-$\gamma$— are in steady state, leading to the mathematical forms in Eq. S2 and Eq. S5.

- Since our focus is on how IL-15-presenting immune and non-immune cells affect $T_M$ cell dynamics, only the overall IL-15 availability is considered in the model; the dynamics of different IL-15-presenting cell types are not modeled.

- Our model assumes that IFN-$\gamma$ expression and secretion by $T_M$ cells is unaffected by IL-15 stimulation. Thus, the increase in IFN-$\gamma$ levels upon IL-15 upregulation is due to the expansion of the $T_M$ cell population. This is a simplifying assumption and IL-15 has been shown to promote IFN-$\gamma$ secretion by $T_M$ cells [27]. The present model can be easily adapted to include an IL-15-dependence of IFN-$\gamma$ secretion by $T_M$ cells.

- We assume that all $T_M$ cells exhibit the same signaling behavior: they express same levels of the IL-15 receptor and have the same responsiveness to IL-15 signaling. All $T_M$ cells secrete the same amount of IFN-$\gamma$ in response to IL-15 signaling. This assumption can be relaxed, and the model easily extended to describe the dynamics of multiple $T_M$ subtypes that may exhibit different sensitivities to IL-15 signaling, *e.g.*, CD45RO+ $T_M$ cells which are more responsive to IL-15 versus CD45RA+ $T_M$ cells that are less responsive to IL-15 stimulation [28]. One can model such behavior by writing a separate differential equation for each $T_M$ cell subtype, each equation following the mathematical form of Eq. S1. Note that cells of all subtypes will compete for the same pool of IL-15 in the tissue niche. If cells of different subtypes have different expression levels of the IL-15 receptor or differ in their responsiveness to IL-15 (*e.g.*, different values of $K^{\text{IL-15}}_{\text{IL-15R}}$), the fraction of cells of each subtype receiving

IL-15 stimulation may no longer be analytically tractable within a Michaelis-Menten type framework. In such a scenario, alternate ways for determining the fraction receiving IL-15 stimulation will need to be developed, or a more generalized framework that does not explicitly involve receptor binding kinetics may be used.

- Our modeling setup only includes the effect of IL-15 and IFN-$\gamma$ signaling on $T_M$ population dynamics. With such a setup, we do not intend to imply that these are the only active signaling pathways between $T_M$ cells and other immune and non-immune cell types in tissues, or that signaling from IL-15 and IFN-$\gamma$ dominates $T_M$ dynamics in all contexts. Our goal here is not to develop a comprehensive mathematical model that incorporates all possible signaling pathways affecting $T_M$ cells. We focus instead on exploring the range of $T_M$ cell behaviors that IL-15 and IFN-$\gamma$ signaling can drive. Given the crucial role of IL-15 in driving $T_M$ cell homeostatic proliferation and that of IFN-$\gamma$ as a key component of the CD8+ T cell inflammatory response, our focus on these two signaling pathways is well-grounded in immunobiology. Further, activities of signaling pathways that drive functional consequences similar to that of IL-15 and IFN-$\gamma$ signaling can at least partially be recapitulated by our model. For example, the activity of a signaling pathway that lowers the overall IL-15 expression in a tissue can be modeled by lowering the maximum fold-change in IL-15 expression that can be caused by the IFN-$\gamma$ secreted by $T_M$ cells.

- $T_M$ cells are not the only immune cells that rely on IL-15 for homeostatic proliferation. Natural killer (NK) cells also rely on IL-15 for long-term maintenance in the periphery [24] and can expand as well as secrete IFN-$\gamma$ in response to increased IL-15 stimulation during an inflammatory challenge [29]. Thus, NK cell dynamics during homeostasis and in response to inflammatory challenges are expected to be mechanistically similar to $T_M$ cells— involving a positive feedback loop driven by IL-15 and IFN-$\gamma$ signaling— and NK cells are predicted to exhibit the same qualitative dynamics as $T_M$ cells. While CD4+ memory T cells generally rely on IL-7 for homeostatic proliferation, they can also express the IL-15 receptor and undergo IL-15-driven homeostatic proliferation [30], particularly when IL-7 is less abundant [31]. The inflammatory response of CD4+ memory T cells however depends on their subtype: while Th1 cells can secrete IFN-$\gamma$, Th2 and Th7 subsets do not. Thus, Th1 cell dynamics, when driven by IL-15 signaling, can be subject to positive feedback. In contrast, Th2 and Th17 cell dynamics are expected to be closer to the weak or no feedback case.

#### II. DYNAMICS OF TRAINED IMMUNITY IN TISSUE MACROPHAGES

##### A. Tissue macrophage biology

*Population dynamics*— Macrophages can be maintained in tissues long-term, without any input of monocytes generated in the bone marrow, via homeostatic proliferation driven by certain cytokines secreted by cells in the tissue niche [32, 33]. Tissue-resident macrophages express CSF-1R, the receptor for M-CSF and IL-34. M-CSF (also known as CSF-1) is constitutively expressed by a variety of cell types in the tissue niche including endothelial cells, fibroblasts, osteoblasts, smooth muscle cells, and macrophages themselves [34], and has been shown to drive the expansion of peritoneal macrophages and lung macrophages [35, 36]. IL-34, expressed by neurons in the brain and by keratinocytes in the skin, promotes the self-renewal of microglia and Langerhans cells [37, 38]. GM-CSF (also known as CSF-2) can also drive the proliferation of alveolar and peritoneal macrophages [36, 39]. While the expression of GM-CSF under homeostatic conditions is typically low, it can be produced by a variety of immune as well as non-immune cell types in response to inflammatory challenges [34]. The population dynamics of tissue-resident macrophages under homeostasis are thus mechanistically similar to that of $T_M$ cells: homeostatic proliferation driven by a cytokine, the availability of which determines the overall population size. One key difference must be noted: unlike IL-15 which is presented on the cell surface, M-CSF, IL-34, and GM-CSF are secreted into the tissue niche. In fact, M-CSF is present in the plasma at a concentration of $\sim 10$ ng$\cdot$ml$^{-1}$.

While monocyte input is not essential for the long-term maintenance of tissue macrophages, macrophages derived from the differentiation of monocytes generated in the bone marrow can form a significant part of the tissue macrophage pool. The fraction of monocyte-derived macrophages in the tissue-resident pool under homeostasis is tissue dependent and increases with age. While inflammatory challenges increase the infiltration of monocytes into the tissue niche, these monocytes contribute to the macrophage pool over the long-term only when inflammation is accompanied by the death of tissue macrophages [33, 40, 41].

*Epigenetic state dynamics*— The epigenetic state of a cell is defined by the type of histone modifications and the presence or absence of DNA methylation at various genomic loci [42]. These covalent modifications to histone tails (nucleotides in the case of DNA methylation) are well-known to modulate the transcriptional response of cells and play a key role in the establishment of cell type-specific gene expression patterns, including in the immune context [43–45]. Conversely, the epigenetic state of immune cells can change in response to transcriptional signaling [46–48].

Various inflammatory challenges have been shown to change the epigenetic state of macrophages and / or precursor monocytes. $\beta$-glucan promotes the deposition of the activating histone mark H3K4me3 over the promoter regions of multiple genes including TNF-$\alpha$, IL-6, IL-18, TLR4, and MYD88 in peritoneal macrophages [49]. Vaccination with the Bacillus Calmette-Guérin (BCG) vaccine can drive epigenetic reprogramming of bone marrow-derived macrophages, including deposition of the activating histone marks H3K27Ac and H3K4me1 in enhancer regions located in the gene body of STAT3 [50]. In muco-obstructive mice, alveolar macrophages exhibit increased chromatin accessibility at genomic regions with motifs bound by inflammatory transcription factors such as IRF1; these macrophages secrete increased amounts of IL-1$\alpha$ and IL-6 upon lipopolysaccharide exposure [51].

The aforementioned scenarios are all examples of innate immune memory termed "trained immunity" wherein an inflammatory challenge induces a reprogrammed epigenetic state, often involving deposition of activating histone marks at genomic loci associated with pro-inflammatory genes; such an epigenetic state can then drive an altered innate immune response to a subsequent inflammatory challenge [52]. The reprogrammed epigenetic state induced by an inflammatory challenge, hereafter referred to as the epigenetic memory state, can be lost during cell division. Studies have shown that while chromatin domains with repressive histone marks associated with heterochromatin are preserved during cell division, domains with activating histone marks are often lost unless persistent transcriptional signaling is present [53–55]. Transcription factor binding to enhancer regions and active transcription are essential for the re-establishment of accessible chromatin regions after cell division [56]. Since tissue macrophages undergo homeostatic proliferation, the epigenetic memory state induced by an inflammatory challenge can be lost over time.

*Positive feedback from macrophage-secreted cytokines to the reprogrammed epigenetic state*— We next note the possible interplay between the epigenetic state of tissue macrophages and their transcriptional response, and how such interplay can create a positive feedback loop. One example is that of IL-6. Tissue macrophages secrete IL-6 in response to foreign pathogens and tissue damage [57, 58]. Macrophages also express the IL-6 receptor and respond to IL-6 autocrine / paracrine signaling via the phosphorylation of STAT3 and its translocation into the nucleus; STAT3 in the nucleus can upregulate the expression of IL-6. STAT3 has also been shown to increase NF-$\kappa$B expression via the micro-RNAs miR-21 and miR-181-b [59]. NF-$\kappa$B signaling is downstream of multiple receptors involved in the macrophage inflammatory response including TLRs, IL-1, and TNF-$\alpha$ [60]. A search of the TRRUST database [61] shows that STAT3 also has multiple other pro-inflammatory targets including IFNAR1, IRF1, JUNB, and JAK2. In this context, the epigenetic memory state is expected to be characterized by open, accessible chromatin in the IL-6 enhancer region bound by STAT3 (and in the IL-6 gene body), as well as in other chromatin regions that can be bound by STAT3. Macrophages in such a state may secrete more IL-6 even in the absence of an inflammatory challenge which, in turn, can maintain the epigenetic memory state via STAT3 activation. Since STAT3 binds not just the IL-6 enhancer region but also genomic regions associated with various other inflammatory genes, macrophages in such an epigenetic state may exhibit an enhanced response to inflammatory signaling driven by factors such as IFN-$\alpha$ or any other signaling modality involving NF-$\kappa$B.

Note that this is just one example of possible positive feedback from a macrophage-secreted cytokine to the macrophage epigenetic state, and feedback involving other signaling molecules is likely to exist. Our goal here is to demonstrate that the presence of such feedback can tune the durability of trained immunity in tissue macrophages; the exact molecular players involved are unimportant for our analysis. Below, we describe a mathematical model of the population dynamics of tissue macrophages and the dynamics of the epigenetic memory state in these cells.

##### B. Mathematical model

*Tissue macrophage population dynamics*— To model the population dynamics of tissue macrophages, we use the same framework as in the case of tissue-resident $T_M$ cells (Sec. I B). We consider the scenario wherein there is no monocyte input to the tissue macrophage pool. Let $N$ be the macrophage count in the tissue niche. We write:

$$\frac{dN}{dt} = rNf_{\text{Bound}} - k_d N \tag{S14}$$

Here, $f_{\text{Bound}}$ is the fraction of tissue macrophages receiving stimulation at a given point in time from a cytokine that can drive macrophage homeostatic proliferation (*e.g.*, M-CSF, IL-34, or GM-CSF), $r$ is the rate of proliferation of cells upon receiving the proliferation signal, and $k_d$ is the death rate. $f_{\text{Bound}}$ can be calculated in the same way as we calculated $f_{\text{Bound}}^{\text{IL-15}}$ (see Eq. S2 and Eq. S3) or a different approach incorporating details of CSF-1 / 2 signaling may be adopted. Here, our focus is on the dynamics of the epigenetic memory state in tissue macrophages and we assume that the population dynamics of tissue macrophages are in steady state throughout. We will see below that in such a case, the exact mathematical form of $f_{\text{Bound}}$ is unimportant; only the following steady state relation is relevant:

$$f^{\text{SS}}_{\text{Bound}} = \frac{k_d}{r} \tag{S15}$$

**Box 1: Estimating the death rate in a population of cells undergoing homeostatic proliferation**

We note that the steady state relation in Eq. S15 will hold for any immune cell type undergoing homeostatic proliferation in response to signaling from a factor whose overall availability is limited. $r$ is determined by the cell cycle duration of cells once they have received a proliferation signal and is thus unlikely to vary significantly across contexts. Given the typical cell cycle time of mammalian cells, we can set $r = 1$ day$^{-1}$. Irrespective of the exact mathematical form of $f^{\text{SS}}_{\text{Bound}}$, it can be interpreted as the fraction of cells that are exhibiting markers of proliferation at steady state under homeostatic conditions. Commonly used markers for proliferating immune cells include Ki67 expression which is restricted to active phases of the cell cycle [62], phosphorylation of histone H3 (pHH3) which occurs during a discrete stage of mitosis [63], and nuclear DNA content. The fraction of cells in a population that are Ki67$^{+}$ or pHH3$^{+}$, or that have $>$ 2N DNA content will roughly equal $f^{\text{SS}}_{\text{Bound}}$. One can then use Eq. S15 to estimate $k_d$.

*Epigenetic state dynamics*— We use a binary variable to describe the epigenetic state of tissue macrophages: an epigenetic state of 1 indicates that the macrophage has a reprogrammed epigenetic state induced by an inflammatory challenge, *i.e.*, is in an epigenetic memory state. An epigenetic state of 0 indicates the lack of any epigenetic reprogramming. Based on the discussion in Sec. II A, we assume that the epigenetic memory state 1 can be lost during cell division with a probability $1 - p_{\text{division}}$; the epigenetic state 0 is maintained during cell division. With $g_{\text{Epi}}$ as the rate of establishment of the epigenetic memory state in tissue macrophages, we can write for $f_{\text{Epi, 1}}$, the fraction of macrophages in an epigenetic memory state:

$$\frac{df_{\text{Epi, 1}}}{dt} = rf_{\text{Bound}} \cdot f_{\text{Epi, 1}} \cdot p_{\text{division}} - k_d f_{\text{Epi, 1}} + g_{\text{Epi}}(1 - f_{\text{Epi, 1}}) \tag{S16}$$

The above equation is for the scenario with no monocyte contribution to the tissue macrophage pool. Assuming that the population dynamics of macrophages are in a steady state, we can substitute from Eq. S15 and write:

$$\frac{df_{\text{Epi, 1}}}{dt} = -k_d f_{\text{Epi, 1}}(1 - p_{\text{division}}) + g_{\text{Epi}}(1 - f_{\text{Epi, 1}}) \tag{S17}$$

For the steady state of epigenetic dynamics, we have:

$$f^{\text{SS}}_{\text{Epi, 1}} = \frac{g_{\text{Epi}}}{g_{\text{Epi}} + k_d\,(1 - p_{\text{division}})} \tag{S18}$$

Eq. S18 can be used to estimate $g_{\text{Epi}}$ from experimental data on the fraction of tissue macrophages in an epigenetic memory state under homeostatic conditions and in the absence of feedback from a macrophage-secreted cytokine to the epigenetic state. Note that if at $t = t_0$, $f_{\text{Epi, 1}} = f^0_{\text{Epi, 1}}$, then for $t > t_0$, we have:

$$f_{\text{Epi, 1}}(t) = \left(\frac{g_{\text{Epi}}}{g_{\text{Epi}} + k_d(1 - p_{\text{division}})}\right) + \left(f^0_{\text{Epi, 1}} - \left(\frac{g_{\text{Epi}}}{g_{\text{Epi}} + k_d(1 - p_{\text{division}})}\right)\right) e^{-(g_{\text{Epi}} + k_d(1 - p_{\text{division}}))(t - t_0)} \tag{S19}$$

Thus, when there is no or very weak feedback from a macrophage-secreted cytokine to the epigenetic reprogramming of tissue macrophages, the kinetics of decay of $f_{\text{Epi, 1}}(t)$ after a perturbation will be determined by an effective decay constant $k_d^{'} \equiv g_{\text{Epi}} + k_d(1 - p_{\text{division}})$.

*Modeling positive feedback from a macrophage-secreted cytokine to epigenetic reprogramming*— We next consider the possibility of interplay between a macrophage-secreted cytokine and the epigenetic state of tissue macrophages. Macrophages in an epigenetic memory state can secrete $\lambda^{\text{Epi}}_{\text{Sig}}$-fold more of a cytokine such as IL-6 than macrophages in epigenetic state 0. The concentration of IL-6 in the tissue niche can then be written as:

$$C_{\text{IL-6}} = C^0_{\text{IL-6}} \cdot N \cdot \left(\lambda^{\text{Epi}}_{\text{Sig}} \cdot f_{\text{Epi, 1}} + \left(1 - f_{\text{Epi, 1}}\right)\right) \tag{S20}$$

Here, $C^0_{\text{IL-6}}$ is the IL-6 expression per macrophage in epigenetic state 0. IL-6, in turn, can promote the establishment of the epigenetic memory state in tissue macrophages. We model this effect using a Hill function (Eq. S6):

$$g_{\text{Epi}} = g^0_{\text{Epi}} \mathscr{H}(C_{\text{IL-6}}, \lambda^{\text{Sig}}_{\text{Epi}}, \Theta^{\text{Sig}}_{\text{Epi}}, n^{\text{Sig}}_{\text{Epi}}) \tag{S21}$$

Here, $g^0_{\text{Epi}}$ is the rate of epigenetic reprogramming (or the rate of establishment of the epigenetic memory state) in the absence of IL-6 signaling. Note that $\lambda^{\text{Epi}}_{\text{Sig}}$ determines the effect of epigenetic memory on IL-6 production and $\lambda^{\text{Sig}}_{\text{Epi}}$ determines the effect of IL-6 signaling on epigenetic memory. Together, these parameters determine the overall strength of the positive feedback. If either of these parameters is set to 1, the positive feedback loop would be incomplete; $\lambda^{\text{Epi}}_{\text{Sig}}, \lambda^{\text{Sig}}_{\text{Epi}} > 1$ represents a scenario with an active positive feedback loop.

*Simulating acute inflammatory challenge*— Under homeostatic conditions $H$, the rate of epigenetic reprogramming of tissue macrophages is $g^{0,\text{ H}}_{\text{Epi}}$. To simulate an acute inflammatory challenge that induces epigenetic reprogramming of tissue macrophages between the time points $t^{\text{I}}_{\text{start}}$ and $t^{\text{I}}_{\text{end}}$, we include an inflammatory challenge-induced increase in the epigenetic reprogramming rate:

$$g^0_{\text{Epi}} = \begin{cases} g^{0,\text{ H}}_{\text{Epi}} \cdot \lambda^{\text{I}}_{\text{Epi}} & \text{for} \quad t^{\text{I}}_{\text{start}} \le t \le t^{\text{I}}_{\text{end}} \\ g^{0,\text{ H}}_{\text{Epi}} & \text{otherwise} \end{cases} \tag{S22}$$

Here, $\lambda^{\text{I}}_{\text{Epi}} > 1$ is the inflammatory challenge-induced fold-change in the rate of epigenetic reprogramming of tissue macrophages.

**Box 2: Macrophage dynamics in the presence of monocyte input**

When there is significant monocyte input into the tissue macrophage pool ($N$), we can modify Eq. S14 to write:

$$\frac{dN}{dt} = rNf_{\text{Bound}} - k_d N + k_{\text{Input}} M_{\text{Blood}} \tag{S23}$$

Here, $M_{\text{Blood}}$ is the monocyte count in the blood and the rate parameter $k_{\text{Input}}$ incorporates both monocyte entry into the tissue niche and the differentiation of monocytes into macrophages. At steady state:

$$f^{\text{SS}}_{\text{Bound}} = \frac{k_d}{r} - \frac{k_{\text{Input}}}{r}\left(\frac{M^{\text{SS}}_{\text{Blood}}}{N^{\text{SS}}}\right) \tag{S24}$$

where $M^{\text{SS}}_{\text{Blood}}$ and $N_{\text{SS}}$ are the counts at steady state.

Let $f^{\text{Blood}}_{\text{Epi, 1}}$ be the fraction of monocytes in the blood that are in an epigenetic memory state and $p_{\text{differentiation}}$ be the probability that the epigenetic memory state is preserved during the differentiation of monocytes into macrophages. We can write:

$$\frac{df_{\text{Epi, 1}}}{dt} = rf_{\text{Epi, 1}} \cdot f_{\text{Bound}} \cdot p_{\text{division}} - k_d f_{\text{Epi, 1}} + k_{\text{Input}} \cdot f^{\text{Blood}}_{\text{Epi, 1}} \cdot p_{\text{differentiation}} \left(\frac{M_{\text{Blood}}}{N}\right) \tag{S25}$$

Assuming that the population dynamics are in steady state, we can substitute from Eq. S24:

$$\frac{df_{\text{Epi, 1}}}{dt} = -k_d(1 - p_{\text{division}}) f_{\text{Epi, 1}} + k_{\text{Input}} \left(f^{\text{Blood}}_{\text{Epi, 1}} \cdot p_{\text{differentiation}} - f_{\text{Epi, 1}} \cdot p_{\text{division}}\right) \left(\frac{M^{\text{SS}}_{\text{Blood}}}{N^{\text{SS}}}\right) \tag{S26}$$

This description will be useful for the analysis in Sec. III.

##### C. Model parameters and analysis of parameter-phenotype mapping

| Macrophage subset | Experimental data point | Reference | $k_d$ estimate | Half-life |
|---|---|---|---|---|
| Alveolar macrophages | $\sim 10\%$ of cells are $\text{BrdU}^+$ 1 day after BrdU injection under homeostasis | [64] | $\sim$ 0.1 day$^{-1}$ | $\sim$ 7 days |
| Langerhans cells | $5 - 7\%$ of cells are $\text{Ki67}^+$ at homeostasis | [65] | $\sim$ 0.06 day$^{-1}$ | $\sim$ 12 days |
| Peritoneal macrophages | $\sim 5\%$ of cells are $\text{Ki67}^+$ at homeostasis | [66] | $\sim$ 0.01 day$^{-1}$ | $\sim$ 14 days |

TABLE S2. Estimates of half-lives of different mouse tissue macrophage subsets based on Eq. S15 and the rationale in Box 1.

Once again, since $r$ in Eq. S14 and Eq. S15 is determined by the cell cycle duration of cells, we set $r = 1$ day$^{-1}$ (also, see Sec. I C). Using the rationale described in Box 1, we can then estimate the death rate $k_d$ for different tissue macrophage subsets. These estimates, along with references for the data used, are shown in Table S2. Next, since $C^0_{\text{IL-6}}$ and $N$ appear together in Eq. S20 and we are not interested in the exact value of $N$, we can set $C^0_{\text{IL-6}} = 1$. The rate of establishment of the epigenetic memory state in tissue macrophages $g^0_{\text{Epi}}$ can be estimated from Eq. S18 provided the fraction of tissue macrophages that are in an epigenetic memory state at baseline, *i.e.*, in the absence of an inflammatory challenge and in the absence of any cytokine signaling that affects the epigenetic memory state, and $p_{\text{division}}$ are known. In the absence of such data, Eq. S18 can still provide an upper bound on $g^0_{\text{Epi}}$ (and $g_{\text{Epi}}$). Under homeostatic conditions, the fraction of tissue macrophages in an epigenetic memory state is not expected to be high; we can choose $g^0_{\text{Epi}}$ accordingly. Unless specified otherwise, we set $g^{0,\text{ H}}_{\text{Epi}} = g_0$ such that for alveolar macrophages with a half-life of 7 days and for $p_{\text{division}} = 0.5$, $f^{\text{SS}}_{\text{Epi, 1}} = 0.05$.

Fig. 2B and Fig. S2A-B show the behavior for the no feedback case, *i.e.*, $\lambda^{\text{Epi}}_{\text{Sig}} = \lambda^{\text{Sig}}_{\text{Epi}} = 1$. The values of $\Theta^{\text{Sig}}_{\text{Epi}}$ and $n^{\text{Sig}}_{\text{Epi}}$ are irrelevant in this case. Fig. 3B and Fig. S3A-G show the behavior for a set of values of $g^{0,\text{ H}}_{\text{Epi}}$, $\lambda^{\text{Epi}}_{\text{Sig}}$, $\lambda^{\text{Sig}}_{\text{Epi}}$, $\Theta^{\text{Sig}}_{\text{Epi}}$, and $n^{\text{Sig}}_{\text{Epi}}$ for which the model behavior is representative of the dynamics in the presence of positive feedback. In all these cases, we set $\lambda^{\text{Epi}}_{\text{Sig}} = 14$, $\Theta^{\text{Sig}}_{\text{Epi}} = 36000$, and $n^{\text{Sig}}_{\text{Epi}} = 6$, and simulate the behavior for the case of mouse alveolar macrophages with a half-life of $\sim$ 7 days ($k_d \sim 0.1$ day$^{-1}$) and for $p_{\text{division}} = 0.5$. Other details concerning Fig. 2B, Fig. 3B-C, Fig. S2A-B, and Fig. S3A-G are mentioned below:

- Fig. 2B: We set $g^{0,\text{ H}}_{\text{Epi}} = 0$ and thus $f^{\text{SS}}_{\text{Epi, 1}} = 0$. At $t = 0$, an epigenetic memory state is induced in each cell in the tissue macrophage population, *i.e.*, we set $f_{\text{Epi, 1}} = 1$. The decay of $f_{\text{Epi, 1}}$ is then tracked over time. Half-life determines $k_d$: $k_d = \frac{\log 2}{\text{Half-life}}$. The plot shows the durability of trained immunity in tissue macrophages, defined as the duration for $t > 0$ for which the fraction of macrophages in an epigenetic memory state is at least 10% higher than the fraction at $t = 0$.

- Fig. 3B (top right): We simulated the dynamics under homeostatic conditions (no inflammatory challenge), varying $\frac{g^{0,\text{ H}}_{\text{Epi}}}{g_0}$ (X axis) and $\lambda^{\text{Sig}}_{\text{Epi}}$ (Y axis). For each combination of $\frac{g^{0,\text{ H}}_{\text{Epi}}}{g_0}$ and $\lambda^{\text{Sig}}_{\text{Epi}}$ values, we simulated the dynamics starting from two different initial conditions: $f_{\text{Epi, 1}} = 0$ and $f_{\text{Epi, 1}} = 1$. The steady states obtained for the two initial conditions were considered distinct if the difference between the reprogrammed fraction in the two cases was greater than 0.01.

- Fig. 3B (top left): We simulated the dynamics under homeostatic conditions (no inflammatory challenge), varying $\frac{g^{0,\text{ H}}_{\text{Epi}}}{g_0}$ (X axis). Y axis shows $f_{\text{Epi, 1}}$ at steady state.

- Fig. 3B (bottom panels): An acute inflammatory challenge was simulated as per Eq. S22 from $t = 0$ to $t = 21$ days and with $\lambda^{\text{I}}_{\text{Epi}} = 10$. Setting $g^{0,\text{ H}}_{\text{Epi}} = g_0$, we varied $\lambda^{\text{Sig}}_{\text{Epi}}$ and plotted $f_{\text{Epi, 1}}$ as a function of time.

- Fig. S3A-D: Same as Fig. 3B (top right).

- Fig. S3E: Same as Fig. 3B (top left). Regime 1: $\lambda^{\text{Sig}}_{\text{Epi}} = 6$; regime 2: $\lambda^{\text{Sig}}_{\text{Epi}} = 12$; regime 3: $\lambda^{\text{Sig}}_{\text{Epi}} = 30$; regime 4: $\lambda^{\text{Sig}}_{\text{Epi}} = 1$.

- Fig. S3F: Same as Fig. 3B (bottom panels).

- Fig. S3G: An acute inflammatory challenge was simulated as per Eq. S22 from $t = 0$ to $t = 21$ days and with $\lambda^{\mathrm{I}}_{\mathrm{Epi}} = 10$. We varied $\frac{g^{0,\,\mathrm{H}}_{\mathrm{Epi}}}{g_0}$ (X axis) and $\lambda^{\mathrm{Sig}}_{\mathrm{Epi}}$ (Y axis) and plot the memory durability, defined as the duration for $t > 21$ for which the fraction of macrophages in an epigenetic memory state is at least 10% higher than the fraction at $t = 0$.

*Parameter-phenotype mapping analysis*— We simulated an acute inflammatory challenge as per Eq. S22 from $t = 0$ to $t = 21$ days with $\lambda^{\mathrm{I}}_{\mathrm{Epi}} = 10$ for an ensemble of parameter sets drawn from within the biological range. Sampled parameters included $k_d$, $p_{\mathrm{division}}$, $g^{0,\,\mathrm{H}}_{\mathrm{Epi}}$, $\lambda^{\mathrm{Epi}}_{\mathrm{Sig}}$, $\lambda^{\mathrm{Sig}}_{\mathrm{Epi}}$, $\Theta^{\mathrm{Sig}}_{\mathrm{Epi}}$, and $n^{\mathrm{Sig}}_{\mathrm{Epi}}$. The range of values for each of these parameters is shown in Table S3. The range for $k_d$ spans the half-lives of different tissue macrophage subsets in mice. $g^{0,\,\mathrm{H}}_{\mathrm{Epi}}$ was sampled from a range dependent on the values of $k_d$ and $p_{\mathrm{division}}$ sampled: after sampling $k_d$ and $p_{\mathrm{division}}$, we sampled $g^{0,\,\mathrm{H}}_{\mathrm{Epi}}$ such that $k_d(1 - p_{\mathrm{division}})\left(\frac{f^{\mathrm{min}}_{\mathrm{Epi},\,1}}{1-f^{\mathrm{min}}_{\mathrm{Epi},\,1}}\right) < g^{0,\,\mathrm{H}}_{\mathrm{Epi}} < k_d(1 - p_{\mathrm{division}})\left(\frac{f^{\mathrm{max}}_{\mathrm{Epi},\,1}}{1-f^{\mathrm{max}}_{\mathrm{Epi},\,1}}\right)$ where $f^{\mathrm{min}}_{\mathrm{Epi},\,1} = 0.05$ and $f^{\mathrm{max}}_{\mathrm{Epi},\,1} = 0.2$ are the minimum and maximum possible values, respectively, of $f^{\mathrm{SS}}_{\mathrm{Epi},\,1}$ at homeostasis and in the absence of feedback. This approach for sampling $g^{0,\,\mathrm{H}}_{\mathrm{Epi}}$ ensures that the values of this parameter are physiological, with only a small fraction of tissue macrophages in an epigenetic memory state under homeostasis. Alternately, one can sample $g^{0,\,\mathrm{H}}_{\mathrm{Epi}}$ independent of $k_d$ and $p_{\mathrm{division}}$ and then discard parameter sets for which too many macrophages are in an epigenetic memory state under homeostasis. $\Theta^{\mathrm{Sig}}_{\mathrm{Epi}}$ was chosen such that it is 2-3 times the IL-6 concentration at homeostasis in the absence of any feedback from IL-6 to the epigenetic reprogramming rate. This choice further ensures that $f^{\mathrm{SS}}_{\mathrm{Epi},\,1}$ is low under homeostatic conditions. Finally, any parameter sets for which the fraction of macrophages in an epigenetic memory state under homeostatic conditions exceeded $f^{\mathrm{max}}_{\mathrm{Epi},\,1}$ were discarded. Overall, we calculated the memory durability for a total of 45000 parameter sets and calculated the memory durability in each case. Memory durability was defined as the duration after inflammatory challenge resolution during which the fraction of macrophages in an epigenetic memory state was at least 10% higher than the fraction at the onset of the challenge. This memory durability is the phenotype of interest for our parameter-phenotype mapping analysis. These parameter sets and durability values formed the basis for the analysis shown in Fig. 3C and Fig. S3H-I:

- Fig. 3C (bottom panels): We define fast decay parameter sets as those for which the effective half-life of $f_{\mathrm{Epi},\,1}$, defined as $\log 2/k'_d \equiv \log 2/\left(g^{0,\,\mathrm{H}}_{\mathrm{Epi}} + k_d(1 - p_{\mathrm{division}})\right)$, is less than 14. Slow decay parameter sets are those for which the effective half-life is greater than 28 days.

- Fig. S3H: Statistics of memory durability for the parameter sets in the ensemble shown as function of $\lambda^{\mathrm{Epi}}_{\mathrm{Sig}}$ and $\lambda^{\mathrm{Epi}}_{\mathrm{Sig}}$.

- Fig. S3I (bottom panel): Distribution of durability values obtained for parameter sets in the ensemble with weak feedback ($\lambda^{\mathrm{Epi}}_{\mathrm{Sig}}, \lambda^{\mathrm{Sig}}_{\mathrm{Epi}} \leq 6$) and for parameter sets with strong feedback ($\lambda^{\mathrm{Epi}}_{\mathrm{Sig}}, \lambda^{\mathrm{Sig}}_{\mathrm{Epi}} > 6$). MATLAB function `ksdensity` was used to obtain a probability density estimate for each histogram; this estimate is shown by the darker curves.

- Fig. S3I (top left): We trained random forest models to predict the memory durability phenotype for a given parameter set. We defined four memory durability phenotypes: $\frac{\mathrm{durability}}{k'_d} \leq 4$, $4 < \frac{\mathrm{durability}}{k'_d} \leq 8$, $8 < \frac{\mathrm{durability}}{k'_d} \leq 16$, and $\frac{\mathrm{durability}}{k'_d} > 16$. Here, $k'_d = g^{0,\,\mathrm{H}}_{\mathrm{Epi}} + k_d(1 - p_{\mathrm{division}})$ and determines the effective rate of loss of epigenetic memory in the absence of feedback (Eq. S19). Instead of training the random forest model on the model parameters $k_d$ and $g^{0,\,\mathrm{H}}_{\mathrm{Epi}}$, we replace them with the composite parameter $k'_d = g^{0,\,\mathrm{H}}_{\mathrm{Epi}} + k_d(1 - p_{\mathrm{division}})$. Random forest models were trained using the MATLAB function `fitcensemble`. Top left panel in Fig. S3I shows the 10-fold cross-validation performance for different forest sizes. A model with 600 trees was used for the analysis in Fig. S3I (top right panel) and Fig. 3C (top panel).

- Fig. S3I (top right): Performance of the 600-tree random forest model on the training set (80% of the overall dataset) and the test set (20% of the overall dataset).

- Fig. 3C (top): For the 600-tree model, we used the MATLAB function `predictorImportance` to calculate the importance of each individual parameter in predicting the memory phenotype. The normalized importance for each parameter (importance divided by the sum of the importance values across all parameters) is shown.

| Parameter | Range |
|---|---|
| $k_d$ | 0.032–0.1 day$^{-1}$(half-life between 7 and 21 days) |
| $p_{\text{division}}$ | 0–0.8 |
| $g^{0,\text{ H}}_{\text{Epi}}$ | Between $k_d(1-p_{\text{division}})\left(\frac{f^{\min}_{\text{Epi, 1}}}{1-f^{\min}_{\text{Epi, 1}}}\right)$ and $k_d(1-p_{\text{division}})\left(\frac{f^{\max}_{\text{Epi, 1}}}{1-f^{\max}_{\text{Epi, 1}}}\right)$ |
| $\lambda^{\text{Epi}}_{\text{Sig}}$ | 2–16 |
| $\lambda^{\text{Sig}}_{\text{Epi}}$ | 2–16 |
| $\Theta^{\text{Sig}}_{\text{Epi}}$ | 2–3 times the IL-6 level when $\lambda^{\text{Sig}}_{\text{Epi}} = \lambda^{\text{Epi}}_{\text{Sig}} = 1$ |
| $n^{\text{Sig}}_{\text{Epi}}$ | 2–8 (integer values only) |

TABLE S3. Ranges from which different model parameters were sampled for the parameter-phenotype mapping in tissue macrophage epigenetic dynamics. Here, $f^{\min}_{\text{Epi, 1}} = 0.05$ and $f^{\max}_{\text{Epi, 1}} = 0.2$

##### D. Assumptions, limitations, and other comments

- The model of trained immunity in tissue macrophages described here relies on some of the same assumptions that were made in the case of the $\text{T}_\text{M}$ cell dynamics model (Sec. I E):
  - We model the tissue niche as a well-mixed "bag" of cells, ignoring the spatial distribution of macrophages in the niche. We do not model the dynamics of any other cell types in the tissue such as those secreting cytokines that drive the homeostatic proliferation of tissue macrophages. We additionally assume that the distribution of the cytokines(s) that drives the homeostatic proliferation of macrophages is uniform throughout the niche; this concentration does not change with time in our simulations.
  - We do not track the dynamics or the epigenetic state of individual macrophages. Only the total macrophage count and the overall fraction of macrophages in an epigenetic memory state is modeled. In doing so, we assume that all tissue macrophages exhibit the same behavior: they express the same level of the receptor for the cytokine that drives macrophage homeostatic proliferation, express the same level of IL-6 (or any other cytokine involved in the interplay between the epigenetic state and macrophage response), and exhibit the same rate of epigenetic reprogramming. Since epigenetic reprogramming typically requires the activity of one or more histone-modifying enzymes, the last assumption implies that all macrophages have similar expression levels of such enzymes.

- In using a single variable to describe the epigenetic state of tissue macrophages, we disregard any heterogeneity in cells categorized as epigenetically reprogrammed. Epigenetically reprogrammed macrophages is likely a heterogeneous group: all macrophages may not have identical histone marks at all genomic loci of interest. Our model treats all epigenetically reprogrammed cells as equivalent. This is a simplifying yet useful assumption, allowing us to track the dynamics of epigenetic memory in tissue macrophages. Our assumption is valid as long as all cells in an epigenetic memory state secrete higher amounts of a cytokine which then promotes the establishment of the epigenetic memory state. The more interesting and biologically relevant scenarios will be the ones wherein the cytokine promotes the establishment of active chromatin at multiple pro-inflammatory loci in addition to the ones associated with the cytokine gene. A more detailed model of epigenetic dynamics may incorporate multiple epigenetic state variables, each corresponding to the chromatin state at a single genomic locus of interest.

- Our choice of a single variable to model the epigenetic state instead of separate variables to track different histone marks is partly motivated by the nature of the outcome of ATAC-seq [67], an assay that is widely used to characterize the epigenetic state of cells across contexts including in trained immunity [68]. ATAC-seq characterizes the accessibility of genomic loci, an aggregate readout that may incorporate information concerning the presence and absence of many histone marks as well as DNA methylation. Further, our choice of a binary valued variable to describe the epigenetic state of tissue macrophages, while further simplifying, is compatible with the typical approach for ATAC-seq data analysis which involves identifying peaks in the ATAC-seq signal, discretizing the continuous ATAC-seq signal into a binary outcome: peak and non-peak.

- Note that activating histone marks associated with a reprogrammed epigenetic state can be removed by histone eraser enzymes, *e.g.*, JARID1 family proteins which can remove the H3K4me3 mark [69] and HDAC enzymes that erase histone acetylation [70]. Our model does not incorporate the loss of epigenetic memory due to the activity of such enzymes, thereby assuming that such loss happens over a timescale much longer than the lifespan of individual tissue macrophages.

- In the present study, we have used the interplay between IL-6 signaling and epigenetic reprogramming of tissue macrophages as a representative example to explore the effect of positive feedback on the durability of trained immunity in tissue macrophages. The proposed framework will be applicable to any scenario involving positive feedback from a macrophage-secreted cytokine to the epigenetic state of tissue macrophages. Moreover, the proposed modeling framework is not limited to tissue macrophages: it can be used to describe the epigenetic memory dynamics for any cell type that undergoes homeostatic proliferation in response to a proliferation signal whose availability is limited.

- While modeling the epigenetic state dynamics of tissue macrophages, we have assumed that the population dynamics are in steady state throughout, including during acute inflammatory challenges. This is generally not the case. Acute response to inflammatory challenges is accompanied by the increased recruitment of monocytes to the tissue niche in response to pro-inflammatory signaling including that from tissue macrophages. Tissue macrophage populations have been shown to contract in the face of inflammatory challenges due to cell death, followed by expansion to reconstitute the macrophage pool once the inflammation has resolved [33]. In some cases, the tissue macrophage pool can expand to respond to inflammatory challenges, particularly those involving IL-4 signaling [71]. We made the modeling choice to keep the population dynamics in steady state since our focus here is on the dynamics of trained immunity in tissue macrophages, and not on how their population changes in response to inflammatory challenges. The model assumption of steady state population dynamics can easily be relaxed within the current framework: instead of relying on Eq. S17 to model the epigenetic dynamics, one can use Eq. S14 and Eq. S16 together to simultaneously track the population size and epigenetic state of tissue macrophages.

- Another facet of epigenetic reprogramming of macrophages and other myeloid cells is immune tolerance wherein activating histone marks are depleted from genomic loci associated with pro-inflammatory genes [52, 72] in response to certain inflammatory challenges. The model of trained immunity dynamics described here can also be used to describe the behavior specific to immune tolerance. Specifically, Eq. S16 can be modified to include a term corresponding to the establishment of less accessible chromatin at loci associated with pro-inflammatory genes in tissue macrophages. Note that as compared to activating histone marks, repressive histone marks and DNA methylation have been shown to be transmitted from parent cells to daughter cells with higher fidelity [53–55]. This effect could drive differences between the kinetics of trained immunity and immune tolerance.

#### III. DYNAMICS OF CENTRAL TRAINED IMMUNITY

##### A. Hematopoiesis biology

*Population dynamics during hematopoiesis*— Nearly all immune cell types ultimately derive from hematopoietic stem cells (HSCs) in the adult bone marrow. HSCs are multipotent stem cells and are capable of long-term self-renewal. During hematopoiesis, this self-renewal capability is progressively lost and multipotency restricted [73]. The part of the hematopoietic process that leads to the production of the various myeloid cell types including monocytes, granulocytes, and dendritic cells is termed myelopoiesis [74]. Myelopoiesis involves a cascade of differentiation events. HSCs differentiate into short-term HSCs (ST-HSCs) which have lower self-renewal capability than HSCs. ST-HSCs differentiate into multipotent progenitors (MPPs). MPPs give rise to common myeloid progenitors (CMPs) and common lymphoid progenitors (CLPs), both of which are oligopotent. CMPs then give rise to lineage-commited progenitors of effector myeloid cell types: megakaryocyte / erythrocyte progenitors (MEPs) which ultimately give rise to platelets and red blood cells, and granulocyte / monocyte progenitors (GMPs) which ultimately give rise to granulocytes including neutrophils, eosinophils, and basophils, and to monocytes (see Figure 1 in [73]). Note that this is a simplified, more classical description of myelopoiesis. Each of the progenitor populations mentioned above has been shown to exhibit heterogeneity in gene expression, self-renewal capability, and differentiation behavior. Each progenitor may thus be further subdivided into subtypes. More recently, single-cell transcriptomics has begun to blur the boundaries between the different cell types in the hematopoietic hierarchy [75]. However, for the purpose of our analysis here, we will stick with the classical, simplified picture (see Sec. III D for further discussion).

In the bone marrow niche, the dynamics of hematopoietic stem and progenitor cells (HSPCs)— their proliferation, differentiation, and migration— are regulated by signaling from a variety of cell types including mesenchymal stem cells, perivascular cells, endothelial cells, sympathetic nerve cells, adipocytes, and osteolineage cells such as osteoblasts and osteolclasts [76, 77]. While numerous signaling molecules have been implicated in the regulation of hematopoiesis by the bone marrow niche, two factors have been shown to be crucial: SCF is needed for the self-renewal and long-term maintenance of HSCs [78, 79] while CXCL12 plays a key role in the retention of HSCs and other progenitors in the bone marrow niche [80–82]. Spatial organization of different cell types in the bone marrow niche and their respective roles in regulating hematopoiesis *in vivo* are still being deciphered with advances in imaging and lineage-tracing techniques, and recent studies point towards a large and complex set of intertwined signaling processes regulating hematopoiesis [76, 77].

*HSPCs in trained immunity*— Epigenetic reprogramming of HSPCs by inflammatory challenges has been proposed as a possible explanation for how very short-lived monocytes can exhibit long-lasting innate immune memory [52]. Multiple studies have shown that inflammatory challenges can reprogram HSPCs in the bone marrow. BCG vaccination in mice can alter the transcriptional state of HSCs and MPPs, promoting myelopoiesis. Such BCG-reprogrammed HSPCs generate monocytes and macrophages with an altered epigenetic state, including with epigenetic changes at the STAT3 locus, capable of providing enhanced protection against pulmonary *Mycobacterium tuberculosis* infection [50]. BCG vaccination in humans has also been shown to both alter the transcriptional state of HSPCs and promote the generation of monocytes with increased chromatin accessibility at genomic loci associated with pro-inflammatory genes [83]. In a mouse model, ischemic stroke was shown to induce epigenetic changes in both HSPCs and the generated monocytes; treatment of HSPCs with recombinant IL-1$\beta$ recapitulated the effect of stroke [84]. IL-1$\beta$ was also shown to reprogram HSPCs in a mouse model of ligature-induced periodontitis [85].

HSPCs can respond to inflammatory challenges both directly and indirectly. They are known to express pattern recognition receptors such as TLRs, and MYD88-mediated signaling via TLR2, TLR4, and TLR9 in HSPCs has been shown to promote myelopoiesis [86, 87]. Chronic lipopolysaccharide exposure has been shown to promote HSC expansion and to drive myelopoiesis at the expense of lymphopoiesis in both mouse and human HSCs via TLR4 activation [88]. HSPCs express receptors for IL-1$\alpha$ / $\beta$, IL-3, IL-6, G-CSF, and TNF-$\alpha$ [85, 89, 90] and can directly respond to IFN-$\alpha$ / $\beta$ [91] as well as to IFN-$\gamma$ [92]. The mechanism for long-lasting central trained immunity proposed here relies on the ability of HSPCs to respond to signaling via these receptors: the epigenetic state of HSPCs may change in response to cytokines secreted by effector immune cells, and since epigenetic reprogramming of HSPCs can alter the hematopoietic process downstream, HSPCs can participate in feedback loops involving epigenetic reprogramming and immune cell-secreted cytokines. We explore the possible consequences of such feedback.

*Positive feedback from effector myeloid cell-secreted cytokines to the epigenetic reprogramming of HSPCs*— As noted above, expression of cytokine receptors by HSPCs can allow for feedback from effector immune cells to hematopoiesis. Below, we discuss two such examples.

- HSCs and MPPs express IL1R1, the receptor for IL-1$\alpha$ / $\beta$ [85, 89] and can thus respond to IL-1$\beta$ secreted by effector myeloid cells such as monocytes and neutrophils. IL-1$\beta$ treatment has been shown to change the epigenetic state of HSPCs, creating more accessible chromatin at loci associated with pro-inflammatory genes including NFKB2, RELA, BATF, and ELK1 [84]. HSPCs in such an epigenetic state generate monocytes with an altered epigenetic state that then exhibit enhanced IL-1$\beta$ production [85]. Thus, a positive feedback loop is formed (see Fig. 3D).

- IL-1$\beta$ [93, 94], TNF-$\alpha$ [90, 95], and M-CSF [96] have all been shown to upregulate the expression of PU.1 in HSCs. PU.1 expression promotes myeloid bias in the hematopoietic process, driving increased production of effector myeloid cells. STAT3, which acts downstream of IL-1$\beta$ receptor signaling, is one of the key transcription factors shown to be associated with myeloid bias in old age [97]. Increased production of effector myeloid cells is likely to be accompanied by the increased production of the very cytokines that promote myeloid bias— IL-1$\beta$, TNF-$\alpha$, and M-CSF— once again creating a positive feedback loop (Fig. 3D).

These are two examples of possible feedback from the effector immune cells to the hematopoietic process. Given that HSPCs express receptors for various other cytokines, other feedback mechanisms likely exist. Here, we developed mathematical models of the two abovementioned positive feedback scenarios and explored the possible effects of feedback on the durability of central trained immunity.

#### B. Mathematical model

Busch *et al.* [98] devised a mouse model that permits inducible genetic labeling of HSCs and, together with data-driven mathematical modeling, allows for the estimation of the rates of self-renewal and differentiation of HSPCs under homeostatic conditions. Here, we adapted the mathematical model described by Busch *et al.* to model the dynamics of the epigenetic memory state during hematopoiesis.

*Population dynamics*— We model the population dynamics of the hematopoietic cell types encountered in the differentiation cascade from HSCs to monocytes in mice. These include HSCs, short-term HSCs (ST-HSCs), MPPs, CMPs, GMPs, and, finally, monocytes. Let $N_{\text{HSC}}$, $N_{\text{ST-HSC}}$, $N_{\text{MPP}}$, $N_{\text{CMP}}$, and $N_{\text{GMP}}$ be the counts of the respective cell types in the bone marrow niche. Following [98], we write:

$$\frac{dN_{\text{HSC}}}{dt} = r_{\text{HSC}} N_{\text{HSC}} f^{\text{HSC}}_{\text{Bound}} - \left(c_{\text{HSC}\to\text{ST-HSC}} + k_{\text{HSC}}\right) N_{\text{HSC}} \tag{S27}$$

$$\frac{dN_{\text{ST-HSC}}}{dt} = r_{\text{ST-HSC}} N_{\text{ST-HSC}} - \left(c_{\text{ST-HSC}\to\text{MPP}} + k_{\text{ST-HSC}}\right) N_{\text{ST-HSC}} + c_{\text{HSC}\to\text{ST-HSC}} N_{\text{HSC}} \tag{S28}$$

$$\frac{dN_{\text{MPP}}}{dt} = r_{\text{MPP}} N_{\text{MPP}} - \left(c_{\text{MPP}\to\text{CMP}} + c_{\text{MPP}\to\text{CLP}} + k_{\text{MPP}}\right) N_{\text{MPP}} + c_{\text{ST-HSC}\to\text{MPP}} N_{\text{ST-HSC}} \tag{S29}$$

$$\frac{dN_{\text{CMP}}}{dt} = r_{\text{CMP}} N_{\text{CMP}} - \left(c_{\text{CMP}\to\text{GMP}} + c_{\text{CMP}\to\text{MEP}} + k_{\text{CMP}}\right) N_{\text{CMP}} + c_{\text{MPP}\to\text{CMP}} N_{\text{MPP}} \tag{S30}$$

$$\frac{dN_{\text{GMP}}}{dt} = r_{\text{GMP}} N_{\text{GMP}} - \left(c_{\text{GMP}\to\text{Monocyte}} + c_{\text{GMP}\to\text{Granulocyte}} + k_{\text{GMP}}\right) N_{\text{GMP}} + c_{\text{CMP}\to\text{GMP}} N_{\text{CMP}} \tag{S31}$$

Here, $r_{\text{i}}$ and $k_{\text{i}}$ are the self-renewal and death rates, respectively, for cell type $i$. $c_{\text{i}\to\text{j}}$ is the rate of differentiation of cell type $i$ into cell type $j$. We note the following:

- For each cell type, the dynamics involve the following processes: self-renewal, death, conversion into a more differentiated cell type downstream in the hematopoietic hierarchy, and influx from the differentiation of a cell type upstream in the hematopoietic hierarchy. For example, for the case of MPPs (Eq. S29), we have:

$$\frac{dN_{\text{MPP}}}{dt} = \underbrace{r_{\text{MPP}} N_{\text{MPP}}}_{\text{Self-renewal}} - \left( \overbrace{c_{\text{MPP}\to\text{CMP}} + c_{\text{MPP}\to\text{CLP}}}^{\substack{\text{Differentiation of MPPs}\\ \text{into CMPs and CLPs}}} + \underbrace{k_{\text{MPP}}}_{\text{Death}} \right) N_{\text{MPP}} + \underbrace{c_{\text{ST-HSC}\to\text{MPP}} N_{\text{ST-HSC}}}_{\substack{\text{Differentiation of}\\ \text{ST-HSCs into MPPs}}} \tag{S32}$$

- In Eq. S27, $f^{\text{HSC}}_{\text{Bound}}$ is the fraction of HSCs that are receiving proliferation signaling (*e.g.*, from SCF-expressing cells in the bone marrow niche) at any given time point. Thus, we model the HSC population size as being limited by competition for proliferation signaling. In such a scenario, HSC self-renewal at steady state will be determined by the HSC death rate, and more importantly, by the rate of their differentiation into ST-HSCs. Thus, the HSC pool in our setup is dynamic, capable of enhanced self-renewal in response to changes in downstream dynamics. Note that the model in [98] models the self-renewal rate of all HSPCs as linear in the population size. While a setup with a linear HSC self-renewal term is sufficient to model homeostatic dynamics with the appropriate choice of model parameters for a given experimental setup, it is unsuitable for modeling the dynamics under non-homeostatic conditions since even a small change in the differentiation rate of HSCs could lead to either exponential growth in the population size or extinction of the HSC population. This led us to modify the equation for HSC dynamics from [98].
- For ST-HSCs, MPPs, CMPs, and GMPs, the self-renewal rate is a linear function of the respective cell counts (same as in [98]). This setup could still correspond to a scenario wherein the self-renewal of HSPCs is limited by the availability of a proliferation factor, provided the availability of the proliferation signal is very high or, equivalently, the cell count is much smaller than the maximum cell count that can be supported by the proliferation signal availability. Our key assumption here is that the counts of these cell types downstream from HSCs are limited by various differentiation rates rather than by competition for a proliferation signal.

GMPs finally give rise to monocytes and various granulocytes including eosinophils, basophils, and neutrophils. For monocytes, we can write:

$$\frac{dN_{\text{Monocyte}}}{dt} = c_{\text{GMP}\to\text{Monocyte}} N_{\text{GMP}} - \left(k_{\text{Monocyte}} + k^{\text{Monocyte}}_{\text{exit}}\right) N_{\text{Monocyte}} \tag{S33}$$

Here, $k^{\text{Monocyte}}_{\text{exit}}$ is the rate of exit of monocytes from the bone marrow niche. Note that $c_{\text{GMP}\to\text{Monocyte}}$ is the overall rate of generation of monocytes from GMPs and we do not model the dynamics of intermediate cell types (*e.g.*, macrophage and DC progenitors, and common monocyte progenitors) on the route from GMPs to monocytes.

*Epigenetic state dynamics*— Following the approach in Sec. II B, we used a binary variable to describe the epigenetic state of different cell types in the hematopoietic hierarchy: an epigenetic state of 1 represents a reprogrammed epigenetic state (*i.e.*, cells in an epigenetic memory state) while an epigenetic state of 0 indicates no epigenetic reprogramming. Note that in addition to loss of the epigenetic memory state during self-renewal as in the case of tissue macrophages, epigenetic memory may also be lost during cell differentiation in the hematopoietic process. We assume that the epigenetic memory state is preserved during self-renewal with probability $p_{\text{division}}$ and during cell differentiation with probability $p_{\text{differentiation}}$. Let $f^{\text{i}}_{\text{Epi, 1}}$ be the fraction of cells of type $i$ that are in an epigenetic memory state. From Eq. S27-S33, we can write:

$$\begin{aligned}\frac{df^{\text{HSC}}_{\text{Epi, 1}}}{dt} &= r_{\text{HSC}} \cdot f^{\text{HSC}}_{\text{Bound}} \cdot f^{\text{HSC}}_{\text{Epi, 1}} \cdot p_{\text{division}} - \left(c_{\text{HSC}\to\text{ST-HSC}} + k_{\text{HSC}}\right) f^{\text{HSC}}_{\text{Epi, 1}} \\ &\quad + g^{\text{HSC}}_{\text{Epi}} \left(1 - f^{\text{HSC}}_{\text{Epi, 1}}\right)\end{aligned} \tag{S34}$$

$$\begin{aligned}\frac{df^{\text{ST-HSC}}_{\text{Epi, 1}}}{dt} &= r_{\text{ST-HSC}} \cdot f^{\text{ST-HSC}}_{\text{Epi, 1}} \cdot p_{\text{division}} - \left(c_{\text{ST-HSC}\to\text{MPP}} + k_{\text{ST-HSC}}\right) f^{\text{ST-HSC}}_{\text{Epi, 1}} \\ &\quad + c_{\text{HSC}\to\text{ST-HSC}} \left(\frac{N_{\text{HSC}}}{N_{\text{ST-HSC}}}\right) \cdot f^{\text{HSC}}_{\text{Epi, 1}} \cdot p_{\text{differentiation}} + g^{\text{ST-HSC}}_{\text{Epi}} \left(1 - f^{\text{ST-HSC}}_{\text{Epi, 1}}\right)\end{aligned} \tag{S35}$$

$$\begin{aligned}\frac{df^{\text{MPP}}_{\text{Epi, 1}}}{dt} &= r_{\text{MPP}} \cdot f^{\text{MPP}}_{\text{Epi, 1}} \cdot p_{\text{division}} - \left(c_{\text{MPP}\to\text{CMP}} + c_{\text{MPP}\to\text{CLP}} + k_{\text{MPP}}\right) f^{\text{MPP}}_{\text{Epi, 1}} \\ &\quad + c_{\text{ST-HSC}\to\text{MPP}} \left(\frac{N_{\text{ST-HSC}}}{N_{\text{MPP}}}\right) \cdot f^{\text{ST-HSC}}_{\text{Epi, 1}} \cdot p_{\text{differentiation}} + g^{\text{MPP}}_{\text{Epi}} \left(1 - f^{\text{MPP}}_{\text{Epi, 1}}\right)\end{aligned} \tag{S36}$$

$$\begin{aligned}\frac{df^{\text{CMP}}_{\text{Epi, 1}}}{dt} &= r_{\text{CMP}} \cdot f^{\text{CMP}}_{\text{Epi, 1}} \cdot p_{\text{division}} - \left(c_{\text{CMP}\to\text{GMP}} + c_{\text{CMP}\to\text{MEP}} + k_{\text{CMP}}\right) f^{\text{CMP}}_{\text{Epi, 1}} \\ &\quad + c_{\text{MPP}\to\text{CMP}} \left(\frac{N_{\text{MPP}}}{N_{\text{CMP}}}\right) \cdot f^{\text{MPP}}_{\text{Epi, 1}} \cdot p_{\text{differentiation}} + g^{\text{CMP}}_{\text{Epi}} \left(1 - f^{\text{CMP}}_{\text{Epi, 1}}\right)\end{aligned} \tag{S37}$$

$$\begin{aligned}\frac{df^{\text{GMP}}_{\text{Epi, 1}}}{dt} &= r_{\text{GMP}} \cdot f^{\text{GMP}}_{\text{Epi, 1}} \cdot p_{\text{division}} - \left(c_{\text{GMP}\to\text{Monocyte}} + c_{\text{GMP}\to\text{Granulocyte}} + k_{\text{GMP}}\right) f^{\text{GMP}}_{\text{Epi, 1}} \\ &\quad + c_{\text{CMP}\to\text{GMP}} \left(\frac{N_{\text{CMP}}}{N_{\text{GMP}}}\right) \cdot f^{\text{CMP}}_{\text{Epi, 1}} \cdot p_{\text{differentiation}} + g^{\text{GMP}}_{\text{Epi}} \left(1 - f^{\text{GMP}}_{\text{Epi, 1}}\right)\end{aligned} \tag{S38}$$

$$\begin{aligned}\frac{df^{\text{Monocyte}}_{\text{Epi, 1}}}{dt} &= c_{\text{GMP}\to\text{Monocyte}} \left(\frac{N_{\text{GMP}}}{N_{\text{Monocyte}}}\right) \cdot f^{\text{GMP}}_{\text{Epi, 1}} \cdot p_{\text{differentiation}} \\ &\quad - \left(k_{\text{Monocyte}} + k^{\text{Monocyte}}_{\text{exit}}\right) f^{\text{Monocyte}}_{\text{Epi, 1}} + g^{\text{Monocyte}}_{\text{Epi}} \left(1 - f^{\text{Monocyte}}_{\text{Epi, 1}}\right)\end{aligned} \tag{S39}$$

Here, $g^{\text{i}}_{\text{Epi}}$ is the rate of epigenetic reprogramming for cell type $i$. Once again, given our focus on epigenetic memory state dynamics rather than cell count dynamics, we assume that the cell count dynamics described by Eq. S27-S33 are in a steady state throughout. Note that the mathematical form for HSC epigenetic state dynamics is the same as in the case of cycling tissue macrophages (compare Eq. S16 and Eq. S34) and that the mathematical form for epigenetic state dynamics for other cell subsets is the same as in the case of cycling macrophages with monocyte input (see Box 2 and compare Eq. S25 with Eq. S35-S38). Assuming that the population dynamics of each HSPC type are in steady state, we can write:

$$\frac{df^{\text{HSC}}_{\text{Epi, 1}}}{dt} = -\left(c_{\text{HSC}\to\text{ST-HSC}} + k_{\text{HSC}}\right)\left(1 - p_{\text{division}}\right) f^{\text{HSC}}_{\text{Epi, 1}} + g^{\text{HSC}}_{\text{Epi}} \left(1 - f^{\text{HSC}}_{\text{Epi, 1}}\right) \tag{S40}$$

$$\begin{aligned}\frac{df^{\text{ST-HSC}}_{\text{Epi, 1}}}{dt} &= -\left(c_{\text{ST-HSC}\to\text{MPP}} + k_{\text{ST-HSC}}\right)\left(1 - p_{\text{division}}\right) f^{\text{ST-HSC}}_{\text{Epi, 1}} \\ &\quad - c_{\text{HSC}\to\text{ST-HSC}} \left(\frac{N^{\text{SS}}_{\text{HSC}}}{N^{\text{SS}}_{\text{ST-HSC}}}\right) \left(f^{\text{ST-HSC}}_{\text{Epi, 1}} \cdot p_{\text{division}} - f^{\text{HSC}}_{\text{Epi, 1}} \cdot p_{\text{differentiation}}\right) \\ &\quad + g^{\text{ST-HSC}}_{\text{Epi}} \left(1 - f^{\text{ST-HSC}}_{\text{Epi, 1}}\right)\end{aligned} \tag{S41}$$

$$\begin{aligned}\frac{df_{\text{Epi, 1}}^{\text{MPP}}}{dt} =& -\left(c_{\text{MPP}\to\text{CMP}} + c_{\text{MPP}\to\text{CLP}} + k_{\text{MPP}}\right)\left(1 - p_{\text{division}}\right) f_{\text{Epi, 1}}^{\text{MPP}} \\ & - c_{\text{ST-HSC}\to\text{MPP}} \left(\frac{N_{\text{ST-HSC}}^{\text{SS}}}{N_{\text{MPP}}^{\text{SS}}}\right) \left(f_{\text{Epi, 1}}^{\text{MPP}} \cdot p_{\text{division}} - f_{\text{Epi, 1}}^{\text{ST-HSC}} \cdot p_{\text{differentiation}}\right) \\ & + g_{\text{Epi}}^{\text{MPP}} \left(1 - f_{\text{Epi, 1}}^{\text{MPP}}\right)\end{aligned} \tag{S42}$$

$$\begin{aligned}\frac{df_{\text{Epi, 1}}^{\text{CMP}}}{dt} =& -\left(c_{\text{CMP}\to\text{GMP}} + c_{\text{CMP}\to\text{MEP}} + k_{\text{CMP}}\right)\left(1 - p_{\text{division}}\right) f_{\text{Epi, 1}}^{\text{CMP}} \\ & - c_{\text{MPP}\to\text{CMP}} \left(\frac{N_{\text{MPP}}^{\text{SS}}}{N_{\text{CMP}}^{\text{SS}}}\right) \left(f_{\text{Epi, 1}}^{\text{CMP}} \cdot p_{\text{division}} - f_{\text{Epi, 1}}^{\text{MPP}} \cdot p_{\text{differentiation}}\right) \\ & + g_{\text{Epi}}^{\text{CMP}} \left(1 - f_{\text{Epi, 1}}^{\text{CMP}}\right)\end{aligned} \tag{S43}$$

$$\begin{aligned}\frac{df_{\text{Epi, 1}}^{\text{GMP}}}{dt} =& -\left(c_{\text{GMP}\to\text{Monocyte}} + c_{\text{GMP}\to\text{Granulocyte}} + k_{\text{GMP}}\right)\left(1 - p_{\text{division}}\right) f_{\text{Epi, 1}}^{\text{GMP}} \\ & - c_{\text{CMP}\to\text{GMP}} \left(\frac{N_{\text{CMP}}^{\text{SS}}}{N_{\text{GMP}}^{\text{SS}}}\right) \left(f_{\text{Epi, 1}}^{\text{GMP}} \cdot p_{\text{division}} - f_{\text{Epi, 1}}^{\text{Monocyte}} \cdot p_{\text{differentiation}}\right) \\ & + g_{\text{Epi}}^{\text{GMP}} \left(1 - f_{\text{Epi, 1}}^{\text{GMP}}\right)\end{aligned} \tag{S44}$$

For monocytes, evaluating Eq. S33 at steady state and substituting in Eq. S39, we have:

$$\frac{df_{\text{Epi, 1}}^{\text{Monocyte}}}{dt} = -\left(k_{\text{Monocyte}} + k_{\text{exit}}^{\text{Monocyte}}\right)\left(f_{\text{Epi, 1}}^{\text{Monocyte}} - f_{\text{Epi, 1}}^{\text{GMP}} \cdot p_{\text{differentiation}}\right) + g_{\text{Epi}}^{\text{Monocyte}}\left(1 - f_{\text{Epi, 1}}^{\text{Monocyte}}\right) \tag{S45}$$

Eq. S40-S45 show that the dynamics of the epigenetic memory state in HSPCs is determined by:

1. Rates of differentiation and death for different cell types
2. Ratios of counts of cell types at steady state
3. Fidelity with which the epigenetic memory state is transmitted during self-renewal and differentiation, quantified by the parameters $p_{\text{division}}$ and $p_{\text{differentiation}}$, respectively

*Modeling positive feedback from monocyte-secreted cytokines to the epigenetic reprogramming of HSPCs*— We consider that monocytes in epigenetic state 1 secrete $\lambda_{\text{Sig}}^{\text{Epi}}$-fold more of a cytokine such as IL-1$\beta$ than monocytes in epigenetic state 0. Then, the concentration of IL-1$\beta$ as seen by HSPCs in the bone marrow niche may be written as:

$$C_{\text{IL-1}\beta} = C_{\text{IL-1}\beta}^{0} \cdot N_{\text{Monocyte}} \cdot \left(\lambda_{\text{Sig}}^{\text{Epi}} \cdot f_{\text{Epi, 1}}^{\text{Monocyte}} + \left(1 - f_{\text{Epi, 1}}^{\text{Monocyte}}\right)\right) \tag{S46}$$

Here, $C_{\text{IL-1}\beta}^{0}$ is the IL-1$\beta$ expression per monocyte in epigenetic state 0. The rate of establishment of epigenetic state 1 in one or more HSPC subtypes can increase in response to IL-1$\beta$ signaling. We model this effect using a shifted Hill function (Eq. S6):

$$g_{\text{Epi}}^{\text{i}} = g_{\text{Epi}}^{\text{i, 0}} \mathscr{H}\left(C_{\text{IL-1}\beta}, \lambda_{\text{Epi}}^{\text{Sig}}, \Theta_{\text{Epi}}^{\text{Sig}}, n_{\text{Epi}}^{\text{Sig}}\right) \tag{S47}$$

Cell type $i$ is any hematopoietic cell type whose epigenetic reprogramming rate can change in response to IL-1$\beta$ signaling. $g_{\text{Epi}}^{\text{i, 0}}$ is the epigenetic reprogramming rate for cell type $i$ in the absence of IL-1$\beta$ signaling. The parameters $\lambda_{\text{Sig}}^{\text{Epi}}$ and $\lambda_{\text{Epi}}^{\text{Sig}}$ together determine the strength of the positive feedback, and the feedback loop is inactive if either of these parameters is set to 1. $\lambda_{\text{Sig}}^{\text{Epi}}, \lambda_{\text{Epi}}^{\text{Sig}} > 1$ indicates an active positive feedback loop.

*Simulating acute inflammatory challenge*— Under homeostatic conditions $H$, the rate of epigenetic reprogramming of cell type $i$ is $g_{\text{Epi}}^{\text{i, 0, H}}$. To simulate an acute inflammatory challenge that causes the epigenetic reprogramming of cell type $i$ between the time points $t_{\text{start}}^{\text{I}}$ and $t_{\text{end}}^{\text{I}}$, we include a challenge-induced increase in the epigenetic reprogramming rate:

$$g_{\text{Epi}}^{\text{i, 0}} = \begin{cases} g_{\text{Epi}}^{\text{i, 0, H}} \cdot \lambda_{\text{Epi}}^{\text{I}} & \text{for} \quad t_{\text{start}}^{\text{I}} \leq t \leq t_{\text{end}}^{\text{I}} \\ g_{\text{Epi}}^{\text{i, 0, H}} & \text{otherwise} \end{cases} \tag{S48}$$

$\lambda^{\mathrm{I}}_{\mathrm{Epi}} > 1$ is the fold-change in the epigenetic reprogramming rate of one or more HSPC subtypes induced by the acute inflammatory challenge.

*Modeling the effect of monocyte-secreted cytokines on myeloid versus lymphoid bias in hematopoiesis*— We consider myeloid bias at the MPP stage as follows: MPPs can exist in one of two states, 0 or 1. MPPs in state 0 are lymphoid-biased and differentiate into CLPs while those in state 1 are myeloid-biased give rise to CMPs. We modify Eq. S30:

$$\frac{dN_{\mathrm{CMP}}}{dt} = r_{\mathrm{CMP}} N_{\mathrm{CMP}} - \left(c_{\mathrm{CMP}\to\mathrm{GMP}} + c_{\mathrm{CMP}\to\mathrm{MEP}} + k_{\mathrm{CMP}}\right) N_{\mathrm{CMP}} + c^{\mathrm{Total}}_{\mathrm{MPP}} N_{\mathrm{MPP}} f^{\mathrm{MPP}}_{\mathrm{Bias,\ 1}} \tag{S49}$$

Here, $f^{\mathrm{MPP}}_{\mathrm{Bias,\ 1}}$ is the fraction of MPPs that are myeloid-biased. $c^{\mathrm{Total}}_{\mathrm{MPP}} = c_{\mathrm{MPP}\to\mathrm{CMP}} + c_{\mathrm{MPP}\to\mathrm{CLP}}$ is the total MPP differentiation rate. Assuming that there is no inherent myeloid bias at the ST-HSC stage, *i.e.*, there is no influx of myeloid-biased MPPs via differentiation from the ST-HSC pool, we can modify Eq. S36 to write:

$$\frac{df^{\mathrm{MPP}}_{\mathrm{Bias,\ 1}}}{dt} = r_{\mathrm{MPP}} \cdot f^{\mathrm{MPP}}_{\mathrm{Bias,\ 1}} \cdot p^{\mathrm{Bias}}_{\mathrm{division}} - \left(c^{\mathrm{Total}}_{\mathrm{MPP}} + k_{\mathrm{MPP}}\right) f^{\mathrm{MPP}}_{\mathrm{Bias,\ 1}} + g^{\mathrm{MPP}}_{\mathrm{Bias}} \left(1 - f^{\mathrm{MPP}}_{\mathrm{Bias,\ 1}}\right) \tag{S50}$$

Here, $p^{\mathrm{Bias}}_{\mathrm{division}}$ is the probability with which the myeloid bias in MPPs is passed on to the daughter cells. $g^{\mathrm{MPP}}_{\mathrm{Bias}}$ is the rate of myeloid bias induction in MPPs.

Note that MPP states 0 and 1 may be interpreted as corresponding to distinct epigenetic states of MPPs with epigenetic state 1 associated with higher likelihood of differentiation into CMPs. Alternately, states 0 and 1 can represent distinct MPP subtypes. For example, MPP subtypes MPP2 (defined as Flk2$^-$ CD150$^+$ CD48$^+$ LSK) and MPP3 (defined as Flk2$^-$ CD150$^-$ CD48$^+$ LSK) have been shown to exhibit myeloid bias while the MPP4 subtype (defined as Flk2$^+$ CD150$^-$ CD48$^+$ LSK) is primed to differentiate along the lymphoid trajectory [99]. At steady state, we have:

$$f^{\mathrm{MPP,\ SS}}_{\mathrm{Bias,\ 1}} = \frac{g^{\mathrm{MPP}}_{\mathrm{Bias}}}{\left(c^{\mathrm{Total}}_{\mathrm{MPP}} + k_{\mathrm{MPP}} + g^{\mathrm{MPP}}_{\mathrm{Bias}} - r_{\mathrm{MPP}} \cdot p^{\mathrm{Bias}}_{\mathrm{division}}\right)} \equiv \frac{g^{\mathrm{MPP}}_{\mathrm{Bias}}}{k''_d} \tag{S51}$$

If at $t = t_0$, $f^{\mathrm{MPP}}_{\mathrm{Bias,\ 1}} = f^{\mathrm{MPP,\ 0}}_{\mathrm{Bias,\ 1}}$, then for $t > t_0$, we have:

$$f^{\mathrm{MPP}}_{\mathrm{Bias,\ 1}}(t) = \left(\frac{g^{\mathrm{MPP}}_{\mathrm{Bias}}}{k''_d}\right) + \left(f^{\mathrm{MPP,\ 0}}_{\mathrm{Bias,\ 1}} - \left(\frac{g^{\mathrm{MPP}}_{\mathrm{Bias}}}{k''_d}\right)\right) e^{-k''_d (t - t_0)} \tag{S52}$$

We used Eq. S27-S29, Eq. S49, Eq. S31, Eq. S33, and Eq. S50 to model hematopoietic dynamics with myeloid bias at the MPP stage. Next, we assume that all monocytes secrete the same level of a myeloid bias-inducing cytokine such as IL-1$\beta$. The effect of IL-1$\beta$ on myeloid bias in MPPs is modeled using a Hill function:

$$g^{\mathrm{MPP}}_{\mathrm{Bias}} = g^{\mathrm{MPP,\ 0}}_{\mathrm{Bias}} \mathscr{H}\left(C_{\mathrm{IL}\text{-}1\beta} \cdot N_{\mathrm{Monocyte}}, \lambda^{\mathrm{Sig}}_{\mathrm{Bias}}, \Theta^{\mathrm{Sig}}_{\mathrm{Bias}}, n^{\mathrm{Sig}}_{\mathrm{Bias}}\right) \tag{S53}$$

Here, $C_{\mathrm{IL}\text{-}1\beta}$ is the IL-1$\beta$ expression per monocyte. The strength of the effect of IL-1$\beta$ on myeloid bias is determined by the model parameter $\lambda^{\mathrm{Sig}}_{\mathrm{Bias}}$ which gives the maximum fold-change in the rate of induction of myeloid bias in MPPs that IL-1$\beta$ can cause.

*Simulating myeloid bias induction during an acute inflammatory challenge*— Under homeostatic conditions $H$, the rate of induction of myeloid bias in MPPs is $g^{\mathrm{MPP,\ 0,\ H}}_{\mathrm{Epi}}$. To simulate an acute inflammatory challenge that induces myeloid bias in MPPs between the time points $t^{\mathrm{I}}_{\mathrm{start}}$ and $t^{\mathrm{I}}_{\mathrm{end}}$, we include a challenge-driven increase in the rate of myeloid bias induction:

$$g^{\mathrm{MPP,\ 0}}_{\mathrm{Bias}} = \begin{cases} g^{\mathrm{MPP,\ 0,\ H}}_{\mathrm{Bias}} \cdot \lambda^{\mathrm{I}}_{\mathrm{Bias}} & \text{for} \quad t^{\mathrm{I}}_{\mathrm{start}} \le t \le t^{\mathrm{I}}_{\mathrm{end}} \\ g^{\mathrm{MPP,\ 0,\ H}}_{\mathrm{Bias}} & \text{otherwise} \end{cases} \tag{S54}$$

Here, $\lambda^{\mathrm{I}}_{\mathrm{Bias}} > 1$ is the fold-change in the rate of myeloid bias induction in MPPs caused by the acute inflammatory challenge.

| Parameter | Value | Parameter | Value |
|---|---|---|---|
| $r_{\text{ST-HSC}}$ | 0.042 day$^{-1}$ | $r_{\text{MPP}}$ | 4.0 day$^{-1}$ |
| $r_{\text{CMP}}$ | 4.0 day$^{-1}$ | $r_{\text{GMP}}$ | 4.0* day$^{-1}$ |
| $c_{\text{HSC}\to\text{ST-HSC}}$ | 0.009 day$^{-1}$ | $c_{\text{ST-HSC}\to\text{MPP}}$ | 0.045 day$^{-1}$ |
| $c_{\text{MPP}\to\text{CMP}}$ | 3.992 day$^{-1}$ | $c_{\text{MPP}\to\text{CLP}}$ | 0.022 day$^{-1}$ |
| $c_{\text{CMP}\to\text{GMP}}$ | 2.0 day$^{-1}$ | $c_{\text{CMP}\to\text{MEP}}$ | 3.0 day$^{-1}$ |
| $c_{\text{GMP}\to\text{Monocyte}}$ | 2.7* day$^{-1}$ | $c_{\text{GMP}\to\text{Granulocyte}}$ | 2.7* day$^{-1}$ |

TABLE S4. Values of self-renewal and differentiation rates from Busch *et al.* [98] used to model hematopoietic dynamics in the present study. The values indicated by an asterisk were estimated from the data in Busch *et al.* under certain assumptions (see Sec. III C 1)

.

#### C. Model parameters and analysis of parameter-phenotype mapping

##### 1. Modeling hematopoietic dynamics

We obtained the values of the various self-renewal and differentiation rates from Busch *et al.* [98] who estimated these values for unpertubed hematopoiesis in mice. Note that Busch *et al.* report net proliferation rates, defined as the self-renewal rate minus the death rate. Noting that the death rate parameter always occurs in sum with differentiation rates in Eq. S40-S44, we can set the death rates for HSCs, ST-HSCs, MPPs, CMPs, and GMPs to 0. In doing so, we assume that the death rate is much smaller than the differentiation rate for each HSPC subtype. We can then use the net proliferation rate values reported by Busch *et al.* as the values of the self-renewal rate parameters. Values of the various self-renewal and differentiation rates used for our analysis are listed in Table S4. The values of $r_{\text{GMP}}$, $c_{\text{GMP}\to\text{Monocyte}}$, and $c_{\text{GMP}\to\text{Granulocyte}}$ are not available from [98] and were estimated based on the ratio of GMPs and CMPs in the bone marrow under homeostasis. At steady state, Eq. S31 gives us:

$$\frac{N^{\text{SS}}_{\text{GMP}}}{N^{\text{SS}}_{\text{CMP}}} = \frac{c_{\text{CMP}\to\text{GMP}}}{c_{\text{GMP}\to\text{Monocyte}} + c_{\text{GMP}\to\text{Granulocyte}} - r_{\text{GMP}}} \tag{S55}$$

From Extended Data Figure 6a in [98], we have $\frac{N^{\text{SS}}_{\text{GMP}}}{N^{\text{SS}}_{\text{CMP}}} = 1.4$. Then, assuming that GMPs have the same self-renewal rate as CMPs, *i.e.*, $r_{\text{GMP}} = 4.0$ day$^{-1}$, and that GMPs generate monocytes and granulocytes at equal rates, we set $c_{\text{GMP}\to\text{Monocyte}} = c_{\text{GMP}\to\text{Granulocyte}} = 2.7$ day$^{-1}$. Unless stated otherwise, simulations in this manuscript that involve dynamics of cells in the hematopoietic hierarchy use the self-renewal and differentiation rates listed in Table S4.

##### 2. Modeling epigenetic state dynamics during hematopoiesis

In Fig. 2D-G and Fig. S3D, our focus is on how the dynamics of the epigenetic memory state in HSPCs are affected by the parameters $p_{\text{division}}$ and $p_{\text{differentiation}}$ in the absence of any feedback from monocyte-secreted cytokines. Thus, we set $\lambda^{\text{Epi}}_{\text{Sig}} = \lambda^{\text{Sig}}_{\text{Epi}} = 1$, making the parameters $\Theta^{\text{Sig}}_{\text{Epi}}$ and $n^{\text{Sig}}_{\text{Epi}}$ irrelevant. We further set $g^{\text{i, 0, H}}_{\text{Epi}} = 0$ for all cell types $i$, *i.e.*, we assume that there is no epigenetic reprogramming of HSPCs in the absence of an inflammatory challenge. Thus, under homeostatic conditions, none of the cells will be in an epigenetic memory state.

- Fig. 2D: This figure shows the analytically calculated half-life of the fraction in an epigenetic memory state for different HSPC subtypes. To calculate the half-life for a given cell type, we assume that none of the cells upstream of it in the hematopoietic hierarchy are in an epigenetic memory state, *e.g.*, to calculate the half-life for ST-HSCs, we set $f^{\text{HSC}}_{\text{Epi, 1}} = 0$ at all time points, to calculate the half-life for MPPs, we set $f^{\text{HSC}}_{\text{Epi, 1}} = f^{\text{ST-HSC}}_{\text{Epi, 1}} = 0$, and so on. Then, from Eq. S40-S44, we get:

$$t^{\text{HSC}}_{\text{Epi, 1/2}} = \log 2 \Big/ \left( \left( c_{\text{HSC}\to\text{ST-HSC}} + k_{\text{HSC}} \right) \left( 1 - p_{\text{division}} \right) \right) \tag{S56}$$

$$t^{\text{ST-HSC}}_{\text{Epi, 1/2}} = \log 2 \Big/ \left( \left( c_{\text{ST-HSC}\to\text{MPP}} + k_{\text{ST-HSC}} \right) \left( 1 - p_{\text{division}} \right) + c_{\text{HSC}\to\text{ST-HSC}} \left( \frac{N^{\text{SS}}_{\text{HSC}}}{N^{\text{SS}}_{\text{ST-HSC}}} \right) p_{\text{division}} \right) \tag{S57}$$

$$t^{\text{MPP}}_{\text{Epi, 1/2}} = \log 2 \Bigg/ \Bigg( \left(c_{\text{MPP}\to\text{CMP}} + c_{\text{MPP}\to\text{CLP}} + k_{\text{MPP}}\right)\left(1 - p_{\text{division}}\right) + c_{\text{ST-HSC}\to\text{MPP}} \left(\frac{N^{\text{SS}}_{\text{ST-HSC}}}{N^{\text{SS}}_{\text{MPP}}}\right) p_{\text{division}} \Bigg) \tag{S58}$$

$$t^{\text{CMP}}_{\text{Epi, 1/2}} = \log 2 \Bigg/ \Bigg( \left(c_{\text{CMP}\to\text{GMP}} + c_{\text{CMP}\to\text{MEP}} + k_{\text{CMP}}\right)\left(1 - p_{\text{division}}\right) + c_{\text{MPP}\to\text{CMP}} \left(\frac{N^{\text{SS}}_{\text{MPP}}}{N^{\text{SS}}_{\text{CMP}}}\right) p_{\text{division}} \Bigg) \tag{S59}$$

$$t^{\text{GMP}}_{\text{Epi, 1/2}} = \log 2 \Bigg/ \Bigg( \left(c_{\text{GMP}\to\text{Monocyte}} + c_{\text{GMP}\to\text{Granulocyte}} + k_{\text{GMP}}\right)\left(1 - p_{\text{division}}\right) + c_{\text{CMP}\to\text{GMP}} \left(\frac{N^{\text{SS}}_{\text{CMP}}}{N^{\text{SS}}_{\text{GMP}}}\right) p_{\text{division}} \Bigg) \tag{S60}$$

Here, $t^{\text{i}}_{\text{Epi, 1/2}}$ is the time it takes for the fraction of cells of type $i$ that are in an epigenetic memory state to decrease by half. These values, as a function of $p_{\text{division}}$, are shown by the solid curves in Fig. 2D. For each cell type, the dashed line shows the half-life of the overall population which is given by $\log 2$ divided by the sum of the differentiation and death rates for the cell type. The population half-life is thus independent of $p_{\text{division}}$.

- Fig. 2E: We set $p_{\text{division}} = p_{\text{differentiation}} = 1$. At $t = 0$, the epigenetic state of all cells of a given subtype (or of a chosen set of subtypes) as indicated by the color is set to 1 (*i.e.*, epigenetic memory state). The fraction of monocytes in epigenetic state 1 is then shown as a function of time for each case.

- Fig. 2F: We set $p_{\text{division}} = 1$. At $t = 0$, the epigenetic state of all the cells of the subtype indicated by the subplot heading is set to 1 (*i.e.*, epigenetic memory state). The fraction of monocytes in epigenetic state 1 is then shown as a function of time for different values of $p_{\text{differentiation}}$.

- Fig. 2G: At $t = 0$, the epigenetic state of all HSCs, ST-HSCs, and MPPs is set to 1 (*i.e.*, epigenetic memory state). The plot shows the duration of trained immunity, defined as the length of the time period for $t > 0$ during which more than 10% of the generated monocytes are in epigenetic state 1 (*i.e.*, in an epigenetic memory state).

- Fig. S2D: Same as Fig. 2D; epigenetic state of all cells of the subtype indicated by the subplot heading is set to 1 at $t = 0$ and the trained immunity durability is shown.

To model the effect of feedback from a monocyte-secreted cytokine to the epigenetic reprogramming of HSPCs (Fig. 3E-H and Fig. S4), we must have a non-zero rate of epigenetic reprogramming at baseline (*i.e.*, in the absence of inflammatory challenges) since we consider a multiplicative effect of monocyte-secreted cytokines on $g^{\text{i, 0}}_{\text{Epi}}$ values (see Eq. S47). We set the value of $g^{\text{i, 0, H}}_{\text{Epi}}$ for cell type $i$ by noting that the value of this parameter will determine the fraction of cells of that type that are in an epigenetic memory state in the absence of feedback and when no inflammatory challenge is present. This fraction is expected to be low. Since we are interested in the effect of feedback when the fidelity of epigenetic memory state transmission is low, we first set $p_{\text{division}} = p_{\text{differentiation}} = 0.5$. Then, solving Eq. S40-S45 at steady state, we set $g^{\text{HSC, 0, H}}_{\text{Epi}}$, $g^{\text{ST-HSC, 0, H}}_{\text{Epi}}$, $g^{\text{MPP, 0, H}}_{\text{Epi}}$, $g^{\text{CMP,0, H}}_{\text{Epi}}$, $g^{\text{GMP, 0, H}}_{\text{Epi}}$, and $g^{\text{Monocyte, 0, H}}_{\text{Epi}}$ such that $f^{\text{HSC, SS}}_{\text{Epi, 1}} = f^{\text{ST-HSC, SS}}_{\text{Epi, 1}} = f^{\text{MPP, SS}}_{\text{Epi, 1}} = f^{\text{CMP, SS}}_{\text{Epi, 1}} = f^{\text{GMP, SS}}_{\text{Epi, 1}} = f^{\text{Monocyte, SS}}_{\text{Epi, 1}} = 0.05$. Hereafter, we will refer to this value for cell type $i$ as $g^{\text{i}}_0$. Finally, without any loss of generality, we can set $C_{\text{IL-1}\beta} = 1$.

Now, the only remaining parameters are those that determine the nature of feedback, namely, $\lambda^{\text{Epi}}_{\text{Sig}}$, $\lambda^{\text{Sig}}_{\text{Epi}}$, $\Theta^{\text{Sig}}_{\text{Epi}}$, and $n^{\text{Sig}}_{\text{Epi}}$. Fig. 3E-G and Fig. S4A-G show the behavior for a parameter set for which the model behavior is representative of the dynamics in the presence of a positive feedback loop. In all these panels, we only include feedback from monocyte-secreted IL-1$\beta$ to the epigenetic reprogramming of MPPs, CMPs, and GMPs; there is no feedback to the epigenetic state of HSCs, ST-HSCs, or monocytes. We set $g^{\text{MPP, 0, H}}_{\text{Epi}} = G \cdot g^{\text{MPP}}_0$, $g^{\text{CMP, 0, H}}_{\text{Epi}} = G \cdot g^{\text{CMP}}_0$, and $g^{\text{GMP, 0, H}}_{\text{Epi}} = G \cdot g^{\text{GMP}}_0$; for any other cell type $i$, we set $g^{\text{i}}_{\text{Epi}} = g^{\text{i}}_0$. Finally, we set $\lambda^{\text{Epi}}_{\text{Sig}} = 12$, $\Theta^{\text{Sig}}_{\text{Epi}} = 4.0 \times 10^7$ and $n^{\text{Sig}}_{\text{Epi}} = 6$. Other details regarding the figures are mentioned below:

- Fig. 3E: We simulated the dynamics under homeostatic conditions (*i.e.*, in the absence of inflammatory challenges), varying $G$ (X axis) and $\lambda^{\text{Sig}}_{\text{Epi}}$ (Y axis). For each combination of $G$ and $\lambda^{\text{Sig}}_{\text{Epi}}$ values, we simulated the dynamics starting from two initial conditions, one with $f_{\text{Epi, 1}} = 0$ for all cell types and one with $f_{\text{Epi, 1}} = 1$

for all types. The steady states obtained for the two initial conditions were considered distinct if the difference between $f_{\text{Epi, 1}}^{\text{Monocyte, SS}}$ in the two cases was greater than 0.01.

- Fig. 3F: We simulated the dynamics under homeostatic conditions, varying $G$ (X axis). Regime 1 here corresponds to $\lambda_{\text{Epi}}^{\text{Sig}} = 9$ while regime 4 corresponds to $\lambda_{\text{Epi}}^{\text{Sig}} = 1$.
- Fig. 3G: We simulated an acute inflammatory challenge as per Eq. S48 from $t = 0$ to $t = 21$ days with $\lambda_{\text{Epi}}^{\text{I}} = 10$. Setting $G = 1$, we varied $\lambda_{\text{Epi}}^{\text{Sig}}$.
- Fig. S4A-D: Same as Fig. 3E.
- Fig. S4E: Same as Fig. 3F. Regime 1: $\lambda_{\text{Epi}}^{\text{Sig}} = 9$; regime 2: $\lambda_{\text{Epi}}^{\text{Sig}} = 15$; regime 3: $\lambda_{\text{Epi}}^{\text{Sig}} = 22$; regime 4: $\lambda_{\text{Epi}}^{\text{Sig}} = 1$
- Fig. S4F: Same as Fig. 3G.
- Fig. S4G: We simulated acute inflammation as per Eq. S48 from $t = 0$ to $t = 21$ days with $\lambda_{\text{Epi}}^{\text{I}} = 10$. We varied both $G$ (X axis) and $\lambda_{\text{Epi}}^{\text{Sig}}$ (Y axis) and plotted the memory durability, defined as the duration for $t > 21$ for which $f_{\text{Epi, 1}}^{\text{Monocyte}}$ is at least 10% higher than at $t = 0$.

*Parameter-phenotype mapping analysis*— We simulated an acute inflammatory challenge as per Eq. S48 from $t = 0$ to $t = 21$ days with $\lambda_{\text{Epi}}^{\text{I}} = 10$ for an ensemble of parameter sets sampled from within the biological range. Once again, we consider the scenario wherein the inflammatory challenge only increases the epigenetic reprogramming rate of MPPs, CMPs, and GMPs, and we include feedback from monocyte-secreted IL-1$\beta$ to these subtypes. Randomly sampled parameters included $g_{\text{Epi}}^{\text{i, 0, H}}$ for each cell type $i$, $p_{\text{division}}$, $p_{\text{differentiation}}$, $\lambda_{\text{Sig}}^{\text{Epi}}$, $\lambda_{\text{Epi}}^{\text{Sig}}$, $\Theta_{\text{Epi}}^{\text{Sig}}$, and $n_{\text{Epi}}^{\text{Sig}}$. The ranges from which these parameters were sampled are shown in Table S5. $g_{\text{Epi}}^{\text{i, 0, H}}$ values were sampled as follows. We first sampled, for each cell type, the fraction of cells that are epigenetically reprogrammed in the absence of cytokine signaling and when no inflammatory challenge is present; these fractions were drawn uniformly between 0.05 and 0.2. We then used these values to calculate $g_{\text{Epi}}^{\text{i, 0, H}}$ for cell type $i$ by solving Eq. S40-S45. Thus, in the absence of inflammatory challenges and cytokine signaling, the fraction of cells of any subtype in an epigenetic memory state is low: between 5% and 20% (see Table S5). To ensure that this fraction remains low in the presence of feedback from monocyte-secreted IL-1$\beta$, $\Theta_{\text{Epi}}^{\text{Sig}}$ was chosen to be 2-3 times the IL-1$\beta$ concentration in the absence of IL-1$\beta$-mediated feedback and any parameter set for which $\mathscr{H}\left(C_{\text{IL-1}\beta}^{\text{No challenge}}, \lambda_{\text{Epi}}^{\text{Sig}}, \Theta_{\text{Epi}}^{\text{Sig}}, n_{\text{Epi}}^{\text{Sig}}\right) > \lambda_{\text{Epi}}^{\text{Sig}}/2$ was discarded. This choice was made to ensure that feedback from IL-1$\beta$ is only weakly active under homeostatic conditions irrespective of the value of $\lambda_{\text{Epi}}^{\text{Sig}}$ and that the epigenetically reprogrammed fraction is low even in the presence of IL-1$\beta$-mediated feedback as long as no inflammatory challenges are present. Finally, we discarded any sampled parameter set for which the fraction of cells in an epigenetic memory state at baseline for any cell type was larger than 20%. Overall, we calculated the trained immunity durability for a total of 45000 parameter sets and used the dataset consisting of parameter sets and the corresponding durability values for the analysis shown in Fig. 3H and Fig. S4H-K:

- Fig. S4H: Statistics of memory durability values shown as a function of $\lambda_{\text{Sig}}^{\text{Epi}}$ and $\lambda_{\text{Epi}}^{\text{Sig}}$. Mean, median, and maximum values were calculated after excluding any parameter sets with the "infinite" memory phenotype.
- Fig. S4I: Statistics of memory durability values shown as a function of $p_{\text{division}}$ and $p_{\text{differentiation}}$. Mean, median, and maximum values were calculated after excluding any parameter sets with the "infinite" memory phenotype.
- Fig. S4K: Distribution of memory durability values for parameter sets with weak feedback ($\lambda_{\text{Sig}}^{\text{Epi}}$, $\lambda_{\text{Epi}}^{\text{Sig}} \leq 6$) and for parameter sets with strong feedback ($\lambda_{\text{Sig}}^{\text{Epi}}$, $\lambda_{\text{Epi}}^{\text{Sig}} > 6$). MATLAB function `ksdensity` was used to obtain a probability density estimate for each histogram; this estimate is shown by the darker curves.
- Fig. 3H (top): The long memory phenotype whose frequency is shown includes memory durability values higher than 180 days; this includes the "infinite" memory phenotype.
- Fig. S4J (top): We first used the memory durability values to define seven phenotypes: durability $\leq 14$, $14 <$ durability $\leq 28$, $28 <$ durability $\leq 42$, $42 <$ durability $\leq 60$, $60 <$ durability $\leq 90$, $90 <$ durability $\leq 180$ and $180 <$ durability. We then used the MATLAB function `fitcensemble` to train random forest models of with different numbers of decision trees to predict the memory durability phenotype for a given parameter set. 10-fold cross-validation performance for different forest sizes is shown in the top panel. The dashed black line indicates a model with 500 trees; this model size was used for the analysis shown in the bottom panel and in Fig. 3H (bottom panels).

| Parameter | Range |
|---|---|
| $p_{\text{division}}$ | 0.1-0.9 |
| $p_{\text{differentiation}}$ | 0.1-0.9 |
| $g_{\text{Epi}}^{\text{i, 0, H}}$ | To keep reprogrammed fraction between 0.05 and 0.2 for each cell type when no cytokine signaling or inflammatory challenge is present |
| $\lambda_{\text{Sig}}^{\text{Epi}}$ | 2-32 |
| $\lambda_{\text{Epi}}^{\text{Sig}}$ | 2-32 |
| $\Theta_{\text{Epi}}^{\text{Sig}}$ | 2-3 times the IL-1$\beta$ level when $\lambda_{\text{Sig}}^{\text{Epi}} = \lambda_{\text{Epi}}^{\text{Sig}} = 1$ |
| $n_{\text{Epi}}^{\text{Sig}}$ | 2-8 (integer values only) |

TABLE S5. Ranges from which different model parameters were sampled for the parameter-phenotype mapping analysis for epigenetic dynamics in the hematopoietic hierarchy.

- Fig. S4J (bottom): Performance of the 500-tree random forest model on the training dataset (80% of the overall dataset) and on the test set (20% of the overall dataset).
- Fig. 3H (bottom): We trained two 500-tree random forest models: one on parameter sets with $p_{\text{differentiation}} \leq 0.5$ and one on parameter sets with $p_{\text{differentiation}} > 0.5$. For both models, we used the MATLAB function `predictorImportance` to calculate the importance of each individual parameter in predicting the memory phenotype. The normalized importance for each parameter (importance divided by the sum of the importance values across all parameters) is shown.

##### 3. *Modeling feedback from monocyte-secreted cytokines to myeloid bias during hematopoiesis*

We use the values of the self-renewal, differentiation, and cell death rate parameters mentioned in Sec. III C 1. Note that the model of feedback to myeloid bias in hematopoiesis includes Eq. S27-S29, Eq. S49, Eq. S31, Eq. S33, and Eq. S50. In Eq. S27, the value of $r_{\text{HSC}}$ and the mathematical form of $f_{\text{Bound}}^{\text{HSC}}$ are unknown (and have not been used for analysis until now). Since we model myeloid versus lymphoid bias at the MPP stage, the dynamics of both HSCs and ST-HSCs remain in steady state throughout, unaffected by IL-1$\beta$ or by the acute inflammatory challenge (Eq. S54). Thus, the values of $r_{\text{HSC}}$ and $f_{\text{Bound}}^{\text{HSC}}$ only control the overall size of the HSC pool, and, consequently, of the ST-HSC and MPP pools, without affecting the nature of the overall dynamics. This allows us to set $r_{\text{HSC}} = 1$ day$^{-1}$ and $f_{\text{Bound}}^{\text{HSC}} = 0.1$ without any loss of generality. Next, we choose $g_{\text{Bias}}^{\text{MPP, 0, H}}$ such that under homeostatic conditions and in the absence of feedback from IL-1$\beta$ to myeloid bias, $f_{\text{Bias, 1}}^{\text{MPP}} = 0.05$; this value of $g_{\text{Bias}}^{\text{MPP, 0, H}}$ is hereafter referred to as $g_{\text{Bias}}^{0}$. Here, we have assumed that only a small fraction of MPPs is likely to exhibit myeloid bias under homeostatic conditions. Finally, without any loss of generality, we can set $C_{\text{IL-1}\beta} = 1$.

Fig. S5A shows the behavior for the no feedback case and we set $\lambda_{\text{Bias}}^{\text{Sig}} = 1$; values of $\Theta_{\text{Bias}}^{\text{Sig}}$ and $n_{\text{Bias}}^{\text{Sig}}$ are irrelevant in such a scenario. We further set $g_{\text{Bias}}^{\text{MPP, 0, H}} = g_{\text{Bias}}^{0}$.

- Fig. S5A (top): We simulated an acute inflammatory challenge as per Eq. S54 from $t = 0$ to $t = 21$ days with $\lambda_{\text{Bias}}^{\text{I}} = 16$. Monocyte count is shown as a function of time for different values of $p_{\text{division}}^{\text{Bias}}$.
- Fig. S5A (bottom): Same as Fig. S5A (top); memory durability, defined as the duration for $t > 21$ for which the monocyte count is at least 10% above the count at $t = 0$.

Fig. S5B-I show the model behavior for values of $\lambda_{\text{Bias}}^{\text{Sig}}$, $\Theta_{\text{Bias}}^{\text{Sig}}$, and $n_{\text{Bias}}^{\text{Sig}}$ for which the model behavior is representative of the dynamics in the presence of a positive feedback loop. For the analysis shown in these panels, we set $\Theta_{\text{Bias}}^{\text{Sig}} = 2.8 \times 10^6$ and $n_{\text{Bias}}^{\text{Sig}} = 4$. Since we are interested in the effect of feedback when the myeloid bias is transmitted with low fidelity during MPP self-renewal, we further set $p_{\text{division}}^{\text{Bias}} = 0.5$.

- Fig. S5B: We simulated the dynamics under homeostatic conditions (*i.e.*, in the absence of inflammatory challenges) and vary $\frac{g_{\text{Bias}}^{\text{MPP, 0, H}}}{g_{\text{Bias}}^{0}}$ (X axis) and $\lambda_{\text{Bias}}^{\text{Sig}}$ (Y axis). For each combination of $\frac{g_{\text{Bias}}^{\text{MPP, 0, H}}}{g_{\text{Bias}}^{0}}$ and $\lambda_{\text{Bias}}^{\text{Sig}}$ values, we simulated the dynamics starting from two distinct initial conditions, one with $f_{\text{Bias, 1}}^{\text{MPP}} = 0$ and one with $f_{\text{Bias, 1}}^{\text{MPP}} = 1$. The steady states obtained in the two cases were considered distinct if difference in monocyte count between them was greater than 100.
- Fig. S5C-D: Same as Fig. S5B.

- Fig. S5E (top): We simulated the dynamics under homeostatic conditions, setting $\lambda^{\text{Sig}}_{\text{Bias}} = 5$ and varying $\frac{g^{\text{MPP, 0, H}}_{\text{Bias}}}{g^{0}_{\text{Bias}}}$ (X axis).

- Fig. S5E (bottom): We simulated the dynamics under homeostatic conditions, setting $\frac{g^{\text{MPP, 0, H}}_{\text{Bias}}}{g^{0}_{\text{Bias}}} = 1$ and varying $\lambda^{\text{Sig}}_{\text{Bias}}$ (X axis).

- Fig. S5F: Same as Fig. S5E (top). Regime 1: $\lambda^{\text{Sig}}_{\text{Bias}} = 5$; regime 2: $\lambda^{\text{Sig}}_{\text{Bias}} = 8$; regime 3: $\lambda^{\text{Sig}}_{\text{Bias}} = 20$; regime 4: $\lambda^{\text{Sig}}_{\text{Bias}} = 2$.

- Fig. S5G-H: We simulated an acute inflammatory challenge as per Eq. S54 from $t = 0$ to $t = 21$ days with $\lambda^{\text{I}}_{\text{Bias}} = 16$. We set $g^{\text{MPP, 0, H}}_{\text{Bias}} = g^{0}_{\text{Bias}}$ and vary $\lambda^{\text{Sig}}_{\text{Bias}}$.

- Fig. S5I: We simulated an acute inflammatory challenge as per Eq. S54 from $t = 0$ to $t = 21$ days with $\lambda^{\text{I}}_{\text{Bias}} = 16$. We vary $\frac{g^{\text{MPP, 0, H}}_{\text{Bias}}}{g^{0}_{\text{Bias}}}$ (X axis) and $\lambda^{\text{Sig}}_{\text{Bias}}$ (Y axis). Memory durability is defined as the duration for $t > 21$ for which the monocyte count is at least 10% higher than the count at $t = 0$.

*Parameter-phenotype mapping analysis*— We simulated an acute inflammatory challenge for an ensemble of parameter sets sampled from within the biological range. Inflammatory challenge was simulated as per Eq. S54 from $t = 0$ to $t = 21$ days with $\lambda^{\text{I}}_{\text{Bias}} = 16$. Sampled model parameters included $p^{\text{Bias}}_{\text{division}}$, $g^{\text{MPP, 0, H}}_{\text{Bias}}$, $\lambda^{\text{Sig}}_{\text{Bias}}$, $\Theta^{\text{Sig}}_{\text{Bias}}$, and $\Theta^{\text{Sig}}_{\text{Bias}}$. The ranges from which these parameters were sampled are shown in Table S6. $g^{\text{MPP, 0, H}}_{\text{Bias}}$ was sampled from a range dependent on the sampled value of $p^{\text{Bias}}_{\text{division}}$: after sampling $p^{\text{Bias}}_{\text{division}}$, we choose $g^{\text{MPP, 0, H}}_{\text{Bias}}$ such that $\left(\frac{f^{\min}_{\text{Bias, 1}}}{1-f^{\min}_{\text{Bias, 1}}}\right) k''_d < g^{\text{MPP, 0, H}}_{\text{Bias}} < \left(\frac{f^{\max}_{\text{Bias, 1}}}{1-f^{\max}_{\text{Bias, 1}}}\right) k''_d$ with $k''_d = c^{\text{Total}}_{\text{MPP}} + k_{\text{MPP}} - r_{\text{MPP}} \cdot p^{\text{Bias}}_{\text{division}}$ (see Eq. S51). We set $f^{\min}_{\text{Bias, 1}} = 0.05$ and $f^{\max}_{\text{Bias, 1}} = 0.2$ as the minimum and maximum possible values, respectively, of $f^{\text{MPP, SS}}_{\text{Bias, 1}}$ under homeostatic conditions and when there is no or very weak feedback from IL-1$\beta$ to MPPs. Alternately, one can sample $g^{\text{MPP, 0, H}}_{\text{Bias}}$ independent of $p^{\text{Bias}}_{\text{division}}$ and then discard parameter sets for which $f^{\text{MPP, SS}}_{\text{Bias, 1}}$ under homeostatic conditions is too high (*i.e.*, greater than $f^{\max}_{\text{Bias, 1}}$). $\Theta^{\text{Sig}}_{\text{Bias}}$ was chosen to be 2-3 times the concentration of IL-1$\beta$ under homeostasis and in the absence of feedback (*i.e.*, when $\lambda^{\text{Sig}}_{\text{Bias}} = 1$). This ensures that the feedback from IL-1$\beta$ to MPPs is only weakly active under homeostatic conditions and that $f^{\text{MPP, SS}}_{\text{Bias, 1}}$ under homeostasis remains slow. Finally, any parameter sets for which $f^{\text{MPP, SS}}_{\text{Bias, 1}}$ under homeostasis exceeded $f^{\max}_{\text{Bias, 1}}$ was discarded. Overall, we sampled a total of 45000 parameter sets and calculated the memory durability in each case. This dataset formed the basis of the analysis shown in Fig. S5J-L:

- Fig. S5J (bottom): Memory durability values for the parameter sets in the ensemble with finite durability values, shown as a function of $\lambda^{\text{Sig}}_{\text{Bias}}$.

- Fig. S5J (top): Frequency of the "infinite" memory phenotype among the parameter sets in the ensemble, shown as a function of $\lambda^{\text{Sig}}_{\text{Bias}}$.

- Fig. S5K (top): We defined four memory durability phenotypes based on the durability values: durability $\leq 90$, $90 <$ durability $\leq 180$, $180 <$ durability $\leq 360$, and $360 <$ durability; the last phenotype includes the "infinite" memory case. We then trained random forest models with different numbers of decision trees to predict the memory phenotype for a given parameter set; MATLAB function `fitcensemble` was used for this purpose. Inputs to these random forest models were the parameters $p^{\text{Bias}}_{\text{division}}$, $f^{\text{MPP, SS}}_{\text{Bias, 1}}$ (fraction of myeloid-biased MPPs when no feedback or inflammatory challenge is present; used as an input in place of $g^{\text{MPP, 0, H}}_{\text{Bias}}$), $\lambda^{\text{Sig}}_{\text{Bias}}$, $\Theta^{\text{Sig}}_{\text{Bias}}$, and $n^{\text{Sig}}_{\text{Bias}}$. 10-fold cross-validation performance of the models with different tree counts is shown in Fig. S5 (top). The black dashed line corresponds to a 600-tree model which was used for the analysis shown in Fig. S5K (bottom) and Fig. S5L (top).

- Fig. S5K (bottom): Performance of the 600-tree random forest model on the training set (80% of the overall dataset) and the test set (20% of the overall dataset).

- Fig. S5L (top): We used the MATLAB function `predictorImportance` to calculate the importance of each individual parameter in predicting the memory phenotype in the case of the 600-tree model. The normalized importance for each parameter (importance divided by the sum of the importance values across all parameters) is shown.

| Parameter | Range |
|---|---|
| $p^{\text{Bias}}_{\text{division}}$ | 0.1–0.9 |
| $g^{0,\,\text{H}}_{\text{Epi}}$ | Between $\left(\frac{f^{\min}_{\text{Bias},\,1}}{1-f^{\min}_{\text{Bias},\,1}}\right) k_d''$ and $\left(\frac{f^{\max}_{\text{Bias},\,1}}{1-f^{\max}_{\text{Bias},\,1}}\right) k_d''$ <br> $k_d'' = c^{\text{Total}}_{\text{MPP}} + k_{\text{MPP}} - r_{\text{MPP}} \cdot p^{\text{Bias}}_{\text{division}}$ <br> $f^{\min}_{\text{Bias},\,1} = 0.05$ and $f^{\max}_{\text{Bias},\,1} = 0.2$ |
| $\lambda^{\text{Sig}}_{\text{Bias}}$ | 2–12 |
| $\Theta^{\text{Sig}}_{\text{Bias}}$ | 2–3 times the IL-1$\beta$ level when $\lambda^{\text{Sig}}_{\text{Bias}} = 1$ |
| $n^{\text{Sig}}_{\text{Epi}}$ | 2–8 (integer values only) |

TABLE S6. Ranges from which different model parameters were sampled for the parameter-phenotype mapping in hematopoietic dynamics with myeloid bias in MPPs.

- Fig. S5L (bottom): Distribution of the memory durability values for the parameter sets in the ensemble with weak feedback ($\lambda^{\text{Sig}}_{\text{Bias}} \leq 4$) and those with strong feedback ($\lambda^{\text{Sig}}_{\text{Bias}} > 4$). MATLAB function `ksdensity` was used to obtain a probability density estimate for each histogram; this estimate is shown by the darker curves.

##### D. Assumptions, limitations, and other comments

- Our model of hematopoietic dynamics involves some of the same assumptions that were made in the case of $\text{T}_{\text{M}}$ cell dynamics (Sec. I E) and in the case of tissue macrophage dynamics (Sec. II D):
  - We model the bone marrow niche as a well-mixed "bag" of cells, disregarding the architecture of the niche (reviewed in [100]) and the spatial organization of hematopoietic and non-hematopoietic cell types in the bone marrow. Our model also does not incorporate the population or epigenetic state dynamics of the many non-hematopoietic cell types in the bone marrow.
  - Our model does not track the dynamics, epigenetic state, or myeloid bias of individual cells. Only the size of each cell type pool, the fraction of each cell type pool that is in an epigenetic memory state, and in the case of MPPs, the fraction that is myeloid-biased, is monitored in our modeling setup.
  - Similar to the model of epigenetic state dynamics in tissue macrophages, we once again use a single binary variable to describe the epigenetic state of different cell types, thereby disregarding any heterogeneity in the behavior of cells categorized as epigenetically reprogrammed.
- We have modeled the overall hematopoietic dynamics in an organism instead of modeling hematopoiesis in different bone marrow compartments separately. Thus, our model cannot account for any heterogeneity in hematopoiesis in the marrow from different bones. This choice was largely made to simplify the modeling task, and is motivated by the fact that while the marrow in different bones differs in morphology and tissue composition, HSPCs from different bone marrow compartments have been shown to exhibit similar gene expression, phenotype, and function [101]. By writing a separate set of differential equations (Eq. S27-S31, S34-S38) for each bone marrow compartment, the present model can easily be extended to account for differences in the hematopoietic process at different sites.
- Note that, typically, effector immune cells respond to inflammatory challenges and secrete cytokines in the periphery, away from the bone marrow niche. Cytokines secreted in the periphery can enter the circulation and ultimately reach the highly vascular bone marrow niche. Thus, the delay between effector immune cell generation in the bone marrow and cells reaching the periphery, kinetics of cytokine transport via circulation, and the half-life of different cytokines are all likely to affect the hematopoietic dynamics responding to feedback from effector immune cell signaling. Given that there is extensive evidence of inflammation in the periphery affecting hematopoiesis, we can safely assume that all such parameters are in a regime that permits cytokine-mediated cross-talk between effector immune cells in the periphery and HSPCs in the bone marrow. Based on this assumption, our model does not explicitly incorporate the kinetics of effector immune cell trafficking and cytokine transport. We model HSPCs and effector immune cells as parts of the same well-mixed compartment.
- As mentioned earlier in Sec. III A, our mathematical description is based on the classical hierarchical model which describes hematopoiesis as a cascade of differentiation events [75, 102]. This classical picture has been revised with the advent of single cell "-omics" technologies that have revealed heterogeneity in HSCs, MPPs,

and other HSPC subsets. For example, the intermediate-term HSC (IT-HSC) subset was defined as having self-renewal capability that is between that of HSCs and ST-HSCs [103]. The MPP subset can be further subdivided into MPP1, MPP2, MPP3, and MPP4, with the MPP4 subset exhibiting lymphoid bias [99]. We chose the classical picture to model hematopoietic dynamics since this allowed us to use the various kinetic parameters estimated by Busch *et al.* [98] who used a flow cytometry panel that allows for the resolution of HSPCs into classical subsets. In any case, our focus here is on the epigenetic state dynamics in HSPCs and on the effect of feedback from effector cell-secreted cytokines on the overall hematopoietic process. Both these phenotypes are unlikely to be affected by the resolution of HSPC subtyping in the mathematical model. The present modeling framework may easily be extended to incorporate specific HSPC subsets of interest.

- To explore the effect of feedback from effector immune cell-secreted cytokines on epigenetic state dynamics during the hematopoietic process, we have used the example of feedback from monocyte-secreted IL-1$\beta$ to the epigenetic reprogramming of MPPs, CMPs, and GMPs. We have additionally modeled the effect of feedback from monocyte-secreted IL-1$\beta$ to myeloid bias in MPPs. These signaling pathways were chosen as representative examples to explore how feedback from effector immune cells can affect trained immunity, and by no means are the only examples of feedback to the hematopoietic process. Once again, the present framework can easily be adapted to model the effect of feedback mediated by a specific cytokine of interest.

#### CODE AND DATA AVAILABILITY

Matlab code used to simulate the various mathematical models and to carry out the various analyses is available on Github at `https://github.com/TsangLabCSEI/antigen-agnostic-memory`. The repository includes instructions on how to run the code as well as the parameter-phenotype mapping datasets used for MAPPA analysis.

---

[10] C. Liu, A. J. Martins, W. W. Lau, N. Rachmaninoff, J. Chen, L. Imberti, D. Mostaghimi, D. L. Fink, P. D. Burbelo, K. Dobbs, O. M. Delmonte, N. Bansal, L. Failla, A. Sottini, E. Quiros-Roldan, K. L. Han, B. A. Sellers, F. Cheung, R. Sparks, T.-W. Chun, S. Moir, M. S. Lionakis, M. S. Abers, R. Apps, M. Bosticardo, P. Milanez-Almeida, M. P. Mulè, E. Shaw, Y. Zhang, F. Castelli, M. L. Muiesan, G. Tomasoni, F. Scolari, A. Tucci, C. Rossi, H. C. Su, D. B. Kuhns, J. I. Cohen, L. D. Notarangelo, and J. S. Tsang, Time-resolved systems immunology reveals a late juncture linked to fatal COVID-19, Cell **184**, 1836 (2021).
